# Establishing a standardized genetic toolkit for the radiation-resistant extremophile *Deinococcus radiodurans*

**DOI:** 10.1101/2025.06.09.658681

**Authors:** Trevor R. Simmons, Antonio Cordova, Kobe B. Grismore, Laci C. Moline, Anna C. Stankes, Jennah Johnson, Christian Bök, Lydia M. Contreras

## Abstract

Deinococcus radiodurans is a highly radiation-resistant extremophile with potential for biomanufacturing and bioremediation in harsh environments, including extraterrestrial settings. However, genetic engineering in this organism has been constrained by limited genetic tools. Here, we establish a comprehensive genetic toolkit for D. radiodurans that enables tunable gene regulation with genome-engineering tools. We have standardized a library of 36 constitutive promoter sequences sourced from the native D. radiodurans genome and from synthetic sources, spanning 45-fold range gene expression in the context of plasmids. We have also identified 136 variants of ribosome binding sites (RBS), variants using a high-throughput screen for precise translational control across a 963-fold range of expression when used in our plasmid-based system. Additionally, we have developed a codon-optimizer script that we leverage to improve the function of four fluorescent proteins in D. radiodurans. Next, we characterized 9 small molecule-inducible promoter systems, and identified four key inducible promoter systems that achieve between 3-fold and 12-fold signal amplification, as well as titratability across induction concentrations in D. radiodurans. To engineer the D. radiodurans genome, we present a novel method for gene integrations, compatible with de novo sequences, up to 3kB in length, doing so with 87% efficiency. Lastly, we repurpose the RNA-directed nuclease, TnpB, as a novel post-transcriptional tool for programmable gene repression analogous to CRISPRi-based systems, and this tool can achieve up to 70% repression of its target. Collectively, this toolkit provides modular, standardized components for both plasmid engineering and chromosomal engineering in D. radiodurans to improve its genetic tractability and to facilitate its deployment.

**TOC Image:** 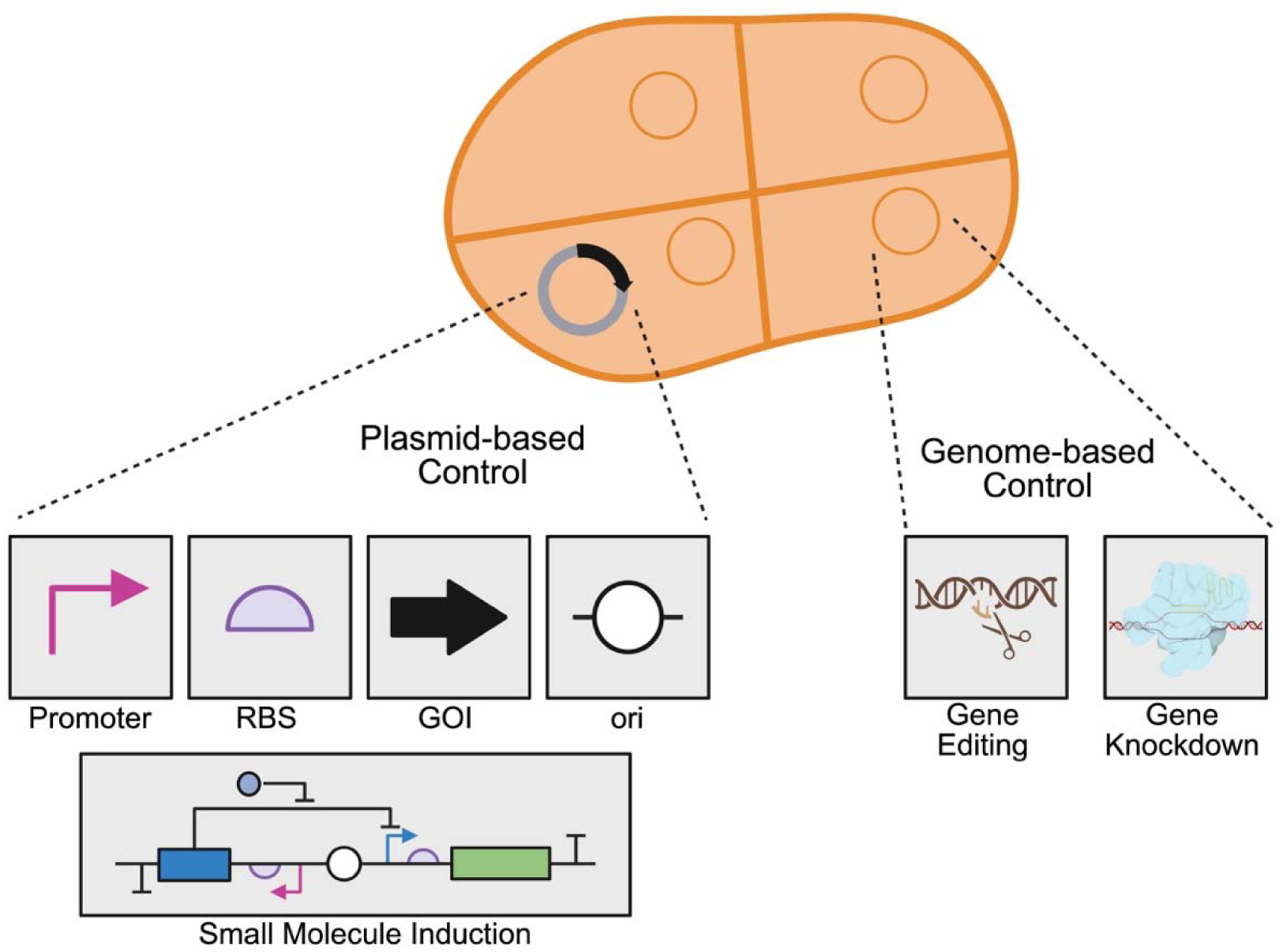

## Introduction

As synthetic biology advances, more microbes become genetically tractable, which opens the door for novel bioprocessing applications. Current model organisms have supported many advances in the field and provide robust chassis for prototyping novel tools. However, these organisms can struggle to perform outside of standard lab conditions in the absence of significant genetic modifications. An underexplored alternative is to deploy genetic tools in microbes that have been natively evolved to survive in harsh or unfavorable environmental conditions unsuitable for standard lab conditions; these include high salinity, harsh pH, extreme desiccation, and elevated radiation^1–4^. Commonly referred to as Next-Generation Industrial Biotechnology (NGIB)^5^, these microbes and associated engineering tools enable new opportunities in both bioprocessing and biomanufacturing. Two examples are *Corynebacteria*^6^ *and Halamonas* species, which as high producers of N-acetylglucosamine and hyaluronic acid and native to sea water, respectively^7,8^, could find high relevance to cosmetic, food pharmaceuticals and agricultural industries. Likewise, use of microorganisms for extraterrestrial applications^9^ is of high interest given the desire to explore Biomanufacturing in space, where the durability of current model organisms (e.g. *Escherichia coli* and *Saccharomyces cerevisiae* adds to the challenges of biomanufacturing logistics, and microbial resource allocation^10^.

An important factor in the development of effective extraterrestrial biomanufacturing tools is the ability for microbes to withstand higher doses of radiation exposure, since most space travel lacks radiation shielding from an atmosphere^11,12^. Most bacteria struggle to survive prolonged radiation exposure, particularly those identified as GRAS or commensal bacteria^13^. One species of interest for extraterrestrial biomanufacturing is *Deinococcus radiodurans*^14^ as it can withstand high levels of ionizing radiation^15^. Additionally, *D. radiodurans* has recently demonstrated the ability to survive for up to three years in space^16^. The ability of *D. radiodurans* to maintain its genome under ionizing radiation and oxidative stress endows it ability to grow in harsh environments, as well as to utilize its metabolism to degrade toxic chemicals^3,17^. For these reasons, several potential applications for both bioprocessing and bioremediation have been demonstrated using *D. radiodurans*; some include degradation of compounds found in nuclear waste, remediation of mercury pollution in soil, and conversion of metallic waste to nanoparticles^2,3,18^. Given the promising utility of this organism, it is crucial to establish a standardized, genetic toolkit in *D. radiodurans* that includes all the critical features required for efficient user-controlled gene regulation and genome engineering, as it has been done for other industrially-relevant microbes^19–26^. The standardization of genetic components not only allows for improved transferability of work, but also allows the field to refine engineering applications in this organism.

Currently, engineering efforts in *D. radiodurans* are currently limited by the number of characterized genetic components and tools available in this organism. Since its discovery and initial characterization in 1956^27^, only one plasmid for heterologous gene expression has been widely established^28^, and only a handful of promoters native to *D. radiodurans* have been studied^29,30^, with only one fully characterized ribosome binding site^29^. Currently, the only small molecule-inducible promoter that has been ported from model organisms into *D. radiodurans* is a LacI-based system developed by Lecointe and colleagues over two decades ago^31^. Since then, few efforts have been made to standardize or identify other small-molecule inducible promoter systems that are functional and titratable in *D. radiodurans*. Beyond plasmid-based gene expression, there has been minimal effort to develop a protocol for efficient genome control, through either editing or regulating genomic expression. Moreover, previous work has demonstrated that Cas-based genome editing is harmful to cellular health in *D. radiodurans,* and this problem has limited the development of tools for use in the organism^32^.

In this work, we establish a genetic toolkit for precise gene regulation by first identifying 2 new origins of replication for plasmid-based expression in addition to the single previously identified origin of replication. We then standardize 36 promoter sequences, sourced from the native *D. radiodurans* genome and from synthetic sequences, all which span a 45-fold range of gene expression. We also design a library of ribosome-binding site (RBS) sequences that provide tunability across almost three orders of magnitude (963x) for precise gene control, both transcriptional and translational. Next, we identify 3 small molecule inducible transcriptional regulator constructs with robust functionality in *D. radiodurans*. Concurrently, we introduce two tools for genome engineering and regulation: (i) an optimized method for genomic integration up to 3kB in length and (ii) a genomic post-transcriptional control system that repurposes the native transposon-associated protein TnpB to allow for targeted repression in the genome of *D. radiodurans*. Overall, with this toolkit, we provide users with both plasmid-based control and genomic-target control, applicable to biological engineering in *D. radiodurans*.

## Results

### Screening putative and synthetic promoter sequences for standardized transcriptional control

Historically, the only plasmid broadly used in *D. radiodurans* is the pRAD1 vector derived from work by Meima et al.,^28^ — a vector that contains the minimal origin of replication (ori) sequence from the SARK cryptic plasmid in *D. radiodurans,* a sequence known as RepU. Meima and colleagues added the ColE1 ori as well as antibiotic resistance cassettes to the sequence to create a shuttle vector between *E. coli* and *D. radiodurans.* This plasmid has served as the base vector for nearly all engineering applications in *D. radiodurans* over the last two decades^15,33–36^. The pRAD1 plasmid also contains the P_GroES_ promoter native to *D. radiodurans* and its ribosome binding site (RBS) sequence. Here, we design a new vector by reducing the size of the original pRAD1 plasmid, removing the ampicillin resistance cassette as well as the truncated P_CAT_ promoter (located upstream of the chloramphenicol resistance cassette). We replace the latter with the full P_CAT_ promoter from the pAcyc184 plasmid. This plasmid design conserves the well characterized promoter and RBS from the original pRAD1 plasmid. To ensure functionality, we inserted a *sfgfp* CDS downstream and we observed constitutive sfGFP fluorescence. We have named this plasmid pAT00.20.001-sfGFP, and this construct serves as our base plasmid for all future constructs in this work (Fig. 1a).

**Figure 1.**
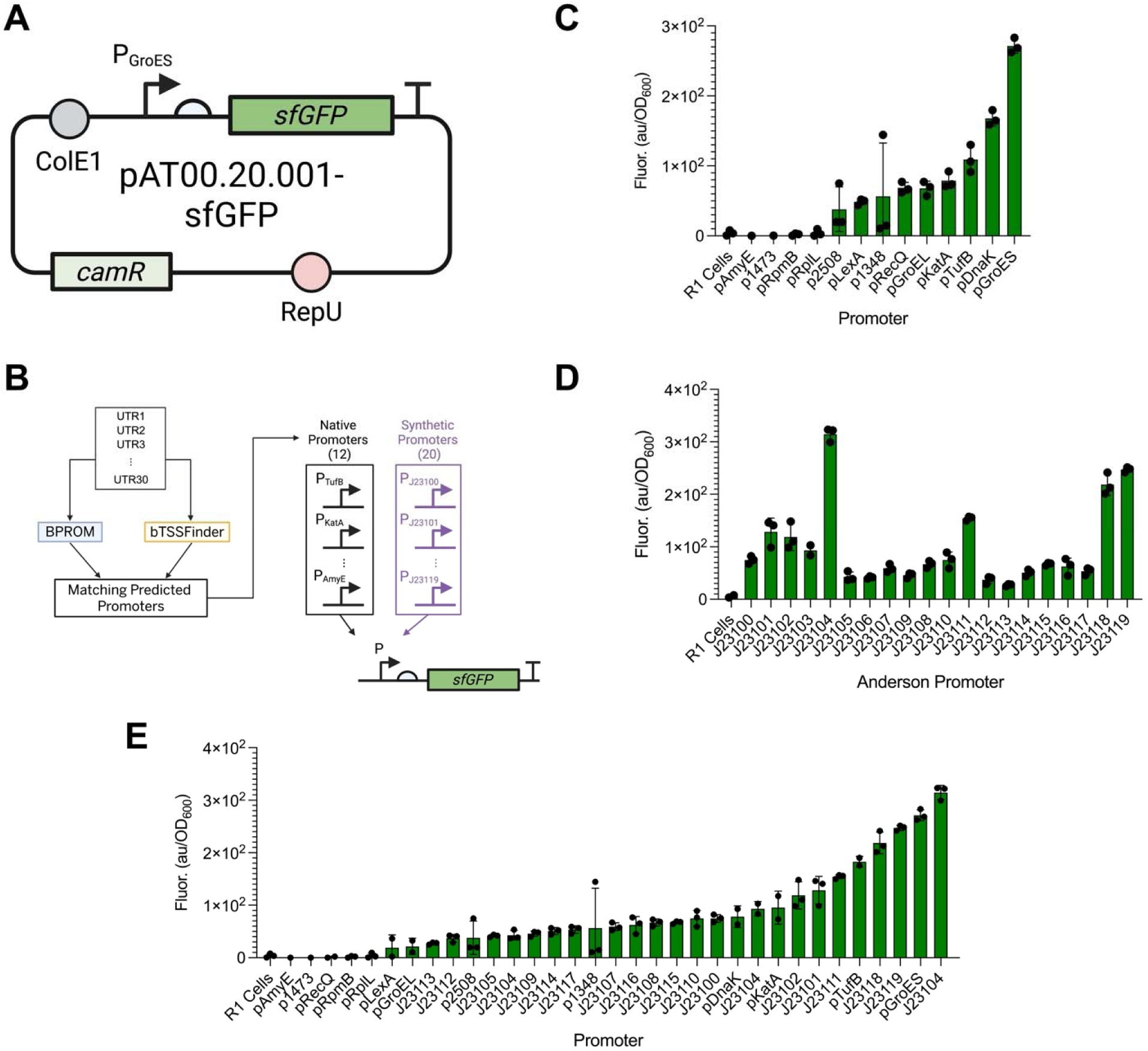
Screening native and synthetic constitutive promoters to establish a standardized set of constitutive promoters in *D. radiodurans.* A) Plasmid map for pAT00.20.001-sfGFP base plasmid. B) Schematic for testing each of the constitutive promoters from the native set as well as the synthetic set. The original P_GroES_ promoter was replaced with each promoter, then fluorescence of sfGFP was measured. C) sfGFP signal after 24 hours of culturing under the control of each native promoter from *D. radiodurans.* D) sfGFP signal after 24 hours of culturing under the control of each synthetic Anderson Promoter. E) Compiled promoter library from the native and synthetic promoter sets from 1C and 1D plotted on the same graph for ease of comparison.

Given that the P_GroES_ promoter is one of the few well-characterized promoters known in *D. radiodurans*, users have been limited in their ability to precisely tune heterologous gene expression. Previous work done by our lab used computational approaches to identify 5’ untranslated regions (5’ UTRs) from the genome of *D. radiodurans*, assessing these regions as sequences that might contain putative promoters and ribosome binding sites^30^. We screened 30 candidate 5’ UTR sequences to generate a standardized set of constitutive promoters for *D. radiodurans*. Since in previously published work, the 5’ UTR sequences contained both the promoter and RBS, it was important that we isolated the promoter sequence to directly compare expression between each promoter. To accomplish this task, we ran the set of 30 candidate sequences through two computational promoter-prediction tools, BPROM^37^ and bTSSfinder^38^. We then selected candidates that contained the same transcription start site identified in both BPROM and bTSSfinder, thus reducing our total number of putative promoters to 12 (Fig. 1b, Supplementary Table S1). We replaced the P_GroES_ promoter in the pAT00.20.001-sfGFP plasmid with each putative promoter sequence and measured sfGFP fluorescence after 24 hours of culturing. Across the 12 new promoters and the P_GroES_ promoter, we observed an approximate 45-fold range of fluorescence, with P_GroES_ being the relatively strongest promoter within the set. Importantly, 8 of the 12 new sequences demonstrated sfGFP fluorescence significantly greater than background fluorescence of R1 *D. radiodurans* cells in liquid media (Fig. 1c).

Concurrently, we screened the well-established Anderson Promoter Collection (https://parts.igem.org/Promoters/Catalog/Anderson), because sequences in this catalogue have been shown to work in a wide range of bacterial species^39–42^. We again replaced the P_GroES_ promoter with each of the promoters from the Anderson Promoter Collection and measured sfGFP signal after 24 hours. All the Anderson Promoters demonstrated significant fluorescence above the background of R1 cells (Fig. 1d). Interestingly, the relative strengths of the promoters differed from what was previously observed in *E. coli*. While the J23113 promoter remained the weakest in both *E. coli* and *D. radiodurans*, the J23104 promoter demonstrated the strongest fluorescence in *D. radiodurans*, compared to J23119 and J23100 being the strongest in *E. coli*. Importantly, the J23104 promoter demonstrated even stronger fluorescence in R1 compared to the P_GroES_ promoter when compiling the two sets of promoters into a single graph (Fig 1e) which was created for ease of comparison between the two promoter sets. In total, the suite of 28 promoters provided precise tunability across a 45-fold signal range (Fig. 1e, Supplementary Table S1). With this newly defined set of promoters, targeted gene-expression can be more precisely tuned. We provided a compiled subset of plasmids containing these constitutive promoters that offer 45-fold range of plasmid-based gene expression in Table 1 (Table 1 - pAT00_33_001_sfGFP, pAT00_28_001_sfGFP, pAT00_08_001_sfGFP, pAT00_11_001_sfGFP, pAT00_04_001_sfGFP).

**Table 1.**
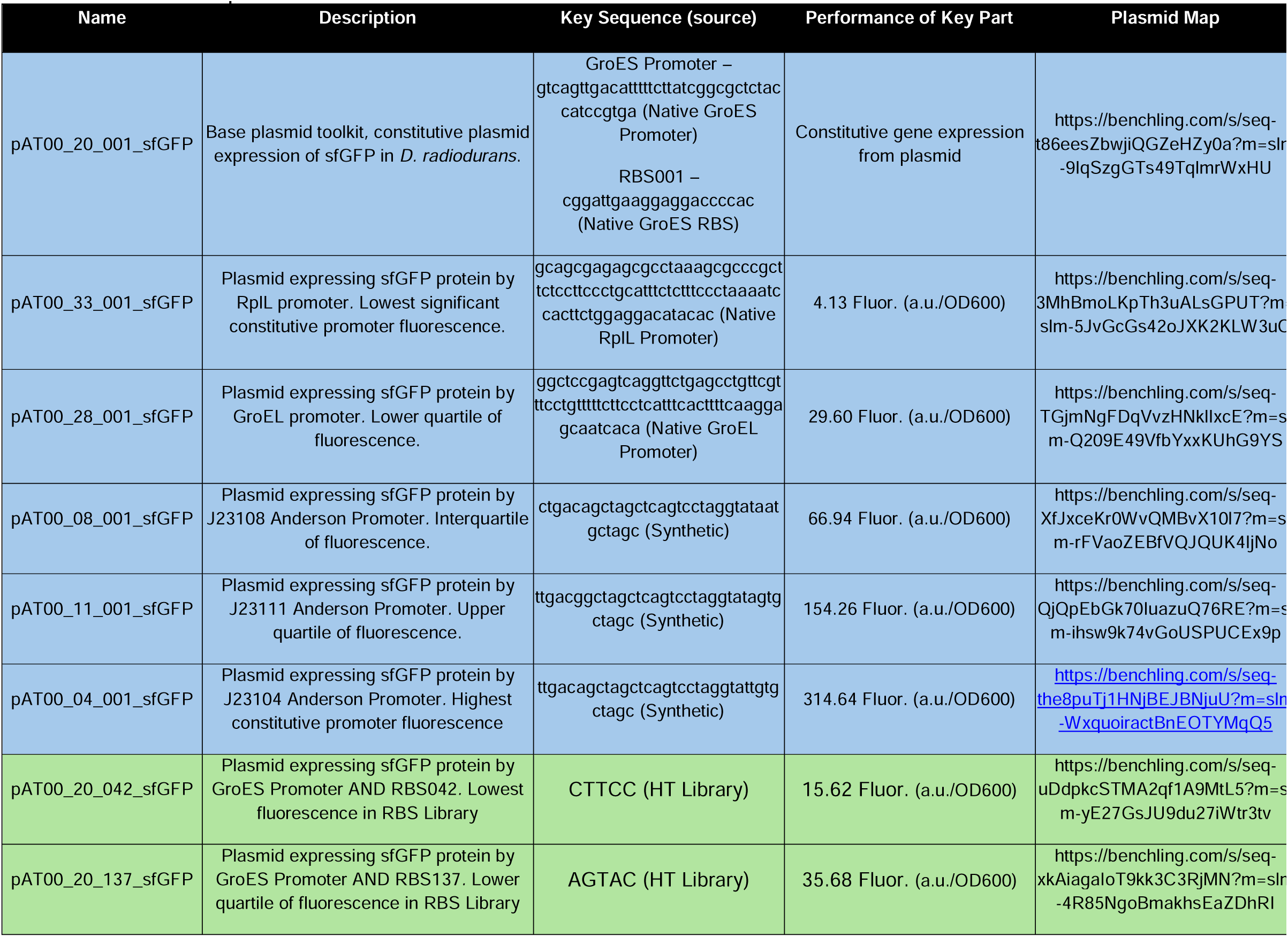
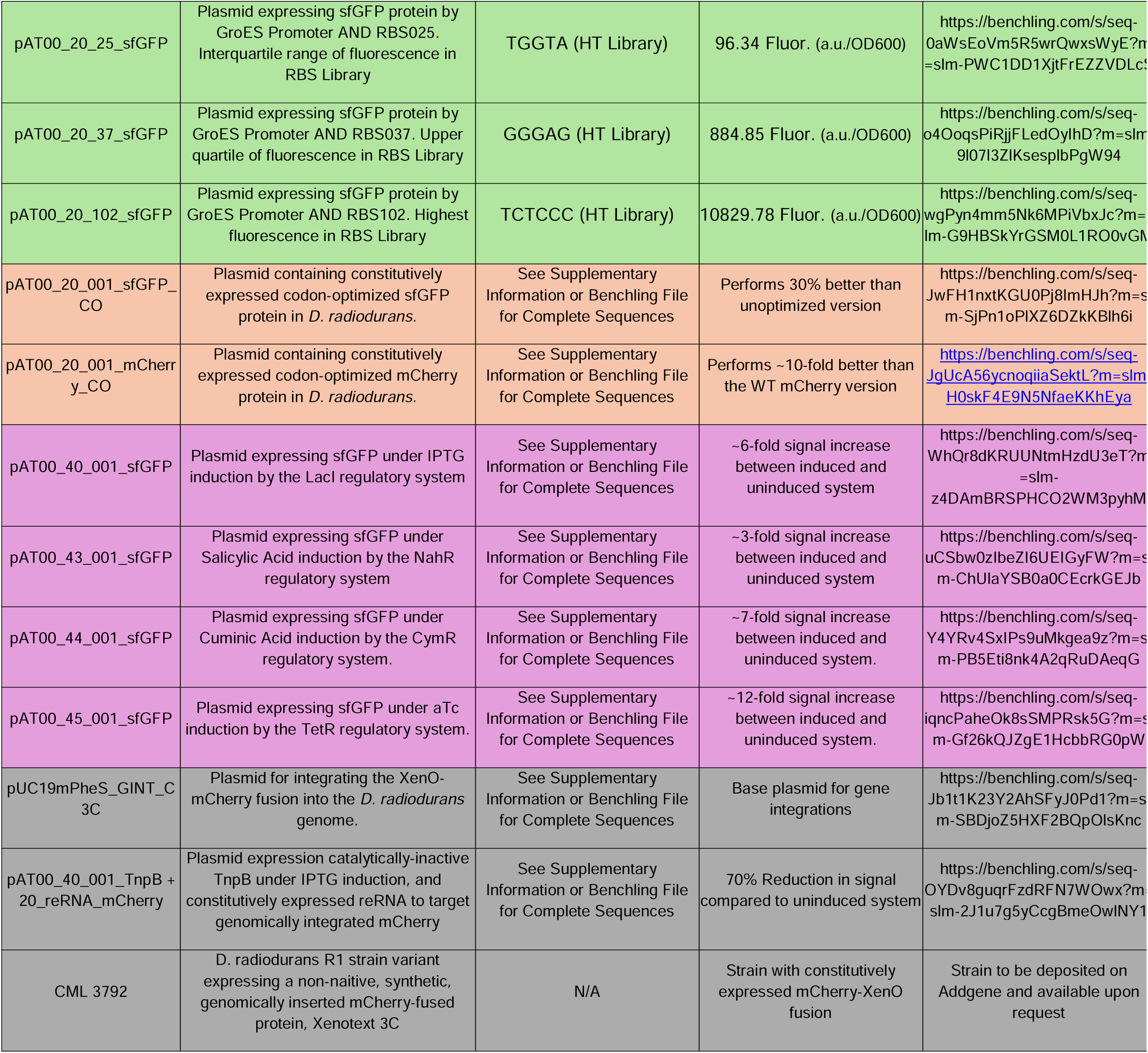
Compiled Plasmids and Strains from the Toolkit of Particular interest to the field. Plasmids and strains were compiled to provide most essential plasmids and strains derived from this toolkit to provide future users with maximum range of control and tunability. Sections are color-coded as the following: Light Blue – Constitutive Promoters, Light Green – RBS Library, Orange – Codon Optimized Genes, Pink – Inducible Promoters, Grey – Genomic Regulatory Plasmids. Sequences of interest, the relative performance, and link to a Benchling plasmid map are also provided.

### Establishing a composable Ribosome Binding Site (RBS) Library for translation control

After establishing a suite of promoter sequences, we developed a suite of well-defined ribosome binding sites (RBS). Earlier computational models have been developed to accurately predict RBS strengths in model organisms, such as *E. coli* ^43–45^. However, these predictive models are not optimized for *D. radiodurans*. Therefore, we chose to establish a library of composable ribosome binding site sequences for tunable translational control. To do so, we adapted a protocol previously utilized for generating an RBS Library in the non-model organism *Cupriavidus necator*^46^. In short, we replaced the GGAGG sequence in the GroES ribosome binding site sequence with an NNNNN sequence pool (N representing any nucleotide) and transformed the library into *E. coli*. During the transformation, we observed that at least 1,024 single colonies formed, ensuring that each potential 5-nucleotide sequence could be represented in the library. Six individual colonies were sent for sequencing, and all 6 plasmids contained different ribosome binding site sequences. These results gave us confidence that our library generated sufficient sequence diversity. The purified library was transformed into *D. radiodurans* R1, and 136 colonies were selected for screening. We chose 136 colonies because this number was greater than 10% of the potential sequence space but still constituted a manageable size such that we could test every sample in a single experiment (Fig. 2a). From our 136 samples, 127 generated fluorescent signals, which became our final library. From those 127 constructs, our library achieved a 963-fold signal range between the weakest, RBS93, and the strongest, RBS102 (Fig. 2b, Supplementary Table S4). Additionally, all constructs demonstrated fluorescence significantly greater than R1 cells alone. After screening the library, we sequenced each colony and estimated the predicted RBS strength for each sequence in the library using the DeNovoDNA RBS Calculator. We found poor correlation (r^2^ = 0.02) between the predicted RBS strength and the fluorescent output of each construct (Supplementary Figure S1). Lastly, we compiled key plasmids containing a subset of the RBS Library thatprovide the largest range of control in Table 1 for ease of access (Table 1 - pAT00_20_042_sfGFP, pAT00_20_014_sfGFP, pAT00_20_025_sfGFP, pAT00_20_037_sfGFP, pAT00_20_102_sfGFP).

**Figure 2.**
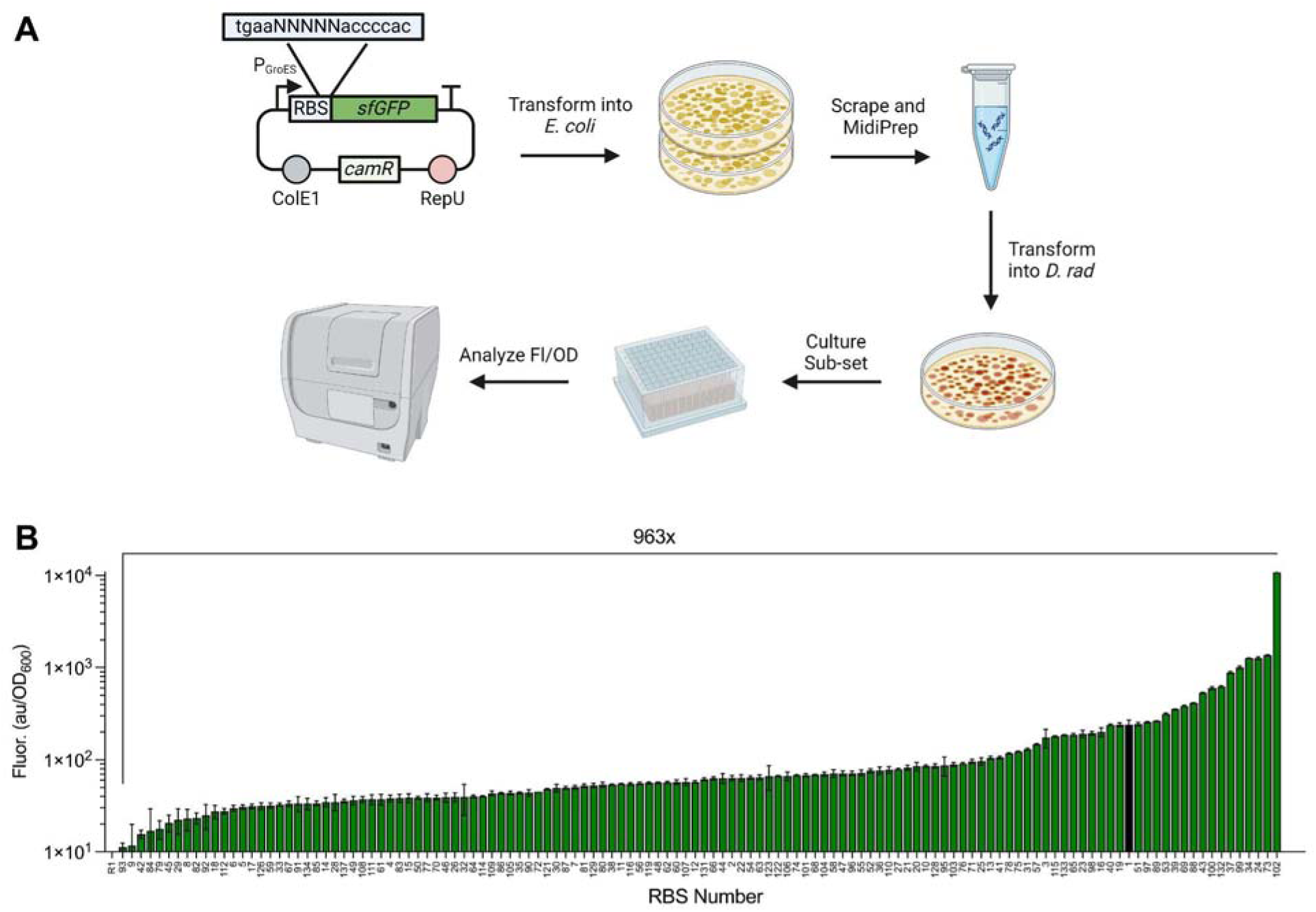
Synthesizing a composable Ribosome Binding Site (RBS) Library for translation tunability in *D. radiodurans.* A) Workflow for generating the RBS Library. Briefly, the GGAGG sequence in the GroES RBS was replaced with an NNNNN sequence, where N is any nucleotide. The library was transformed into *E. coli* then purified via MidiPrep. The purified library was transformed into *D. radiodurans*, and sfGFP signal was measured after 24 hours of growth for 127 colonies in addition to the original RBS sequence. B) Relative fluorescence of sfGFP for each of the 127 RBS sequences. The original RBS sequence is denoted as the black bar on the plot. Overall, the resulting library generated a 963-fold range of sfGFP expression.

### Analyzing codon usage and start codon sequence preferences for codon optimization of heterologous genes

Because *D. radiodurans* has a high GC-content (~66%) relative to microbes such as *E. coli* (50%), codon preferences could significantly differ between these organisms, meaning the sequences of the heterologous genes developed in *E. coli* could lead to imperfect protein translation. Therefore, we wanted to build a codon optimization tool that could recode sequences to match codon usages in *D. radiodurans.* To build our codon optimizer, we leveraged the codon usage data for *D. radiodurans* collected by the Lowe Lab^47^. We then developed a program that would analyze the DNA sequence for each codon in a CDS and modify the sequence as needed to ensure each codon of the CDS corresponded to the sequence with the highest usage rate in *D. radiodurans* (code provided in Supplementary Material). To validate this approach, we ran 4 fluorescent reporter sequences (*sfgfp, mcherry, morange*, and *mturquoise*) through our program to optimize for expression in *D. radiodurans* (Supplemental Methodologies, Supplementary Table S3). We then compared fluorescence of the original (WT) proteins against their codon-optimized counterpart (Figure 3A-D). mCherry, mOrange, and mTurquoise all showed a significant increase in fluorescence, while sfGFP demonstrated an increase that was not statistically significant. To complement our codon-optimizer tool, we evaluated preferred start codon sequences in *D. radiodurans* and found that the “GTG” start codon led to significantly increased fluorescence compared to the start codons “ATG” and “TTG” in the sfGFP CDS (Figure 3E). Our codon-optimizer tool and our start codon preference work help translate protein-based systems previously developed in other hosts, into *D. radiodurans.* We provide our sfGFP and mCherry codon optimized plasmids in Table 1 for ease of access (Table 1 - pAT00_20_001_sfGFP_CO, pAT00_20_001_mCherry_CO).

**Figure 3.**
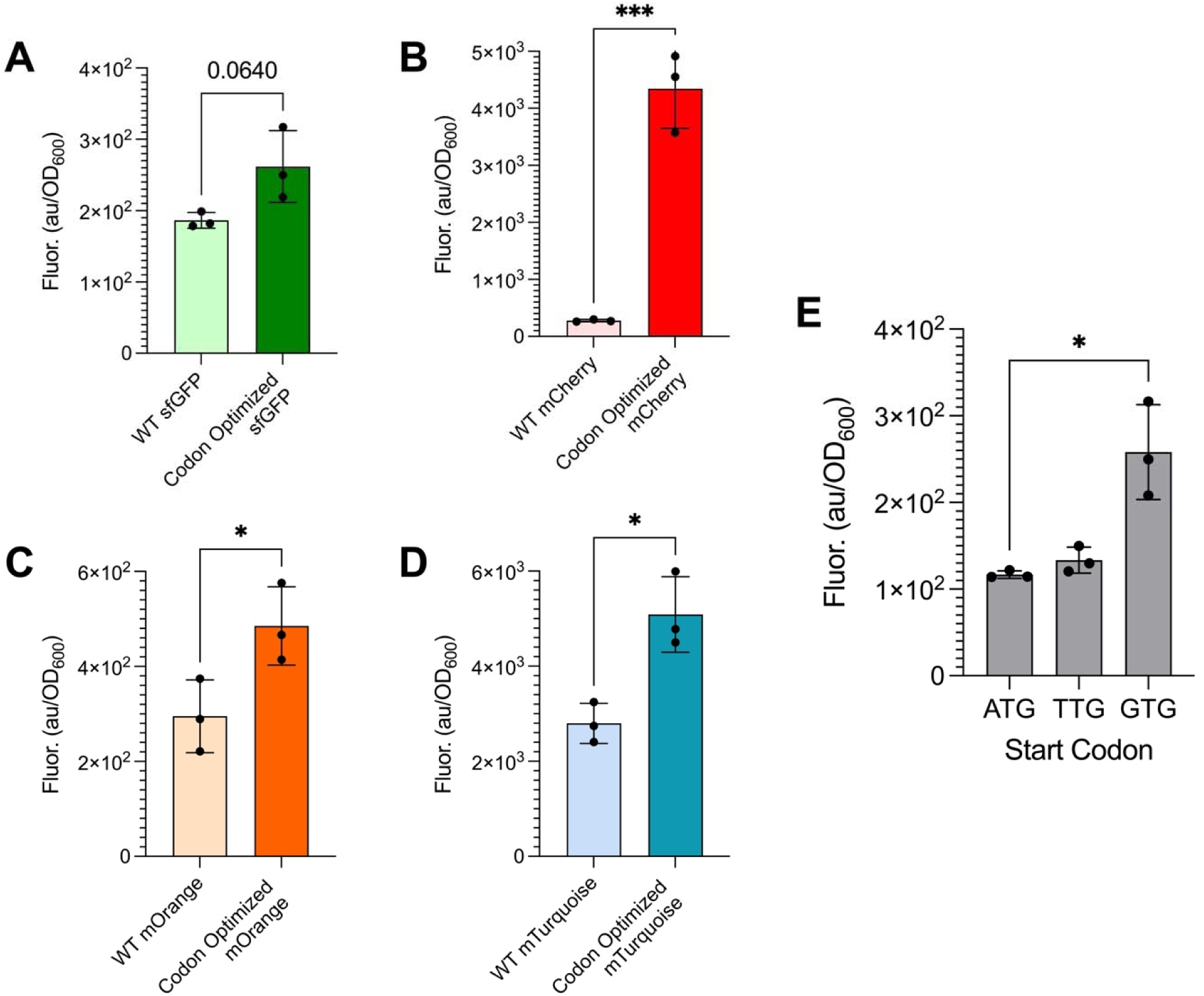
Codon optimization of heterologous genes and start codon preference in *D. radiodurans*. A-D) Relative fluorescence of wild type fluorescent proteins versus their *D. radiodurans* codon optimized version. E) Relative fluorescence of WT sfGFP with the specific start codon ATG, TTG, or GTG. Asterisks indicate a p-value < 0.05 or less using a heteroscedastic T-test between experimental conditions.

### Screening and optimizing transcriptional regulators for small molecule-inducible gene expression

While tuning gene expression through constitutive promoters is an important building block for developing a genetic toolkit, another key step is to develop components for user-controlled gene expression. The gold standard approach is to utilize small molecule-responsive transcriptional regulators such that, upon addition of a small molecule, transcription of a target sequence is initiated. Most transcriptional regulators currently utilized in synthetic biology are optimized for applications in *E. coli* or other model bacteria. As observed in the Anderson Promoter Collection, trends observed in *E. coli* do not necessarily correlate to those in *D. radiodurans.* Currently, there is one established transcriptional regulator system that has been previously shown to function in *D. radiodurans*—the Spac system^31^. In this previous work, the LacI regulatory protein was shown to inhibit transcription from the P_spac_ promoter in *D. radiodurans* and, upon addition of Isopropyl β-D-1-thiogalactopyranoside (IPTG), transcription was activated. While this system has been a valuable foundation to build sophisticated genetic systems in *D. radiodurans*, it is important to have multiple, orthogonal transcriptional regulators, to precisely tune gene expression. Therefore, we screened several well-established transcriptional regulators and then subsequently optimized the most promising candidates for effective performance in *D. radiodurans.* We started with the “Marionette” promoter regulator collection developed by Meyer and colleagues. The “Marionette” collection consists of 14 optimized transcriptional reporters, all engineered to operate orthogonally of each other^48^. We hypothesized that this collection would be an effective place to start, since entries in this catalogue have been shown to function in other non-model organisms^46,49^; moreover, this catalogue enabled us to survey a large number of small molecule-inducible promoters.

To screen the “Marionette” promoter collection, we replaced the P_GroES_ promoter from our pAT00.20.001-sfGFP vector with each of the inducible promoter sequences from the “Marionette” collection, and we inserted each corresponding regulatory protein directly downstream of the rrnBT1T2 terminator. Of the original 14 promoter constructs created in the “Marionette” promoter collection, we were able to clone 9 of them for our plasmid-based system. With the 9 inducible promoter designs, we titrated each respective small molecule inducer at different concentrations in liquid culture, and then we measured the fluorescent signal of the sfGFP protein after 24 hours of culturing at each induction condition. The anhydrotetracycline-(aTc), Cuminic acid-(CA), IPTG- and Salicylic Acid (SA)-inducible regulators all achieved at least 5-fold increase in signal between the uninduced conditions and the saturated, induced conditions. Additionally, the IPTG-, aTc-, and SA-inducible constructs demonstrated precise tunability between induction concentrations (Fig. 4a, c, f). The CA-inducible regulator demonstrated rapid increase in signal between the 100 nM conditions and the 1 μM conditions, but then saturated at all greater induction concentrations (Fig. 4h). Ultimately, after screening 9 systems, we identify four inducible promoter systems (IPTG, SA, CA, and aTc) that offer up to a 12-fold range of titratable gene expression via plasmids (Table 1 – pAT00_40_001_sfGFP, pAT00_43_001_sfGFP, pAT00_44_001_sfGFP, and pAT00_45_001_sfGFP).

**Figure 4.**
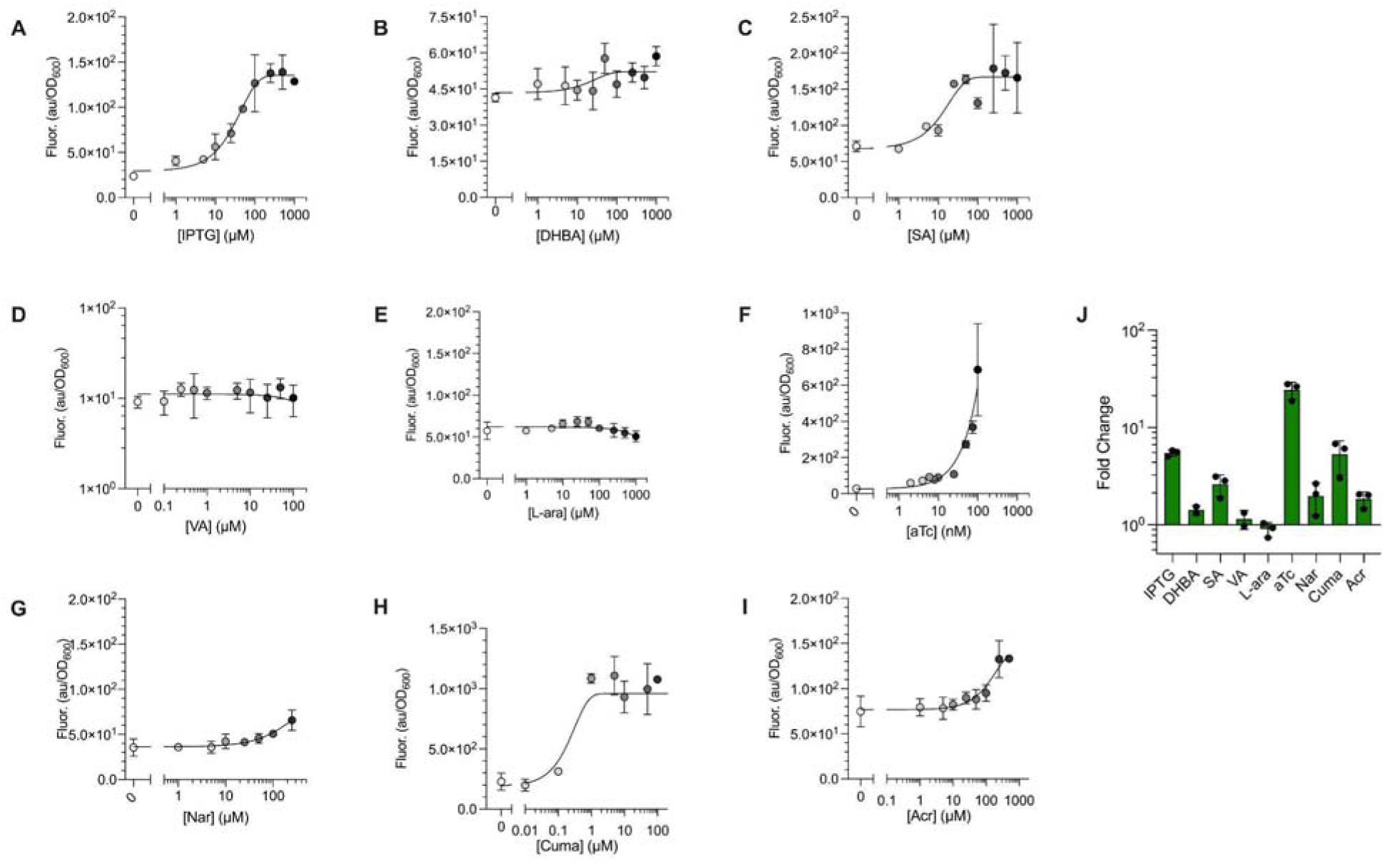
Screening and optimizing small molecule-inducible transcriptional regulators in *D. radiodurans*. A-I) Titration curves for each small molecule inducer measured by sfGFP signal 24 hours after induction. J) Fold change in sfGFP expression for each small molecule inducer between the uninduced and fully induced conditions.

### Integrating stable expression of synthetic genes and biological signals in D. radiodurans

In addition to plasmid-based genetic components, developing tools for precise genome editing is critical for making any microbe a versatile biological chassis given that integrations eliminate the need for plasmid-based expression and therefore eliminates the need to culture with antibiotic selection. There has been recent work to develop protocols for genomic deletions in *D. radiodurans*, all of which rely on conjugation-based techniques^50^, leveraging the bacterium’s natural double-crossover and homologous recombination abilities^15,33,51^. Additionally, due to its resistance to both oxidative stress and ionizing radiation, *D. radiodurans* is a top candidate for biological storage of data within its genome. However, there has yet to be a well-established method for efficient genome integrations in *D. radiodurans*.

To develop our integration protocol, we selected a large (> 2kb) non-native synthetic protein, nicknamed “The Xenotext” (XenO) as a proof of concept. XenO is a synthetic, his-tagged protein designed in collaboration with Dr. Christian Bök, an experimental poet. XenO was developed for the purpose of encoding messages into the genome of a stress-resistant bacterium (Supplementary Table S5)^52^. We also chose to fuse XenO to a fluorescent mCherry protein, since the integrated XenO synthetic protein could then be tracked over time and easily identified without genome sequencing. The application of encoding messages through bacterial genomes has been previously established in model organisms^53,54^, and the stress-resistance properties of *D. radiodurans* position this bacterium as an ideal candidate for long-term biological storage of information regardless of external conditions^55^. With this goal in mind, we established and optimized a protocol for the integration of information into the genome of *D. radiodurans*. It is worth noting that a challenge with genomic engineering of *D. radiodurans* lies on the fact that it has multiple copies of its genome ranging from 8-10 at exponential phase^56^.

Our protocol leverages the double-crossover, homologous recombination mechanism that *D. radiodurans* employs for genome repair and horizontal gene-transfer^57^. We chose to insert the XenO-mCherry fusion between the DR_0963 and DR_0937 coding sequences in the R1 genome, because there are approximately 200 base pairs between the two genes, and because there are no annotated open reading frames (ORFs) on the reverse strand. We hypothesized that these two factors combined would minimize disruption to the bacterial genome. The 1000 nucleotides upstream and downstream of the insertion site were then amplified from the genome to flank the 5’- and 3’-regions of the synthetic sequence, which contained the XenO-mCherry fusion sequence and a kanamycin resistance marker. This marker contained flanking sequences of both lox66 and lox71 for subsequent removal of the resistance marker, if desired (Fig. 5a). We then cloned the entire design into the pUC19mPheS plasmid (Fig. 5a). To begin the process of integration, we transformed this plasmid into *D. radiodurans* R1 cells (where the homology arms are recognized and homologous recombination occurs), then incubated the sample overnight. Strains containing the XenO-mCherry integration were selected by using TGY agar plates supplemented with 16µg/mL kanamycin and 5 mM 4-chloro-phenylalanine. The 4-chloro-phenylalanine and kanamycin selected for single colonies that had undergone successful recombination and had cured out the pUC19mPheS plasmid during the overnight incubation (Fig. 5a). Using this protocol, we were able to successfully integrate our XenO-mCherry fusion sequence into the genome of *D. radiodurans* with 70% efficiency after one round of selection, 86% efficiency after two rounds of selection, and 100% efficiency after three rounds of selection, for an average efficiency of 60%. These integrations were confirmed with colony PCR (Fig 5b) as well as full genome sequencing (Supplementary Figure S2). We chose to repeat this process to evaluate the number of rounds required to have all screened colonies fully saturated with the synthetic integration. To ensure proper functionality of our mCherry fusion protein, we tested multiple variations of the XenO sequence and found that fusing mCherry to the C-terminus of the XenO version 3 generated the greatest fluorescence signal (Supplementary Figure S2). We then validated the production of the XenO-mCherry fusion by measuring fluorescence of our strain in liquid culture (Fig. 5c), as well as by microscopy (Fig. 5d). Our results demonstrated that genome integrations of at least 2kB could be achieved using a single plasmid in a two-step process that yielded 60% efficiency for target integration and present a plasmid (Table 1 - pUC19mPheS_GINT_C3C) that can be easily modified for the desired target of the user.

**Figure 5.**
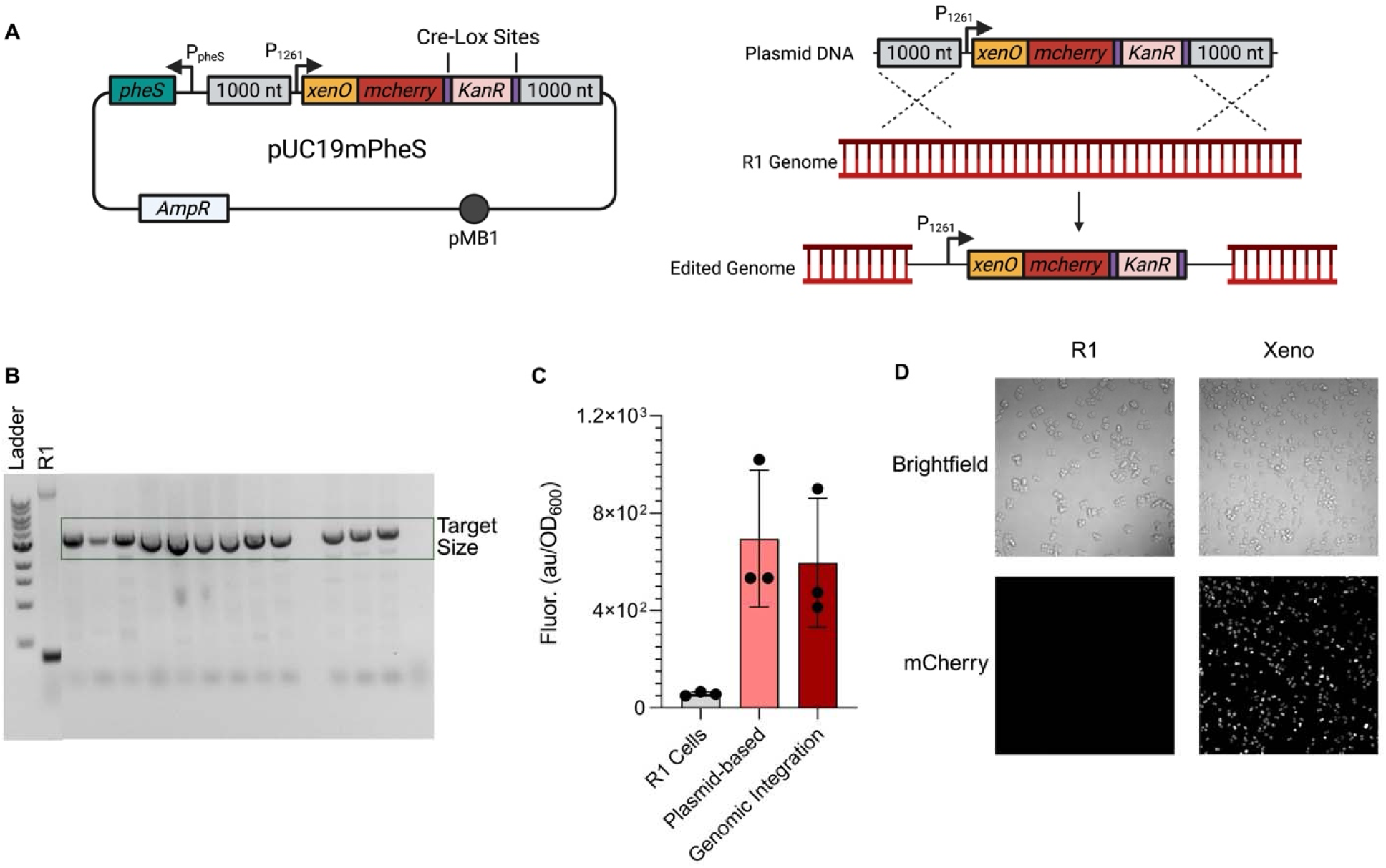
Genomic Integration protocol in *D. radiodurans*. A) pUC19mPheS plasmid schematic to integrate XenO-mCherry fusion (left) and mechanistic diagram of how region of interest is inserted into the *D. radiodurans* R1 Genome (right). B) cPCRs of colonies after two rounds of screening. The successful desired insert length is approximately 3 kB (highlighted in green), colonies with unsuccessful insertion attempts do not have that band (lanes 12 and 16). C) Fluorescence of R1 cells, a plasmid containing the XenO-mCherry fusion, and the final XenO strain containing the integrated mCherry fusion after 24 hours of culturing. D) Microscopy of R1 and XenO strains under brightfield imaging, as well as voltage for 561 nm to excite mCherry fluorescence.

### Repurposing the transposon-associated protein TnpB for programmable genetic knockdowns

There have been considerable efforts towards genomic editing technology in bacteria in the last decade, particularly through Cas-based tools that allow up and down regulation of target genes in bacteria. Most efforts have leveraged dCas9 (or engineered derivatives) to selectively silence or activate gene expression^58–60^. Interestingly, expressing dCas9 in *D. radiodurans* is toxic and leads to a significant loss in colony forming units, and growth defects in liquid culture^32^. Therefore, finding alternate approaches for targeted gene-repression in *D. radiodurans* is essential for establishing similar technology. Recently, the transposon-associated protein TnpB was identified in *D. radiodurans* as an RNA-directed nuclease capable of being programmed to cleave target DNA sites in cells^61^. In this work, we showed that, within *D. radiodurans*, the catalytically inactive mutant (D191A, which we termed dTnpB), could be repurposed and leveraged in the same way that dCas9 engineering applications have been used in other bacteria^58,59^.

To generate construct, we replaced the *sfgfp* CDS in our aTc-inducible plasmids and IPTG-inducible plasmids with that of the *dtnpB* CDS. We selected the aTc-, and IPTG-inducible systems, because these systems were two of the most titratable inducible promoter systems that we screened. Additionally, we integrated a constitutively expressed guiding RNA sequence defined by Karvelis and colleagues^61^, incorporating this sequence as the right end element RNA (reRNA) downstream of our regulator sequence (Fig. 6a). The reRNA sequence was approximately 150 nucleotides, wherein the last 16 nucleotides of the 3’-end contained the guide sequence for TnpB. According to Karvelis and colleagues, this guide sequence directs TnpB to cut at a “TTGAT” sequence, functioning similarly to a PAM site for Cas9 systems^61^. For our work, we initially designed the reRNA to target a sequence of 44 nucleotides upstream of the start codon: i.e. within the constitutive promoter upstream of our integrated XenO-mCherry sequence (Fig. 6a). Our reRNA sequence also contained a self-cleaving hepatitis delta ribozyme (HDV) sequence to ensure that the final 16 nucleotides of the reRNA sequence were our desired targeting sequence, which was shown to be critical for effective targeting^61^. To evaluate the ability of the dTnpB system to repress genomic expression of our target sequence, we tracked mCherry signal in liquid culture up to 24 hours after induction (Fig. 6b-d). Within four hours of dTnpB induction, we observed an approximate 30% reduction in mCherry signal in the IPTG-inducible construct. Maximum repression occurred 24 hours post-induction, when we observed a range of signal reduction (from 66% to 87%) in mCherry fluorescence between our induced samples and uninduced samples (Fig 6e). Ultimately, we constructed a plasmid-based system for dTnpB-guided repression of genomic targets (Table 1 - pAT00_40_001_TnpB + 20_reRNA_mCherry) and these results serve as preliminary evidence that the catalytically inactivated dTnpB could be leveraged for targeted gene-silencing in *D. radiodurans*.

**Figure 6.**
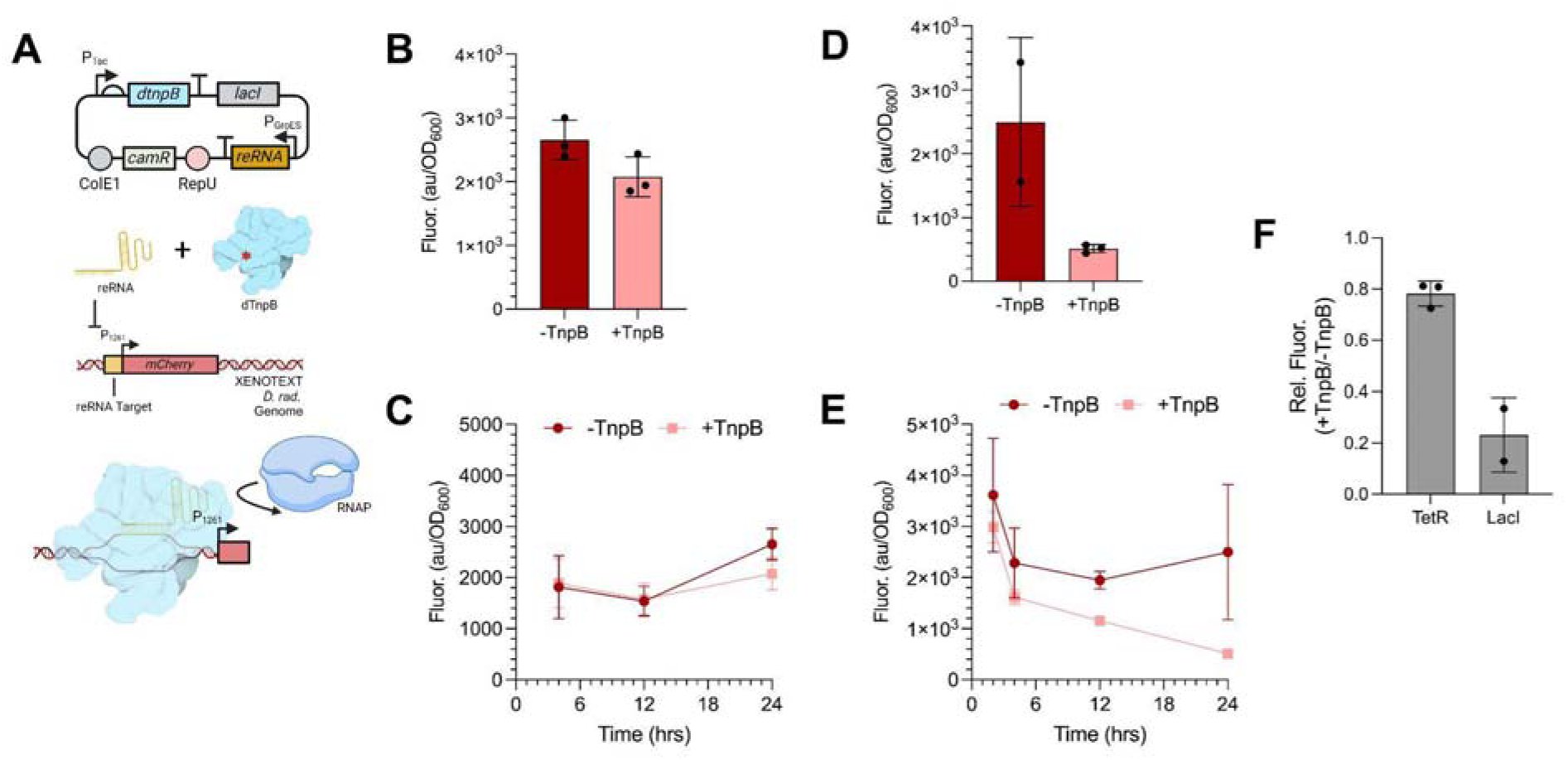
Programmable dTnpB-assisted interference system for repression of genomic targets in *D. radiodurans*. A) Plasmid map and schematic for the dTnpB-based system. dTnpB is under small molecule induction, and the reRNA is constitutively expressed via the P_GroES_ promoter. Upon expression of both components, the dTnpB and reRNA complex targets the promoter sequence of constitutively expressed mCherry fusion protein of the XENOTEXT strain. The binding of the dTnpB occludes RNA polymerase from binding and reduces transcription, therefore reducing mCherry signal. B) mCherry signal of cultures with dTnpB expression induced (aTc) and uninduced after 24 hours of growth. C) Time course of mCherry signal between the cultures with and without dTnpB induction under aTc. D) mCherry signal of cultures with dTnpB expression induced (IPTG) and uninduced after 24 hours of growth. E) Time course of mCherry signal between the cultures with and without dTnpB induction under IPTG. F) Relative mCherry signal of cultures under tetR and LacI regulation.

## Discussion

In this work, we established a genetic toolkit in *D. radiodurans* that expands the available repertoire for origin of replications, standardized constitutive promoters, RBS sequences, and small molecule-inducible promoters. Additionally, we established a process for effective gene insertions as well as a tool for programmable gene repression. These new genetic components improve tunability of heterologous gene expression as well as the ability to edit and tune native gene expression. We have compiled a final list of the most versatile plasmids and strains and indicate their relative performance context in Table 1 below for ease of access and request. As a note, all sequences used in this study are compiled in the supporting information.

Our standardized constitutive promoters provide a 45-fold range for tuning gene expression (Figure 1) — a range comparable to other genetic toolkits recently established for non-model organisms^24,46,49,62^. These constitutive promoters can be used to express a heterologous target on a plasmid in *D. radiodurans*, making them useful in developing more complex biological control schemes. The ability to vary constitutively expressed components, such as the LacI regulatory protein or a small RNA transcript, provides the opportunity to improve coordination between components and can reduce the rate of failure as system complexity increases^63^.

Separately, of the nine inducible promoter systems that we tested for plasmid-based control, the IPTG-, aTc-, CA-, and SA-inducible promoter systems from the Marionette collections inducible promoter systems all demonstrated at least 3-fold activation (Figure 4a, c, f, h, j), which is similar to magnitudes of fold activation for inducible promoter systems previously developed in *D. radiodurans*^31^. It is important to note that 5 of the inducible promoter systems did not demonstrate any titratable signal control (Figure 4b, d, e, g, i). This could be due to many factors such as low promoter strength or overexpression of the repressor protein. These features could be further explored if additional inducible promoter systems are needed; however, four functional systems provide a solid foundation for inducible transcription-based control in *D. radiodurans*. Additionally, because these systems are derived from the “Marionette” collection, they were designed to be orthogonal; therefore, they should experience minimal crosstalk between the systems, allowing for improved circuit dynamics. Overall, these inducible promoters lay the foundation of more complex genetic control, such as two-input logic gates or other bacterial computation, including BANDPASS Filters or three-input circuits^40,64^.

Next, we generated a library of 127 composable RBS sequences that offered a 963-fold range of translational control (Figure 2b) — a standard that meets or significantly exceeds previous RBS libraries generated in non-model bacteria^46,65^. This RBS library is also the first of its kind in *D. radiodurans* and can be used in concert with the promoter library for extremely precise genetic tuning. In many model microbes, there are predictive computational tools that can accurately generate sequences for effective RBS tuning. However, in our work we found poor correlation between the *in vivo* fluorescent values and the *in silico* predicted RBS strengths for our library (Supplementary Figure S1). This may be due to a variety of variables: the high GC content of *D. radiodurans*; the mechanism translation initiation; the environmental conditions; or other confounding factors. Importantly, these results demonstrate that additional work is required to understand the relationship between RBS and translation initiation in *D. radiodurans*. For now, our library provides the most effective option for precise RBS tuning. Additionally, our RBS library could service as a basis to develop computational models that more accurately predict RBS strength in *D. radiodurans* in the future. In addition to our RBS Library, our codon-optimization tool will be able to work in concert to ensure both efficient translation initiation as well as preferred codon usage to increase the likelihood of a correctly synthesized protein. Overall, this RBS library and codon-optimization tool will provide opportunities for precise optimization of heterologous pathways and other genetic tools originally developed in other host systems.

We also sought to identify novel origin of replication (ori) sequences that were functional in *D. radiodurans*. Currently, the only plasmid ori is RepU derived from pRAD1. Identifying other origins of replications would allow for multi-plasmids strains or allow for users to vary copy number based on the ori, similar to how pMB1, p15a, and pSC101 are leveraged in *E. coli.* To find other origins of replication, we initially searched for sequences similar to *repU*, however no other homologous sequences were identified. Therefore, we took a rational engineering approach (Supplementary Figure S3a) by selecting three different subsets of origin of replications: (i) common origins of replication used in *E. coli* (ColE1, and p15a), (ii) broad host range origins of replication (BBR1, R2K, RK6, RO1600), and (iii) origins of replications from plasmids used in *Thermus thermophilus* (UB110, MY1, TT8, TsYS45), a similar high GC-content extremophile. Of these origins of replication, we found that UB110 and RO1600 generated colony forming units in *D. radiodurans*, and those single colonies could be grown in liquid media to produce a fluorescent sfGFP signal (Supplementary Figure S3b). However, these liquid cultures could not be passaged a second time nor could copy numbers be quantified from these oris. Therefore, the RepU ori remains the only ori that stably maintains plasmids in *D. radiodurans*. Discovering or engineering novel oris compatible in *D. radiodurans* will be important for expanding the utility of plasmid-based gene expression in *D. radiodurans*.

In parallel to building plasmid-based tools, we established methods for editing and tuning expression from the genome of *D. radiodurans.* Our protocol for gene integration leverages the natural competency and recombination of *D. radiodurans* to create novel gene-integration systems. Moreover, this approach achieves a 60% integration efficiency with only two rounds of screening (Figure 5). These results are comparable to Cas-based genome engineering approaches in model bacteria^66,67^. Previous efforts have established a process for knocking out genes in *D. radiodurans*^50^; however, our system is the first of its kind to achieve large (>2kB) genomic integrations. Additionally, our system only relies on a single plasmid and combines the genomic integration and plasmid curing into a single step (Figure 5a), reducing the time to develop new strains. This technology opens the door to engineering strains of *D. radiodurans* that can be used in large-scale metabolic engineering or bioprocessing applications, as synthetic systems integrated into the genome eliminate the need for continual plasmid maintenance.

Concurrently, our programmable dTnpB-assisted gene repression system is equally useful, because it allows for targeted repression within the genome. This system achieves a range of repression between 66% and 87% of its target sequence relative to the uninduced system when using the LacI to control expression of the TnpB protein (Figure 6d, e) — a result comparable to Cas-based systems in both model organisms and non-model organisms^41,59,68–70^. This approach is also useful, because it can help simulate genomic deletions without requiring the labor to ensure that the gene is knocked out of all four chromosomes of *D. radiodurans*. One particular systems is the histone-like protector, HU, which plays a role in nucleoid compaction, but cannot be knocked out^71^. Therefore, this TnpB-based knockdown system could serve as a way to evaluate the phenotype of a genotype that cannot be otherwise synthesized *in vivo*. Previous work has demonstrated that dCas9 regulatory systems can function in *D. radiodurans*; however, they greatly alter growth and prevent growth in liquid culture^32^. These effects are not observed with our application of the TnpB-based system. This outcome may occur because TnpB is native to *D. radiodurans*, conferring an additional benefit from using a protein in its native environment. Although TnpB has been recently used as a programmable gene-editing tool^61,72^, our work is the first demonstration of utilizing the catalytically inactive version of TnpB to repress genes from any genome in a targeted fashion. This system may be transferable to other organisms that also cannot utilize Cas-based system or other species of interest similar to *D. radiodurans* — species that, for example. do not have a robust gene-knockdown protocol, such as *Deinococcus geothermalis*. Additionally, this system provides a foundation for developing programmable gene-activation by fusing protein activators, such as transcription factors or subunits of RNA polymerase, to the dTnpB protein.

Overall, we have built a toolkit that allows for precise user-control of gene-expression via the plasmid or the genome, as well as a novel protocol for editing the genome of *D. radiodurans*. We anticipate that this toolkit will be leveraged in future work to advance fundamental research and engineering applications of *D. radiodurans.* Importantly, all of the sequences, scripts, and methods utilized in this paper are provided in the Methods as well as in Supplementary Materials, and we encourage the research community to utilize these tools. This toolkit aims to advance the utility of *D. radiodurans* in bioprocessing, biomanufacturing, and biological data storage, as well as other applications in fundamental research and synthetic biology.

## Materials and Methods

### Bacterial strain and culture conditions

Escherichia coli strain DH5α and its plasmid-containing derivatives were cultured aerobically at 37°C in Luria-Bertani (LB) liquid media (10 g/L tryptone, 10 g/L NaCl, and 5 g/L yeast extract) or LB solid (1.5% agar) medium. When necessary, chloramphenicol was used at a concentration of 34 μg/ml for E. coli strains. *Deinococcus radiodurans* R1 (ATCC 13939) and its plasmid-containing derivatives were cultured aerobically at 32°C in TGY liquid media (1% tryptone, 0.1% glucose, 0.5% yeast extract) or TGY solid (1.5% agar) medium. When necessary, antibiotics were used at a concentration of 34 μg/ml (Chloramphenicol in *E.* coli strains and 16 μg/ml (Kanamycin), 3.4 μg/ml (Chloramphenicol) for *D. radiodurans* strains All strains utilized and tested are listed in Supplementary Table S7.

### Plasmid Construction

Plasmids were synthesized using Gibson Assembly of PCR fragments. Either Q5 Hot Start High-Fidelity 2x Master Mix (New England Biolabs) or Phusion Polymerase (New England Biolabs) were used. For Gibson Assemblies, NEBuilder® HiFi DNA Assembly Master Mix (New England Biolabs Inc.) was used to assemble full plasmids from their parts. Plasmids were purified from *E. coli* using the Qiagen QIAprep Spin MiniPrep Kit (Cat. No. 27104). Plasmids developed in this study are contained in Supplementary Table S7. Primers used in Plasmid development are contained in Supplementary Table S8.

### Measuring Fluorescent Signal from liquid culture of *D. radiodurans*

To measure fluorescence of our synthetic toolkit constructs in *D. radiodurans*, sequence verified single colonies were grown in 5 mL of TGY supplemented with chloramphenicol for 24 hours in a shaking incubator at 32 °C. The next day a new 5 mL culture of TGY supplemented with chloramphenicol was inoculated with the saturated overnight, such that the final OD_600_ was approximately 0.1- 0.2. These samples were grown for another 22 to 24 hours in a shaking incubator at 32 °C. Next, 1 mL of culture per sample was centrifuged for 5 minutes at 3000 rpms. The supernatant was discarded, and the remaining pellet was resuspended in 200 µL of 1x PBS, which was prepared according to the Cold Springs Harbor Protocol (^63^). The resuspension was then transferred into a Greiner 96-well µClear, F-Bottom, medium binding plate (Ref. No. 655096). Fluorescence was measured using a BioTek Cytation 3 Plate Reader. For sfGFP the excitation and emission wavelengths were 487 nm and 513 nm. For mCherry the excitation and emissions wavelengths were 579 nm and 616 nm. For mTurquoise the excitation and emission wavelengths were 434 nm and 479 nm. For mOrange the excitation and emission wavelengths were 544 nm and 570 nm. To measure OD_600_, absorbance was taken using a wavelength of 600 nm. All samples were taken in biological triplicate, unless otherwise specified. As a blank, 200 µL of PBS was used.

### Plasmid purification out of *D. radiodurans*

Plasmids were purified from *D. radiodurans* using a three-phase protocol. First, single colonies were grown in 5 mL of TYG supplemented with appropriate antibiotics for 20-24 hours in a shaking 32 °C incubator. The entire volume of the cells was then pelleted via centrifugation at 3000 rpm for 5 minutes. After centrifugation, the supernatant was discarded and the pellet was resuspended in 1 mL of TGY and incubated at 95 °C for 10 minutes. After incubating, the cells were centrifuged again for 5 minutes at 3000 rpms, and the supernatant was discarded. Next, the pellet was resuspended in 250 µL of Qiagen P1 Buffer supplemented with 6 mg/mL Chicken Lysozyme (Muramidase - US Biological), which was freshly mixed. The suspension was incubated for 15 minutes at 37 °C, and could be extended up to 1 hour of incubation for hard to lyse samples. Next, these samples were purified using Qiagen QIAprep Spin MiniPrep Kit (Cat. No. 27104) starting from Step 3, which is the addition of Buffer P2 Step. As a last note, letting the P2 lysis step proceed for at least 3 minutes generated the highest plasmid yields. Plasmids were then sequenced using Plasmidsaurus or Eton Bioscience Inc.

### RBS Library Construction

To create the RBS Library, a protocol was derived from previous work (^41^). To summarize, the pAT00.20.001 plasmid was amplified except for the region that includes the P_GroES_ promoter, the RBS, and the first 153 nucleotides of the *sfgfp* sequence. This sequence was incubated overnight with DnpI (New England BioLabs) enzyme to digest any parent plasmid. Then, it was purified using the QIAquick PCR Purification Kit (Qiagen). A pool of gBlocks containing the P_GroES_ promoter through the 153^rd^ nucleotide of the *sfgfp* CDS sequence was synthesized via Twist Bioscience, however the “GGAGG” sequence within the RBS was replaced with “NNNNN”, where N means any nucleotide. This sequence is listed in Supplementary Table S5. Our pool of gBlocks had 1024 combinations of possible RBS sequences. The two purified fragments were placed into NEBuilder® HiFi DNA Assembly Master Mix reactions totaling 20 µL. The fragments were added in a 5:1 molar ratio of RBS insert to pAT00 backbone. Two of these Gibson Assembly reactions were created. The reaction mixtures were then incubated at 50 °C for 1 hour. Next, 15 µL of product from each reaction were desalted using MF-Millipore 0.025 µm MCE Membranes (Millipore Sigma) for 15 minutes. Then, 10 µL of desalted Gibson Assembly Product was added to 100 µL of ElectroMAX DH10β cells (ThermoFisher) on for each reaction. The assemblies were transformed via electroporation using Gene Pulser/MicroPulser Cuvettes (Bio-Rad) and the GenePulse Xcell electroporator (Bio-Rad), set at a voltage of 1.8 kV. The corresponding time constants for each transformation were between 4.5-5. The transformations had 900 µL of S.O.C. media added, provided with the ElectroMAX DH10β Kit (ThermoFisher), before being recovered for 90 minutes in a 37 °C shaking incubator. The recoveries were then centrifuged for 5 minutes at 5000 rpm, the supernatant was discarded, and the cells were resuspended before each being plated on LB Agar plates supplemented with chloramphenicol. The petri dishes for the LB Agar plates used were 150 x 15 mm Falcon (Corning) plates. The plates were grown overnight at 37 °C. The next day, colony counts were made between both plates to ensure at least 1,024 single colonies were formed. 6 single colonies were purified via the Qiagen QIAprep Spin MiniPrep Kit (Cat. No. 27104), and sent for sequencing via Plasmidsaurus. All six single colonies had different RBS sequences in the “NNNNN” region, which provided confidence there was sequence diversity in the rest of the library. Next, both plates were scraped using 3 mL of ice-cold LB per plate and plastic plate scrapers. The solution of LB and scraped colonies were transferred into 50 mL of LB supplemented with chloramphenicol, and grown overnight at 37 °C in a shaking incubator. The next day, the culture was split into two 25 mL samples and the libraries were purified using the QIAGEN Plasmid Mini Kit (Qiagen) according to the user guide. 2 µg of Library DNA per sample was added to a fresh aliquot of *D. radiodurans* R1 competent cells and transformed according to the protocol below. The transformations were plated on TGY Agar plates supplemented with chloramphenicol. The petri dishes for the plates used were 150 x 15 mm Falcon (Corning) plates. After single colony formation (~2-3 days), 136 single colonies were grown for 24 hours in 5 mL of TGY supplemented with chloramphenicol. After 24 hours of growth, the fluorescence of each culture was measured in technical triplicate, according to the protocol for measuring sfGFP fluorescence in *D. radiodurans* stated above. Additionally, with the remaining liquid culture, the plasmids were purified and sequenced according to the plasmid purification out of *D. radiodurans* stated directly above. Once all sequences were returned, the RBS strengths were estimated using the Salis Lab RBS Calculator at (https://salislab.net/software/predict_rbs_calculator). The RBS sequences run through the calculator, their predicted strengths, as well as sfGFP signal measured *in vivo* are reported in Supplementary Table S4, and the correlation plot between predicted strength and fluorescent output is available in Supplementary Figure S2.

### Transformation of recombinant plasmids in *D. radiodurans*

Recombinant plasmids were transformed into a chemically competent wild-type (R1 ATCC 13939) *D. radiodurans* strain as previously described (^32^). Briefly, *D. radiodurans* cells were grown overnight in 5 mL of TGY in a 32 °C shaking incubator. The next day, the saturated overnight was used to seed a fresh 5 mL of TGY to a final OD_600_ of 0.2. The cultures were grown to late log phase (OD600 = 1.0), then 955 µL of cells were mixed with 45µL of 1M CaCl2 and 500 µL of ice cold 30% glycerol to create the chemically competent cell line. 300 µL of the mixture were aliquoted into sterile 1.7 mL microtubes (Olympus). Competent cells were either stored at −80 °C for future use immediately. For transformation into the competent cell stock, 1.5μg plasmid DNA was added, followed by an incubation of 1 hour on ice, and a subsequent incubation at 32°C shaking for 1 hour. Cells were then incubated at 32°C shaking overnight in 1 mL of fresh TGY media in test tubes. Post-incubation, cells were plated onto TGY plates with the appropriate antibiotic concentration (above), and incubated for 3 days at 32°C. Transformants were verified via polymerase chain-reaction (PCR) and sequencing (Plasmidsaurus, Eton Bioscience Inc.).

### Construction of genomic integration strains in *D. radiodurans*

The genomic integration strains of *D. radiodurans* were constructed using a modified homologous recombination method reported previously for the deletion of genes^51^. In short, 1kb regions upstream and downstream from the selected genomic insertion site were amplified via PCR from the R1 genome, and cloned into the pUC19mPhes plasmid alongside the gene of interest to be inserted, a fragment containing a kanamycin cassette, and both lox66 and lox71 sequences. All fragments were then gel-purified using QIAquick Gel Extraction Kit (Qiagen) and assembled with a linearized, HindIII-digested, pUC19mPheS plasmid using NEBuilder® HiFi DNA Assembly Master Mix (New England Biolabs Inc.). The resulting plasmid was then transformed into *E. coli* DH5α strain and isolated via QIAprep Spin MiniPrep Kit (Qiagen). The isolate was subsequently transformed into *D. radiodurans* R1 chemically competent cells (above). Mutant strains were selected on TGY agar plates with the appropriate kanamycin concentration and 4-chlorophenylalanine (5 mM) for 3 rounds of selection. The total process selected for cells that underwent homologous recombination to replace the target region with the kanamycin cassette, and no longer contained the pUC19mPheS recombinant plasmid. The primers used for the gene integration processes are included in Supplementary Table S7. All plasmids utilized for the gene integration processes are listed in Supplementary Table S6.

### Measuring genomic repression of mCherry-XenO target using dTnpB

To measure repression of the mCherry-XenO fusion protein, we transformed our dTnpB + reRNA-containing plasmid (pAT00.40.001-dTnpB + 20-reRNA1-mCherry) into competent cells of the XenO strain, according to the *D. radiodurans* competent cell preparation and transformation protocols. Single colonies were grown in a shaking incubator at 32 °C for 24 hours in 5 mL of TGY supplemented with chloramphenicol and kanamycin to ensure plasmid retention and select against any potential R1 contamination. After 24 hours of growth, 250 µL of saturated culture was added to

5 mL of fresh TGY supplemented with chloramphenicol, these cultures were grown for 10 minutes in a shaking incubator at 32 °C. Then to induce dTnpB expression, 5 µL of 1M IPTG, or 2.5 µL of 100 µM aTc were added and the samples were returned to the 32 °C shaking incubator. Periodically, fluorescence of the induced and uninduced samples were measured according to above protocols. Final fluorescent measurements were taken 24 hours post-induction.

### Imaging of bacterial strains

Microscopy was performed via confocal microscopy using a Fluoview 1000 confocal microscope on an Olympus IX81 microscope base.The 543 nm laser line was used to observe mCherry fluorescence. The Fluoview settings for observation were as follows, Gain: 1500 (1.5x), Offset: 2000 (20%), PMT Voltage: 664-880.

## Acknowledgements

We would like to thank Gina Partipilo, from the lab of Dr. Benjamin Keitz for her helpful discussion and insights on approaches for developing a protocol to synthesize the RBS Library, as well as providing support and guidance in generating the protocols to measure fluorescence of the *D. radiodurans* cultures. We would also like to thank Dr. Philip Sweet and other members of the Contreras Lab for helpful conversations and discussions during the ideation of the project, and drafting of the manuscript. We’d also like to thank Dr. Vernita Gordon, and Dr. Brandon Niese for their contributions in imaging the XenO strain. Lastly, we would like to thank Dr. Christian Bök for providing us with the inspiration to generate the XenO strain. Further reading regarding the Xenotext (XenO) and its inception are contained within THE XENOTEXT (BOOK 1) [ISBN: 9781552453216] and THE XENOTEXT (BOOK 2) [ISBN: 9781552454985].

