## Supplementary Materials for "Establishing a standardized genetic toolkit for the radiation-resistant extremophile *Deinococcus radiodurans*"

**Table of Contents**

**Supplementary Methods**

**Supplementary Tables**

Table S1 – Screened Promoter Sequences

Table S2 – Regulator Sequences

Table S3 – Fluorescent Report Sequences – Original and Codon Optimized for *D. radiodurans*

Table S4 – RBS Library Sequences

Table S5 – XenO Fusion Proteins used in this study

Table S6 – Strains and Plasmids used in this study

Table S7 – Oligonucleotides used in this study

**Supplementary Figures**

Figure S1 – Computationally Predicted RBS strengths vs Relative fluorescence of each RBS design expressing sfGFP

Figure S2 – Determining XenO-mCherry Fusion Candidates

Figure S3 – Designing a minimal base plasmid for expression in *D. radiodurans* and identifying novel origin of replications to expand plasmid versatility

**Supplementary Methods**

These supplementary methods are written in detail, such that other can use these as protocols directly.

***Deinococcus radiodurans* Heat Competent Cell Creation**

**Introduction**

This section details how to create heat-shock competent *Deinococcus radiodurans* cells for uptake of bacterial DNA.

**Regents, Supplies, Materials, and Equipment**

- 15 mL Plastic culture tubes
- TGY Media
- Bucket of ice
- 1M CaCl2
- 30% glycerol
- Autoclaved Eppendorf Tubes (~15 tubes per 5mL of overnight culture)

**Procedure**

Overnight Preparation

1. Grow *Deinococcus radiodurans* in TGY to OD about 0.8~1 (~2.5-3 hours from an OD600 of 0.3)
2. Prepare a bucket of ice, and chill 1M CaCl2 solution, and 30% glycerol solution.
3. To make *D. rad* cells competent: mix 955µL of cells, 45µL of 1M CaCl2 (cold), 500µL of 30% glycerol (cold) in an autoclaved Eppendorf tube. Aliquot 300µL of the competent cell mixture into autoclaved Eppendorf tubes.
4. Freeze the competent cells -80˚C for later usage; *alternatively* – they can be used immediately.

***Deinococcus radiodurans* Heat Competent Cell Transformation**

**Introduction**

This section details how to create heat-shock competent *Deinococcus radiodurans* cells for uptake of bacterial DNA.

**Regents, Supplies, Materials, and Equipment**

- 15 mL Plastic culture tubes
- TGY Media
- Bucket of ice
- 1M CaCl2
- 30% glycerol
- Autoclaved Eppendorf Tubes (~15 tubes per 5mL of overnight culture)

**Procedure**

1. Thaw the competent cells on ice.
2. Add 1~1.5 µg of plasmid DNA into each Eppendorf tube of competent cells.
3. Incubate competent cells on ice for 30 min – 1 hour.
4. Incubate competent cells at 32˚C for 1 hour.
5. Add 800µL of TGY media into the Eppendorf tube and transfer to a test tube andgrow at 30˚C for at least 5 hours; *alternatively* – perform this process overnight.
6. Spin down the cells at 3000rpm for 5 minutes.
7. Gently re-suspend the cells with 150µL of TGY and plate on a TGY plate with the appropriate amount of antibiotic.
   a. For pRad-Gro plasmids, this would be a 1000x dilution from 3.4mg/mL Chloramphenicol stock, or 10,000x dilution from 34mg/mL Chloramphenicol stock
8. Incubate the plated cells at 32˚C for 2-3 days.

***Deinococcus radiodurans* Genomic Deletion & Insertion Protocol**

**Introduction**

This section details how to create heat-shock competent *Deinococcus radiodurans* cells for uptake of bacterial DNA.

**Regents, Supplies, Materials, and Equipment**

15 mL Plastic culture tubes

- TGY Media
- puc19mPhes Prepped Plasmid
- HindIII Enzyme
- LB Media
- NEBuilder HiFi DNA Assembly mix
- Gel Extraction Purification Kit of Choice

**Procedure**

1. Linearize pUC19mPheS by HindIII (R0104 New England Biolabs, Ipswich MA) restriction enzyme digest following the enzyme manufacture’s protocol.
2. PCR amplify the DNA fragments specifically needed for the desired gene knockout/knock-in from *D.rad* genomic DNA.
   a. 1 kb of upstream homology [from gDNA]
   a. Your selectable antibiotic resistance marker (KAN, traditionally)
   a. 1 kb of downstream homology [from gDNA]
   a. (If Insertion) Your desired Gene of Interest.

- Gene of interest is inserted at the in-between coordinates of Chrom-1 following:
  - 1015932
  - 1015933
  - pUC19 Plasmid Example: <https://benchling.com/s/seq-cHNaK5JVHbK4QkOE718D?m=slm-iyWVDkkFDlKdqp0GQkoD>

3. Use a high-fidelity thermostable polymerase (Phusion DNA Polymerase, M0530L New England Biolabs, Ipswich MA) by the manufacture’s protocol. The primers used for PCR need to have extensions that result in 20-25 bp of overlapping homology to the DNA fragment that will be adjacent to it in the plasmid. These requirements are described in the NEBuilder HiFi DNA assembly protocol (E2621 New England Biolabs, Ipswich MA). If desired enter the sequence of the fragments and plasmid backbone into the NEBuilder tool on their web site to obtain primers that incorporate the needed overlaps for assembly.
3. Agarose gel purify all DNA fragments using Qiagen MinElute (28606 Qiagen, Germantown MD) gel extraction mini columns by the manufacturer’s protocol and quantitate the DNA using a Nanodrop spectrophotometer.
   a. Note: High GC content PCR fidelity is improved dramatically using the PCR kit supplied high GC Buffer or simply having DMSO at 3% final in the PCR reaction.
3. The 4-5 DNA fragments (linear plasmid, [inserted gene], upstream homology, resistance marker, and downstream homology) are assembled using NEBuilder HiFi DNA mix (E2621 New England Biolabs, Ipswich MA) by the manufacturer’s protocol.
4. The assembly mix is used to transform chemically competent *E. coli* strains by the manufacturer’s protocol, a high efficiency DH5alpha (*E. coli* MAX Efficiency DH5α, 18258012 Thermo Fisher Scientific/Invitrogen/Life Technologies, Waltham MA) There is a lower efficiency *dam*^-^ *dcm*^-^ strain (C2925H New England Biolabs, Ipswich MA) should initial transformations fail. The *dam*^-^ *dcm*^-^ *E. coli* strain will yield plasmid DNA that is unmethylated. Unmethylated DNA is introduced into *D. radiodurans* at a significantly higher rate than methylated DNA. Thus, any clones obtained from the *dam*^-^ *dcm*^-^ strain can be directly used for *D. radiodurans*. These *E. coli* transformations will be plated on LB agar plates with kanamycin 16 ug/ml (5.000000 Werner BioAgents GmbH, Jena, Germany).
3. The plasmid construction is confirmed in *E. coli* by full sequencing with Plasmidsaurus.

**Mutant Selection for Marked Deletions:**

1. From the transformation, kanamycin resistant individual colonies are inoculated into 5 mL of TGY broth without antibiotics and incubated with aeration overnight at 32°C. This grow out allows time for second-cross homologous recombination and for mutant chromosomes to segregate from wildtype by cell division.
2. The overnight culture is serially diluted 1:10, seven times and spotted on TGY agar with antibiotic and 5 mM 4-chloro-phenylalanine (4-CP) (C6506 Sigma-Aldrich, St. Louis MO). The plates are incubated ~48 hours at 32°C. The 4-CP selects against any cells that have not undergone a second-cross over and still have plasmid mutated PheS gene.
3. Colonies growing on antibiotic and 4-CP are picked, suspended in 20 uL of TGY broth and then checked by colony PCR (see above protocol). Note, since Deinococcus has multiple copies of its genome it may be necessary to restreak to ensure that mutant is homozygous and both a wild-type and mutant version are present .  It may take several rounds (usually five rounds are enough to get clean knockout colonies) of such streaking to bring to homozygosity.  If it proves impossible to purify mutant from wild-type it may be presumptive evidence that the gene (or cis regulatory) element is essential and the mutant can only be maintained if a wild-type copy is also present.
4. If the desired deletion is confirmed with no indication of wildtype sequences remaining, the portion of the suspended colony not used for PCR template is inoculated into 5 mL TGY broth with antibiotic and incubated with aeration for ~48hrs.
5. In a cryovial, 500 uL of the culture is mixed with 500 uL of 50% Glycerol in TGY broth and stored as a glycerol stock at -80°C.
6. A portion of the remaining culture is pelleted and the chromosomal DNA is isolated using QIAamp DNA Mini Kit (51306 Qiagen, Germantown MD) with the gram positive bacteria lysis protocol. 100 ng of chromosomal DNA is used for PCR template in a 50 uL reaction as a final confirmation of a marked deletion free of wildtype sequences. The genomic DNA can also be used for sequencing to confirm incorporation of the intended changes.

**Cre/*lox* Removal of *D. radiodurans* Genomic Integration Resistance Marker:**

1. *D. radiodurans* deletion mutants with *lox* site flanked resistance markers are transformed by the above-mentioned transformation protocol with pDeinoCre and plated on TGY agar with 3 ug/mL chloramphenicol.
2. After incubation at 32^o^C for ~48hrs, individual colonies are picked and suspended in 200 μL of TGY broth. These cell suspensions are serially diluted 1:10 seven times. As a check for the efficiency with which Cre activity led to loss of the resistance marker via recombination at the *lox* sites, 5 μL of each dilution are spotted on TGY agar, TGY agar with 3 ug/mL chloramphenicol, and TGY agar with antibiotic for the resistance marker being removed. These agar plates are incubated ~48 hours at 32°C and only isolates showing extensive marker removal are further processed.
3. The chloramphenicol resistant suspended colonies are also diluted 1:10,000 into 5 ml TGY broth with no antibiotic and incubated with aeration for ~48 hours at 32°C to allow for the culture to go through several divisions and potentially lose the pDeinoCre plasmid at a low frequency because of uneven segregation at each cell division.
4. These cultures are serially diluted 10-fold seven times, spotted on TGY agar and TGY agar with 5 mM 4-CP, and incubated ~48 hours at 32°C to select for candidates having lost the plasmid.
5. To screen 4-CP resistant colonies for sensitivity to chloramphenicol and the Cre/*lox* removed resistance marker, isolated colonies are picked from the 4-CP plates, suspended in 200 μL TGY broth, serially diluted 10-fold seven times, and 5 uL spotted on TGY agar with 5 mM 4-CP, 3 ug/ml chloramphenicol, and the antibiotic for the marker intended to be removed by site-specific recombination at *lox*.
6. After incubation at 32^o^C for ~48hrs, colonies are picked from the 4-CP plates for isolates that show 4-CP resistance and complete sensitivity to all antibiotics tested. These final isolates are screened by PCR for having the resistance marker removed and a PCR fragment size consistent with gene deletion and the presence of a *lox* scar.

**Miniprepping plasmids DNA from *Deinococcus radiodurans***

**Introduction**

This section details how to purify plasmids out of *Deinococcus radiodurans* utilizing the Qiagen Spin Miniprep plasmid kit, with amendments from other previously generated Contreras Lab protocols, which will be highlighted below.

**Regents, Supplies, Materials, and Equipment**

- 15 mL Plastic culture tubes
- TGY Media
- 3.4 mg/mL chloramphenicol antibiotic
- Qiagen Spin Miniprep Kit
- Chicken Egg Lysozyme (Muramidase)
- Heat Block
- Shaking incubator

**Procedure**

Overnight Preparation

1. Add 5 mL of TGY Media to 15 mL plastic overnight tubes
1. Add 5 µL of 3.4 mg/mL (1000x) chloramphenicol to the TGY
2. Add single colony of *D. rad.* to TGY media using a pipet tip
3. Grow at 32 °C for 12-24 hours or until cultures are turbid
4. Spin 1.5 mL of cultures in the centrifuge at 5000g for 5 minutes
5. Decant supernatant and repeat step 5
6. Resuspend pellet in supernatant from step 6
7. Heat cultures for 10 minutes at 95 °C in the shaking heat block
8. Take off the heat block to cool for minute
9. Spin down cells at 5000g for 5 minutes
10. While the cells centrifuge, aliquot 3 mL of Buffer P1 to a fresh 15 mL Falcon Tube and dissolve 18 mg of Chicken Egg Lysozyme into the solution, such that the final concentration is ~6 mg/mL

Note 1: Always prepare this solution fresh to maximize lytic activity

Note 2: You may have to vortex the solution to get the Lysozyme to fully dissolve

11. After the cells finish centrifuging, aspirate the supernatant as much as possible
12. Resuspend the pellet in 250 µL of the P1 + Lysozyme Buffer
13. Incubate the samples in the 37 °C shaking incubator for 15 minutes

Note 3: Incubation time can be extended up to 60 minutes, if more lysis is needed

14. Proceed with the Qiagen Spin MiniPrep kit instructions from Step 3 (P2 Buffer addition step)

Note 4: Allow the P2 Lysis step to proceed for at least 3 minutes to maximize alkaline lysis

Note 5: When eluting at the end of the protocol, use 25 µL of DI water instead of the suggested 50 µL EB Buffer provided with the kit

**Codon optimization for optimal gene expression in the organism *Deinococcus radiodurans***

**Introduction**

This section details how to perform codon optimization utilizing a novel script from this manuscript in the organism *D. radiodurans*. Examples of codon optimization are contained in Supplementary Table S3, with results from optimization on fluorescent reporters in Figure 3.

**Procedure**

Code contained below as generated by authors, copy and run in your preferred Python IDE. Use the pop-up GUI window to optimize CDS for expression in *D. radiodurans*:

import tkinter as tk

from tkinter import scrolledtext

from tkinter import messagebox

\# Codon usage table sourced from http://lowelab.ucsc.edu/GtRNAdb/Dein_radi/Dein_radi-summary-codon.html

codon_usage_table = {

'A': {'GCT': 0.88, 'GCC': 5.97, 'GCG': 4.74, 'GCA': 0.63},

'G': {'GGT': 0.73, 'GGC': 5.85, 'GGG': 2.08, 'GGA': 0.54},

'P': {'CCT': 0.52, 'CCC': 2.74, 'CCG': 2.5, 'CCA': 0.3},

'T': {'ACT': 0.36, 'ACC': 3.47, 'ACG': 1.73, 'ACA': 0.21},

'V': {'GTT': 0.36, 'GTC': 2.31, 'GTG': 4.84, 'GTA': 0.23},

'S': {'TCT': 0.2, 'TCC': 0.76, 'TCG': 1.16, 'TCA': 0.17, 'AGT': 0.43, 'AGC': 2.38},

'R': {'CGT': 0.68, 'CGC': 4.1, 'CGG': 2.06, 'CGA': 0.23, 'AGG': 0.21, 'AGA': 0.11},

'L': {'CTT': 0.49, 'CTC': 3.43, 'CTG': 6.75, 'CTA': 0.13, 'TTG': 0.73, 'TTA': 0.07},

'F': {'TTT': 1.05, 'TTC': 2.07},

'N': {'AAT': 0.35, 'AAC': 2.03},

'K': {'AAG': 1.94, 'AAA': 0.78},

'D': {'GAT': 0.53, 'GAC': 4.52},

'E': {'GAG': 3.22, 'GAA': 2.52},

'H': {'CAT': 0.33, 'CAC': 1.76},

'Q': {'CAG': 3.4, 'CAA': 0.71},

'I': {'ATT': 0.99, 'ATC': 2.19, 'ATA': 0.08},

'M': {'ATG': 1.77},

'Y': {'TAT': 0.3, 'TAC': 1.92},

'C': {'TGT': 0.1, 'TGC': 0.55},

'W': {'TGG': 1.38},

'*': {'TAG': 0.03, 'TAA': 0.08, 'TGA': 0.21},

}

def validate_dna_sequence(sequence):

valid_nucleotides = set("ATGCatgc")

return all(nucleotide in valid_nucleotides for nucleotide in sequence)

def validate_aa_sequence(sequence):

valid_amino_acids = set("ACDEFGHIKLMNPQRSTVWY")

return all(aa in valid_amino_acids for aa in sequence)

def optimize_codons_dna(dna_sequence, codon_usage_table):

if not validate_dna_sequence(dna_sequence):

raise ValueError("Invalid DNA sequence. Please provide a sequence containing only A, T, G, and C.")

optimized_sequence = ""

for i in range(0, len(dna_sequence), 3):

codon = dna_sequence[i:i+3]

amino_acid = translate_codon(codon.upper())

if amino_acid == '*': # If it's a stop codon, just add it without modification

optimized_sequence += codon

elif amino_acid in codon_usage_table:

optimal_codon = max(codon_usage_table[amino_acid], key=codon_usage_table[amino_acid].get)

optimized_sequence += optimal_codon

else:

optimized_sequence += codon

return optimized_sequence

def optimize_codons_aa(aa_sequence, codon_usage_table):

if not validate_aa_sequence(aa_sequence):

raise ValueError("Invalid amino acid sequence. Please provide a valid sequence.")

dna_sequence = translate_aa_to_dna(aa_sequence)

return optimize_codons_dna(dna_sequence, codon_usage_table)

def translate_aa_to_dna(aa_sequence):

\# Convert amino acid sequence to DNA sequence using the optimal codons

dna_sequence = ''

for aa in aa_sequence:

if aa in codon_usage_table:

\# If it's a stop codon, we can simply skip it here

if aa == '*':

dna_sequence += 'TAA' # Or any valid stop codon; we'll use 'TAA' here

else:

optimal_codon = max(codon_usage_table[aa], key=codon_usage_table[aa].get)

dna_sequence += optimal_codon

return dna_sequence

def translate_codon(codon):

translation_table = {

'GCT': 'A', 'GCC': 'A', 'GCA': 'A', 'GCG': 'A',

'GGT': 'G', 'GGC': 'G', 'GGG': 'G', 'GGA': 'G',

'CCT': 'P', 'CCC': 'P', 'CCG': 'P', 'CCA': 'P',

'ACT': 'T', 'ACC': 'T', 'ACG': 'T', 'ACA': 'T',

'GTT': 'V', 'GTC': 'V', 'GTG': 'V', 'GTA': 'V',

'TCT': 'S', 'TCC': 'S', 'TCG': 'S', 'TCA': 'S', 'AGT': 'S', 'AGC': 'S',

'CGT': 'R', 'CGC': 'R', 'CGG': 'R', 'CGA': 'R', 'AGG': 'R', 'AGA': 'R',

'CTT': 'L', 'CTC': 'L', 'CTG': 'L', 'CTA': 'L', 'TTG': 'L', 'TTA': 'L',

'TTT': 'F', 'TTC': 'F',

'AAT': 'N', 'AAC': 'N',

'AAG': 'K', 'AAA': 'K',

'GAT': 'D', 'GAC': 'D',

'GAG': 'E', 'GAA': 'E',

'CAT': 'H', 'CAC': 'H',

'CAG': 'Q', 'CAA': 'Q',

'ATT': 'I', 'ATC': 'I', 'ATA': 'I',

'ATG': 'M',

'TAT': 'Y', 'TAC': 'Y',

'TGT': 'C', 'TGC': 'C',

'TGG': 'W',

'TAG': '*', 'TAA': '*', 'TGA': '*',

}

return translation_table.get(codon, '')

def on_optimize_button_click():

input_sequence = original_sequence_entry.get("1.0", tk.END).strip()

input_type = input_type_var.get()

try:

if input_type == 'Amino Acids':

optimized_sequence = optimize_codons_aa(input_sequence, codon_usage_table)

elif input_type == 'Nucleotides':

optimized_sequence = optimize_codons_dna(input_sequence, codon_usage_table)

optimized_sequence_text.config(state=tk.NORMAL)

optimized_sequence_text.delete("1.0", tk.END)

optimized_sequence_text.insert(tk.END, optimized_sequence)

optimized_sequence_text.config(state=tk.DISABLED)

except ValueError as e:

messagebox.showerror("Error", str(e))

\# GUI setup

app = tk.Tk()

app.title("The D. Rad. Codon Optimizer by The Contreras Lab at UT-Austin")

\# Input type dropdown

tk.Label(app, text="Select Input Type:").pack()

input_type_var = tk.StringVar(value="Nucleotides")

input_type_menu = tk.OptionMenu(app, input_type_var, "Amino Acids", "Nucleotides")

input_type_menu.pack()

\# Input

tk.Label(app, text="Input Original Sequence:").pack()

original_sequence_entry = scrolledtext.ScrolledText(app, wrap=tk.WORD, width=80, height=18)

original_sequence_entry.pack()

\# Optimize button

optimize_button = tk.Button(app, text="Optimize!", command=on_optimize_button_click)

optimize_button.pack()

\# Output

tk.Label(app, text="Your Optimized DNA Sequence:").pack()

optimized_sequence_text = scrolledtext.ScrolledText(app, wrap=tk.WORD, width=80, height=18, state=tk.DISABLED)

optimized_sequence_text.pack()

app.mainloop()

**Supplementary Tables**

**Supplementary Table S1.** Screened Promoter Sequences

| Name | Sequence |
| --- | --- |
| pGroES | gtcagttgacatttttcttatcggcgctctaccatccgtga |
| p1261 | gacctgccctgggagccacgcaccatctctaacgacgccttgatgttgctatcacgccggggctg |
| p1348 | ttgcttgagcgccgcccgctttctttacgcgtcgccaacgtccggctcactggcagctctggatcagcgcgcacgataagtcc |
| p1473 | gaggaggagtctgacttgctccctcatctacagcccctgagacaaaaggaaaaccccgcgccaggcggggctttttattggagccgagggtcggacttgaaccgacgacctactgattacgaatcagttgctctac |
| p2508 | gaggtaattctggattggcccctattctgaaaatcaagccgtccctgaacttaatcaaagctacactctatggcaattcagtattcatctgacacatatcgatctatccgcgactttatgagacaaacattttaactgtgtcagacagtgtttttgatcacctattgttgccaccaagtcaggctgaccaacctgacctcaaactgacgaaaaaagccgacttttgcgctttagacgacaaaaagtcaccttgacagagctattatctcatctctataattcgcgccgggatctc |
| pAmyE | caccagaaggcgacgatggaagtggcgaccgcgctgtggggcgtgaccaccgaccagaccggggcgagccgggtactgagcatcaagcaggagcgggaataagggcacttcgcaggctgg |
| pClpB | tatgccgtcacctggctccccgacgcgcgccgcgcggatgtggaacgggtgcggaaaattccgggtggactgtcacagacacagggcgaactggcgggccttaagctaaagccgacgcgcctgaatgcctccgcagaagtacggtttcaacctctgaagaacgaaccagcatttgaggagctgtttctctcgtcaccagcgcgagaattggtgcgtatctggccgtttttt |
| pDnaK | gcttttcttgatgttgccagcaaacttgagtctaatacactcaagtctattgactctgggggattgtagtcgtatgatacttgcagttcaag |
| pGroEL | ggctccgagtcaggttctgagcctgttcgtttcctgtttttcttcctcatttcacttttcaaggagcaatcaca |
| pKatA | cctgcgccttcttcttgaacccactgcacagaacccactggacagaacccactgcacagccgccgcgagggcctgagggccatggagaccgagggccctggacattgagaatgattctcaatatggtgcagggagcttcggg |
| pLexA | gagcgctacgccttcatcgccagttatccgggcgccgcagaggagacttttctggtgccttcttccgaactcacggaagctgcggccacccagttgtcagtcagggacgacggtcagccctattcttttccgttcaa |
| pRecA | gccggttgccgtaaagctgcggcagaacaccagaaaaaaagagcgcgaagaacagcggcaacaccgcaacggccagccagggccagcgccgaacggtccagcaagtcagccccgccgcgcaggccaacagggccacggtctgaaggataaaagcctccatcacagcggcgtgaccgggcacccgcaccgccgacacctgcgcctcgtgcacgaaaccgctcaacgaataagccggcaaggcgagcgccagccccgccgccaccaagctccagccgcgcctgcgggtgcccagccagccgagcgcgaaccccaggagcagcggcacaaagagcagcgccaggaaaacgagaaacaccagcatgatcgcaaggtaggctgctccgcgcactggggcgtccgccgaaaggtgcagcgcgcctcaccttcccttgacccatcccattcagggccgcaccctgtagacactggttacgtc |
| pRecQ | gaagcctccagatcaaaggcgcacgaagcaggcctgccccaaaggaaggcgggcctgctgggttgcgtgacagccgggacgtaaataatggccccagggtaggggcg |
| pRplL | gcagcgagagcgcctaaagcgcccgcttctccttccctgcatttctctttccctaaaatccacttctggaggacatacac |
| pRpmB | cggccacatgacgacccgatgccggtgcgggccacagcagggagtggcgaaggctccgcgccccaggtctatctttcgggctcgccagtaggttaagatagggtttggtgtcgc |
| pTufB | ccaacgaagaaatgcggcccggataagagggccgcatttcgattttgaccgagcatggccctacgccagcccgcgcacttttgatatagtgttcagg |
| pTac (IPTG) | TGTTGACAATTAATCATCGGCTCGTATAATGTGTGGAATTGTGAGCGCTCACAATT |
| p3B5B (DHBA) | TTTTGTTCGATTATCGAACAAATTATTGAAATATCGAACAAAACCTCTAAACTACTGTGGCACTGAATCAAAAAATTATAAACCATGATCAGA |
| pCin (OHC14) | CCCTTTGTGCGTCCAAACGGACGCACGGCGCTCTAAAGCGGGTCGCGATCTTTCAGATTCGCTCCTCGCGCTTTCAGTCTTTGTTTTGGCGCATGTCGTTATCGCAAAACCGCTGCACACTTTTGCGCGACATGCTCTGATCCCCCTCATCTGGGGGGGCCTATCTGAGGGAATTTCCGATCCGGCTCGCCTGAACCATTCTGCTTTCCACGAACTTGAAAACGCT |
| pSalTTC (Sal) | GGGGCCTCGCTTGGGTTATTGCTGGTGCCCGGCCGGGCGCAATATTCATGTTGATGATTTATTATATATCGAGTGGTGTATTTATTTATATTGTTTGCTCCGTTACCGTTATTAAC |
| pCymR (CA) | AACAAACAGACAATCTGGTCTGTTTGTATTATGGAAAATTTTTCTGTATAATAGATTCAACAAACAGACAATCTGGTCTGTTTGTATTAT |
| pTet_Star (aTc) | TTTTCAGCAGGACGCACTGACCTCCCTATCAGTGATAGAGATTGACATCCCTATCAGTGATAGAGATACTGAGCAC |
| pBetI (Cho) | AGCGCGGGTGAGAGGGATTCGTTACCAATAGACAATTGATTGGACGTTCAATATAATGCTAGC |
| pMph (Ery) | GGATTGAATATAACCGACGTGACTGTTACATTTAGGTGGCTAAACCCGTCAA |
| pTtg (Nar) | CACCCAGCAGTATTTACAAACAACCATGAATGTAAGTATATTCCTTAGCAA |
| pVanCC (Va) | ATTGGATCCAATTGACAGCTAGCTCAGTCCTAGGTACCATTGGATCCAAT |
| pBad (Ara) | AGAAACCAATTGTCCATATTGCATCAGACATTGCCGTCACTGCGTCTTTTACTGGCTCTTCTCGCTAACCAAACCGGTAACCCCGCTTATTAAAAGCATTCTGTAACAAAGCGGGACCAAAGCCATGACAAAAACGCGTAACAAAAGTGTCTATAATCACGGCAGAAAAGTCCACATTGATTATTTGCACGGCGTCACACTTTGCTATGCCATAGCATTTTTATCCATAAGATTAGCGGATCCTACCTGACGCTTTTTATCGCAACTCTCTACTGTTTCTCCATACCCG |
| PacU (Acr) | CGCTAGCAAGTAAGGCCGACGCTTCACAACCGCACTTGATTTAATAGACCATACCGTCTATTATTTCTGGCCAT |
| pLuxB (Oc6) | ACCTGTAGGATCGTACAGGTTTACGCAAGAAAATGGTTTGTTACAGTCGAATAAA |
| pLacI | GCGGCGCGCCATCGAATGGCGCAAAACCTTTCGCGGTATGGCATGATAGCGCCC |
| pLacIQ | GCGGCGCGCCATCGAATGGTGCAAAACCTTTCGCGGTATGGCATGATAGCGCCC |
| araC | ATGGCTGAAGCGCAAAATGATCCCCTGCTGCCGGGATACTCGTTTAATGCCCATCTGGTGGCGGGTTTAACGCCGATTGAGGCCAACGGTTATCTCGATTTTTTTATCGACCGACCGCTGGGAATGAAAGGTTATATTCTCAATCTCACCATTCGCGGTCAGGGGGTGGTGAAAAATCAGGGACGAGAATTTGTTTGCCGACCGGGTGATATTTTGCTGTTCCCGCCAGGAGAGATTCATCACTACGGCCGTCATCCGGAGGCTCGCGAATGGTATCACCAGTGGGTTTACTTTCGTCCGCGCGCCTACTGGCATGAATGGCTTAACTGGCCGTCAATATTTGCCAATACGGGGTTCTTTCGCCCGGATGAAGCGCACCAGCCGCATTTCAGCGACTTTTTTGGGCAAATCATTAACGCCGGGCAAGGGGAAGGGCGCTATTCGGAGCTGCTGGCGATAAATCTGCTTGAGCAATTGTTACTGCGGCGCATGCTAGCGATTAACGGATCGCTCCATCCACCGATGGATAATCGGGTACGCGAGGCTTGTCAGTACATCAGCGATCACCTGGCAGACAGCAATTTTGATATCGCCAGCGTCGCACAGCATGTTTGCTTGTCGCCGTCGCGTCTGTCACATCTTTTCCGCCAGCAGTTAGGGATTAGCGTCTTAAGCTGGCGCGAGGACCAACGTATCAGCCAGGCGAAGCTGCTTTTGAGCACCACCCGGATGCCTATCGCCACCGTCGGTCGCAATGTTGGTTTTGACGATCAACTCTATTTCTCGCGGGTATTTAAAAAATGCACCGGGGCCAGCCCGAGCGAGTTCCGTGCCGGTTTGGAAGAAAAAGTGAATGATGTAGCCGTCAAGTTGTCATGA |

**Supplementary Table S2.** Regulator Sequences

| **Name** | **Sequence (5'-3')** |
| --- | --- |
| lacI | ATGAAACCAGTAACGTTATACGATGTCGCAGAGTATGCCGGTGTCTCTTATATGACCGTTTCCCGCGTGGTGAACCAGGCCAGCCACGTTTCTGCGAAAACGCGGGAAAAAGTGGAAGCGGCGATGGTGGAGCTGAATTACATTCCCAACCGCGTGGCACAACAACTGGCGGGCAAACAGTCGTTGCTGATTGGCGTTGCCACCTCCAGTCTGGCCCTGCACGCGCCGTCGCAAATTGTCGCGGCGATTAAATCTCGCGCCGATCAACTGGGTGCCAGCGTGGTGGTGTCGATGGTAGAACGAAGCGGCGTCGAAGCCTGTAAAGCGGCGGTGCACAATCTTCTCGCGCAACGCGTCAGTGGGCTGATCATTAACTATCCGCTGGATGACCAGGATGCCATTGCTGTGGAAGCTGCCTGCACTAATGTTCCGGCGTTATTTCTTGATGTCTCTGACCAGACACCCATCAACAGTATTATTTACTCCCATGAGGACGGTACGCGACTGGGCGTGGAGCATCTGGTCGCATTGGGTCACCAGCAAATCGCGCTGTTAGCGGGCCCATTAAGTTCTGTCTCGGCGCGTCTGCGTCTGGCTGGCTGGCATAAATATCTCACTCGCAATCAAATTCAGCCGATAGCGGAACGGGAAGGCGACTGGAGTGCCATGTCCGGTTTTCAACAAACCATGCAAATGCTGAATGAGGGCATCGTTCCCACTGCGATGCTGGTTGCCAACGATCAGATGGCGCTGGGCGCAATGCGCGCCATTACCGAGTCCGGGCTGCGCGTTGGTGCGGATATCTCGGTAGTGGGATACGACGATACCGAAGATAGCTCATGTTATATCCCGCCGTTAACCACCATCAAACAGGATTTTCGCCTGCTGGGGCAAACCAGCGTGGACCGCTTGCTGCAACTCTCTCAGGGCCAGGCGGTGAAGGGCAATCAGCTGTTGCCAGTCTCACTGGTGAAAAGAAAAACCACCCTGGCGCCCAATACGCAAACCGCCTCTCCCCGCGCGTTGGCCGATTCATTAATGCAGCTGGCACGACAGGTTTCCCGACTGGAAAGCGGGCAGTGA |
| pcaU | ATGTGGTCGAACATGGATGACAAGAAAGTGAAAGAGGAGAATATTCTGCACAATTCCACCAACAAGAAGATCATCCGCCACGAAGATTTTGTAGCCGGCATTAGCAAAGGGATGGCGATTCTGGATTCGTTTGGTACAGATCGTCATCGCCTCAATATCACCATGGCCGCAGAGAAAACCGGTATGACACGTGCAGCAGCTCGTCGCCACCTGCTTACTCTGGAGTATCTGGGCTATCTGGAAAGTGACGGCCACTACTTCTACTTAACTCCCAAAATCCTGAAATTCAGTGGTTCATATTTGGGTGGTGCTCAATTGCCGAAAATTTCCCAACCACTGTTGAACTTGCTTACGACCCAGACCAGCCTGATTTACAGCGTGATGGTGTTGGATGGCTATGAAGCCATTACCATTGCGCGTTCTGCCGCTCATCAGCAAACCGACCGCGTTAACCCGTATGGTTTACATCTCGGGAATCGCTTACCAGCGCATACAACGTCAGCGGGCAAAATCCTGTTAGCGTATTTGGATGACCATGCCCAGCAAGAGTGGCTCAATCAGTACCCTCTGCAACGGCTCACGAAATACACGTATACCAACCACATCGACTTTCTGCGCCTTTTGAGTGAAATCAAGGAACAGGGTTGGTGCTATAGTTCGGAAGAACACGAACTGGGAGTACACGCCCTTGCGGTTCCGATTTACGGACAGCAGTCTCGCGTCGTAGCGGCACTGAACATTGTCAGCCCGACAATGCGGACCACGAAAGAATACCTGATTCAGCATATTCTGCCGTTACTGCAAGAAACTGCGCGTGAATTGCGCAATATCCTGTAATGA |
| cinR | ATGATTGAGAATACCTATAGCGAAAAGTTCGAGTCCGCGTTCGAACAGATCAAAGCGGCGGCCAACGTGGATGCCGCCATCCGTATTCTCCAGGCGGAATATAACCTCGATTTCGTCACCTACCATCTCGCCCAGACAATCGCGAGCAAGATCGATTCGCCCTTCGTGCGCACCACCTATCCGGATGCCTGGGTTTCCCGTTACCTCCTCAACTGCTATGTGAAGGTCGATCCGATCATCAAGCAGGGCTTCGAACGCCAGCTGCCCTTCGACTGGAGCGAGGTCGAACCGACGCCGGAGGCCTATGCCATGCTGGTCGACGCCCAGAAACACGGCATCGATGACAATGGCTACTCCATCCCCGTCGCCGACAAGGCGCAGCGCCGCGCCCTGCTGTCGCTGAATGCCCATATACCGGCCGACGAATGGACCGAGCTCGTGCGCCGCTGCCGCAATGAGTGGATCGAGATCGCCCATCTGATCCACCGCAAGGCCGTATATGAGCTGCATGGCGAAAACGATCCGGTGCCGGCATTGTCGCCGCGCGAGATCGAGTGTCTGCACTGGACCGCCCTCGGCAAGGATTACAAGGATATTTCGGTCATCCTGGGCATATCAGAGCATACCACACGCGATTACCTGAAAACCGCCCGCTTCAGGCTCGGCTGCACCACGATCTCGGCCGCCGCGTCGCGGGCTGTTCAATTGCGCATCATCAATCCCTATAGGATCCGCATGACGCGACGTAATTGGTAA |
| nahR | ATGGAACTGCGTGACCTGGATTTAAACCTGCTGGTGGTGTTCAACCAGTTGCTGGTCGACAGACGCGTCTCTGTCACTGCGGAGAACCTGGGCCTGACCCAGCCTGCCGTGAGCAATGCGCTGAAACGCCTGCGCACCTCGCTACAGGACCCACTCTTCGTGCGCACACATCAGGGAATGGAACCCACACCCTATGCCGCGCATCTGGCCGAGCACGTCACTTCGGCCATGCACGCACTGCGCAACGCCCTACAGCACCATGAAAGCTTCGATCCGCTGACCAGCGAGCGTACCTTCACCCTGGCCATGACCGACATTGGCGAGATCTACTTCATGCCGCGGCTGATGGATGCGCTGGCTCACCAGGCCCCCAATTGCGTGATCAGTACGGTGCGCGACAGTTCGATGAGCCTGATGCAGGCCTTGCAGAACGGAACCGTGGACTTGGCCGTGGGCCTGCTTCCCAATCTGCAAACTGGCTTCTTTCAGCGCCGGCTGCTCCAGAATCACTACGTGTGCCTATGTCGCAAGGACCATCCAGTCACCCGCGAACCCCTGACTCTGGAGCGCTTCTGTTCCTACGGCCACGTGCGTGTCATCGCCGCTGGCACCGGCCACGGCGAGGTGGACACGTACATGACACGGGTCGGCATCCGGCGCGACATCCGTCTGGAAGTGCCGCACTTCGCCGCCGTTGGCCACATCCTCCAGCGCACCGATCTGCTCGCCACTGTGCCGATATGTTTAGCCGACTGCTGCGTAGAGCCCTTCGGCCTAAGCGCCTTGCCGCACCCAGTCGTCTTGCCTGAAATAGCCATCAACATGTTCTGGCATGCGAAGTACCACAAGGACCTAGCCAATATTTGGTTGCGGCAACTGATGTTTGACCTGTTTACGGATTGA |
| cymR | ATGAGCCCGAAACGTCGTACCCAGGCAGAACGTGCAATGGAAACCCAGGGTAAACTGATTGCAGCAGCACTGGGTGTTCTGCGTGAAAAAGGTTATGCAGGTTTTCGTATTGCAGATGTTCCGGGTGCAGCCGGTGTTAGCCGTGGTGCACAGAGCCATCATTTTCCGACCAAACTGGAACTGCTGCTGGCAACCTTTGAATGGCTGTATGAGCAGATTACCGAACGTAGCCGTGCACGTCTGGCAAAACTGAAACCGGAAGATGATGTTATTCAGCAGATGCTGGATGATGCAGCAGAATTTTTTCTGGATGATGATTTTAGCATCGGCCTGGATCTGATTGTTGCAGCAGATCGTGATCCGGCACTGCGTGAAGGTATTCAGCGTACCGTTGAACGTAATCGTTTTGTTGTTGAAGATATGTGGCTGGGTGTGCTGGTGAGCCGTGGTCTGAGCCGTGATGATGCCGAAGATATTCTGTGGCTGATTTTTAACAGCGTTCGTGGTCTGGTAGTTCGTAGCCTGTGGCAGAAAGATAAAGAACGTTTTGAACGTGTGCGTAATAGCACCCTGGAAATTGCACGTGAACGTTATGCAAAATTCAAACGTTGA |
| tetR | ATGTCCAGATTAGATAAAAGTAAAGTGATTAACAGCGCATTAGAGCTGCTTAATGAGGTCGGAATCGAAGGTTTAACAACCCGTAAACTCGCCCAGAAGCTAGGTGTAGAGCAGCCTACATTGTATTGGCATGTAAAAAATAAGCGGGCTTTGCTCGACGCCTTAGCCATTGAGATGTTAGATAGGCACCATACTCACTTTTGCCCTTTAGAAGGGGAAAGCTGGCAAGATTTTTTACGTAATAACGCTAAAAGTTTTAGATGTGCTTTACTAAGTCATCGCGATGGAGCAAAAGTACATTTAGGTACACGGCCTACAGAAAAACAGTATGAAACTCTCGAAAATCAATTAGCCTTTTTATGCCAACAAGGTTTTTCACTAGAGAATGCATTATATGCACTCAGCGCTGTGGGGCATTTTACTTTAGGTTGCGTATTGGAAGATCAAGAGCATCAAGTCGCTAAAGAAGAAAGGGAAACACCTACTACTGATAGTATGCCGCCATTATTACGACAAGCTATCGAATTATTTGATCACCAAGGTGCAGAGCCAGCCTTCTTATTCGGCCTTGAATTGATCATATGCGGATTAGAAAAACAACTTAAATGTGAAAGTGGGTCCTGA |
| betI | ATGCCGAAACTGGGTATGCAGAGCATTCGTCGTCGTCAGCTGATTGATGCAACCCTGGAAGCAATTAATGAAGTTGGTATGCATGATGCAACCATTGCACAGATTGCACGTCGTGCCGGTGTTAGCACCGGTATTATTAGCCATTATTTCCGCGATAAAAACGGTCTACTGGAAGCAACCATGCGTGATATTACCAGCCAGCTGCGTGATGCAGTTCTGAATCGTCTGCATGCACTGCCGCAGGGTAGCGCAGAACAGCGTCTGCAGGCAATTGTTGGTGGTAATTTTGATGAAACCCAGGTTAGCAGCGCAGCAATGAAAGCATGGCTGGCATTTTGGGCAATCAGCATGCATCAGCCGATGCTGTATCGTCTGCAGCAGGTTAGCAGTCGTCGTCTGCTGAGCAATCTGGTTAGCGAATTTCGTCGTGAACTGCCTCGTGAACAGGCACAAGAGGCAGGTTATGGTCTGGCAGCACTGATTGATGGTCTGTGGCTGCGTGCAGCACTGAGCGGTAAACCGCTGGATAAAACCCGTGCAAATAGCCTGACCCGTCATTTTATCACCCAGCATCTGCCGACCGATTGA |
| mphR | ATGCCTCGTCCGAAACTGAAAAGTGATGATGAAGTTCTGGAAGCAGCAACCGTTGTTCTGAAACGTTGTGGTCCGATTGAATTTACCCTGAGCGGTGTTGCAAAAGAAGTTGGTCTGAGTCGCGCAGCACTGATTCAGCGTTTTACCAATCGTGATACCCTGCTGGTTCGTATGATGGAACGTGGTGTTGAACAGGTTCGTCATTATCTGAATGCAATTCCGATTGGTGCAGGTCCGCAGGGTCTGTGGGAATTTCTGCAGGTTCTGGTTCGTAGCATGGATACCCGTAATGATTTCAGCGTGAACTATCTGATCAGCTGGTATGAACTGCAGGTTCCGGAACTGCGTACCCTGGCAATTCAGCGTAATCGTGCAGTTGTTGAAGGTATTCGTAAACGTCTGCCTCCGGGTGCACCGGCAGCAGCAGAACTGCTGCTGCATAGCGTTATTGCCGGTGCAACCATGCAGTGGGCAGTTGATCCGGATGGTGAACTGGCAGATCATGTTCTGGCACAGATTGCAGCAATTCTGTGTCTGATGTTTCCGGAACATGATGATTTTCAGCTGCTGCAGGCACATGCATAA |
| ttgR | ATGGTTCGTCGTACCAAAGAAGAGGCACAAGAAACCCGTGCACAGATTATTGAAGCAGCAGAACGTGCATTCTATAAACGTGGTGTTGCACGTACCACCCTGGCAGATATTGCAGAACTGGCAGGCGTTACCCGTGGTGCAATTTATTGGCATTTTAACAACAAAGCCGAACTGGTTCAGGCACTGCTGGATAGCCTGCATGAAACCCATGATCATCTGGCACGTGCAAGCGAAAGCGAAGATGAAGTTGATCCGCTGGGTTGTATGCGTAAACTGCTGCTGCAGGTTTTTAATGAACTGGTTCTGGATGCACGTACCCGTCGTATTAATGAAATCCTGCATCACAAATGCGAGTTCACCGATGATATGTGTGAAATTCGTCAGCAGCATCAGAGCGCAGTTCTGGATTGTCATAAAGGTATTACCCTGACACTGGCAAATGTAGTTCGTCGTGGTCAGCTGCCTGGTGAACTGGATGCAGAACGTGCCGCAGTTGCAATGTTTGCCTATGTTGATGGTCTGATTCGTCGTTGGCTGCTGCTGCCGGATAGCGTTGATCTGCTGGGTGATGTTGAAAAATGGGTTGATACCGGTCTGGATATGCTGCGTCTGAGTCCGGCACTGCGTAAATAATGA |
| vanR | ATGGACATGCCTCGTATTAAACCGGGTCAGCGTGTTATGATGGCACTGCGTAAAATGATTGCAAGCGGTGAAATCAAAAGTGGTGAACGTATTGCAGAAATTCCGACCGCAGCAGCACTGGGTGTTAGCCGTATGCCGGTTCGTATCGCACTGCGTTCACTGGAACAAGAAGGTCTGGTTGTTCGTCTGGGTGCACGTGGTTATGCAGCCCGTGGTGTTAGCAGCGATCAGATTCGTGATGCAATTGAAGTTCGTGGTGTTCTGGAAGGTTTTGCAGCACGTCGTCTGGCAGAACGTGGTATGACCGCAGAAACCCATGCACGTTTTGTTGTACTGATTGCAGAAGGTGAAGCACTGTTTGCAGCCGGTCGCCTGAATGGTGAAGATCTGGATCGTTATGCCGCATATAATCAGGCATTTCATGATACCCTGGTTAGCGCAGCAGGTAATGGTGCAGTTGAAAGCGCACTGGCACGTAATGGTTTTGAACCGTTTGCAGCAGCCGGTGCACTGGCCCTGGATCTGATGGACCTGTCTGCCGAATATGAACATCTGCTGGCAGCACATCGTCAGCATCAGGCAGTTCTGGATGCAGTTAGCTGTGGTGATGCCGAAGGTGCAGAACGTATTATGCGTGATCATGCACTGGCAGCAATTCGTAATGCAAAAGTTTTTGAAGCAGCAGCAAGCGCAGGCGCACCGCTGGGTGCAGCATGGTCAATTCGTGCAGATTGA |
| araE | ATGGTTACTATCAATACGGAATCTGCTTTAACGCCACGTTCTTTGCGGGATACGCGGCGTATGAATATGTTTGTTTCGGTAGCTGCTGCGGTCGCAGGATTGTTATTTGGTCTTGATATCGGCGTAATCGCCGGAGCGTTGCCGTTCATTACCGATCACTTTGTGCTGACCAGTCGTTTGCAGGAATGGGTGGTTAGTAGCATGATGCTCGGTGCAGCAATTGGTGCGCTGTTTAATGGTTGGCTGTCGTTCCGCCTGGGGCGTAAATACAGCCTGATGGCGGGGGCCATCCTGTTTGTACTCGGTTCTATAGGGTCCGCTTTTGCGACCAGCGTAGAGATGTTAATCGCCGCTCGTGTGGTGCTGGGCATTGCTGTCGGGATCGCGTCTTACACCGCTCCTCTGTATCTTTCTGAAATGGCAAGTGAAAACGTTCGCGGTAAGATGATCAGTATGTACCAGTTGATGGTCACACTCGGCATCGTGCTGGCGTTTTTATCCGATACAGCGTTCAGTTATAGCGGTAACTGGCGCGCAATGTTGGGGGTTCTTGCTTTACCAGCAGTTCTGCTGATTATTCTGGTAGTATTTCTGCCAAATAGCCCGCGCTGGCTGGCGGAAAAGGGGCGTCATATTGAGGCGGAAGAAGTATTGCGTATGCTGCGCGATACGTCGGAAAAAGCGCGAGAAGAACTCAACGAAATTCGTGAAAGCCTGAAGTTAAAACAGGGCGGTTGGGCACTGTTTAAGATCAACCGTAACGTCCGTCGTGCTGTGTTTCTCGGTATGTTGTTGCAGGCGATGCAGCAGTTTACCGGTATGAACATCATCATGTACTACGCGCCGCGTATCTTCAAAATGGCGGGCTTTACGACCACAGAACAACAGATGATTGCGACTCTGGTCGTAGGGCTGACCTTTATGTTCGCCACCTTTATTGCGGTGTTTACGGTAGATAAAGCAGGGCGTAAACCGGCTCTGAAAATTGGTTTCAGCGTGATGGCGTTAGGCACTCTGGTGCTGGGCTATTGCCTGATGCAGTTTGATAACGGTACGGCTTCCAGTGGCTTGTCCTGGCTCTCTGTTGGCATGACGATGATGTGTATTGCCGGTTATGCGATGAGCGCCGCGCCAGTGGTGTGGATCCTGTGCTCTGAAATTCAGCCGCTGAAATGCCGCGATTTCGGTATTACCTGTTCGACCACCACGAACTGGGTGTCGAATATGATTATCGGCGCGACCTTCCTGACACTGCTTGATAGCATTGGCGCTGCCGGTACGTTCTGGCTCTACACTGCGCTGAACATTGCGTTTGTGGGCATTACTTTCTGGCTCATTCCGGAAACCAAAAATGTCACGCTGGAACATATCGAACGCAAACTGATGGCAGGCGAGAAGTTGAGAAATATCGGCGTCTGA |
| rhiR | ATGCCGCTGACCGACACCCCGCCGTCTGTTCCGCAGAAACCGCGTCGTGGTCGTCCGCGTGGTGCTCCGGACGCTTCTCTTGCTCACCAGTCTCTGATCCGTGCTGGTCTGGAACACCTGACCGAAAAAGGTTACTCTTCGGTTGGTGTTGACGAAATCCTGAAAGCTGCTCGTGTTCCGAAAGGTTCTTTCTACCACTACTTCCGTAACAAAGCTGACTTCGGTCTGGCTCTGATCAAAGCTTACGACACCTACTTCGCTCGTCTCCTCGACCAGGCGTTCCTGGACGGTTCGCTGGCTCCGCTGGCTCGTCTGCGTCTGTTCACCCGTATGGCTGAAGAAGGTATGGCTCGTCACGGTTTCCGTCGTGGTTGCCTGGTTGGTAACCTGGGTCAGGAAATGGGCGCTCTGCCGGACGACTTCCGTGCTGCTCTGATCGGTGTTCTGGAAACCTGGCAACATCGTACCGCTCAGCTGTTCCGTGAAGCTCAGGCTTGCGGTGAACTGTCTGCTGACCATGACCCGGACGCTCTGGCTGAAGCTTTCTGGATCGGATGGGAAGGTGCTATCCTGCGTGCTAAACTGGAACTGCGTCCGGACCCGATGCACTCTTTCACCCGTACCTTCGGTCGTCACTTCGTTACCCGTACCCAGGAATAATGA |
| luxR | ATGAAAAACATAAATGCCGACGACACATACAGAATAATTAATAAAATTAAAGCTTGTAGAAGCAATAATGATATTAATCAATGCTTATCTGATATGACTAAAATGGTACATTGTGAATATTATTTACTCGCGATCATTTATCCTCATTCTATGGTTAAATCTGATATTTCAATCCTAGATAATTACCCTAAAAAATGGAGGCAATATTATGATGACGCTAATTTAATAAAATATGATCCTATAGTAGATTATTCTAACTCCAATCATTCACCAATTAATTGGAATATATTTGAAAACAATGCTGTAAATAAAAAATCTCCAAATGTAATTAAAGAAGCGAAAACATCAGGTCTTATCACTGGGTTTAGTTTCCCTATTCATACGGCTAACAATGGCTTCGGAATGCTTAGTTTTGCACATTCAGAAAAAGACAACTATATAGATAGTTTATTTTTACATGCGTGTATGAACATACCATTAATTGTTCCTTCTCTAGTTGATAATTATCGAAAAATAAATATAGCAAATAATAAATCAAACAACGATTTAACCAAAAGAGAAAAAGAATGTTTAGCGTGGGCATGCGAAGGAAAAAGCTCTTGGGATATTTCAAAAATATTAGGTTGCAGTGAGCGTACTGTCACTTTCCATTTAACCAATGCGCAAATGAAACTCAATACAACAAACCGCTGCCAAAGTATTTCTAAAGCAATTTTAACAGGAGCAATTGATTGCCCATACTTTAAAAATTGA |

**Supplementary Table S3.** Fluorescent Report Sequences – Original and Codon Optimized for *D. radiodurans*

| Name | Nucleotide Sequence | Amino Acids |
| --- | --- | --- |
| sfGFP | atggtgagcaagggcgaggagctgttcaccggggtggtgcccatcctggtcgagctggacggcgacgtaaacggccacaagttcagcgtgcgcggcgagggcgagggcgatgccaccaacggcaagctgaccctgaagttcatctgcaccaccggcaagctgcccgtgccctggcccaccctcgtgaccaccctgacctacggcgtgcagtgcttcagccgctaccccgaccacatgaagcgccacgacttcttcaagtccgccatgcccgaaggctacgtccaggagcgcaccatcagcttcaaggacgacggcacctacaagacccgcgccgaggtgaagttcgagggcgacaccctggtgaaccgcatcgagctgaagggcatcgacttcaaggaggacggcaacatcctggggcacaagctggagtacaacttcaacagccacaacgtctatatcaccgccgacaagcagaagaacggcatcaaggccaacttcaagatccgccacaacgtggaggacggcagcgtgcagctcgccgaccactaccagcagaacacccccatcggcgacggccccgtgctgctgcccgacaaccactacctgagcacccagtccgtgctgagcaaagaccccaacgagaagcgcgatcacatggtcctgctggagttcgtgaccgccgccgggatcactcacggcatggacgagctgtacaagtaa | MVSKGEELFTGVVPILVELDGDVNGHKFSVRGEGEGDATNGKLTLKFICTTGKLPVPWPTLVTTLTYGVQCFSRYPDHMKRHDFFKSAMPEGYVQERTISFKDDGTYKTRAEVKFEGDTLVNRIELKGIDFKEDGNILGHKLEYNFNSHNVYITADKQKNGIKANFKIRHNVEDGSVQLADHYQQNTPIGDGPVLLPDNHYLSTQSVLSKDPNEKRDHMVLLEFVTAAGITHGMDELYK* |
| sfGFP; D. radiodurans Codon-Optimized | ATGGTGAGCAAGGGCGAGGAGCTGTTCACCGGCGTGGTGCCCATCCTGGTGGAGCTGGACGGCGACGTGAACGGCCACAAGTTCAGCGTGCGCGGCGAGGGCGAGGGCGACGCCACCAACGGCAAGCTGACCCTGAAGTTCATCTGCACCACCGGCAAGCTGCCCGTGCCCTGGCCCACCCTGGTGACCACCCTGACCTACGGCGTGCAGTGCTTCAGCCGCTACCCCGACCACATGAAGCGCCACGACTTCTTCAAGAGCGCCATGCCCGAGGGCTACGTGCAGGAGCGCACCATCAGCTTCAAGGACGACGGCACCTACAAGACCCGCGCCGAGGTGAAGTTCGAGGGCGACACCCTGGTGAACCGCATCGAGCTGAAGGGCATCGACTTCAAGGAGGACGGCAACATCCTGGGCCACAAGCTGGAGTACAACTTCAACAGCCACAACGTGTACATCACCGCCGACAAGCAGAAGAACGGCATCAAGGCCAACTTCAAGATCCGCCACAACGTGGAGGACGGCAGCGTGCAGCTGGCCGACCACTACCAGCAGAACACCCCCATCGGCGACGGCCCCGTGCTGCTGCCCGACAACCACTACCTGAGCACCCAGAGCGTGCTGAGCAAGGACCCCAACGAGAAGCGCGACCACATGGTGCTGCTGGAGTTCGTGACCGCCGCCGGCATCACCCACGGCATGGACGAGCTGTACAAGTGA | MVSKGEELFTGVVPILVELDGDVNGHKFSVRGEGEGDATNGKLTLKFICTTGKLPVPWPTLVTTLTYGVQCFSRYPDHMKRHDFFKSAMPEGYVQERTISFKDDGTYKTRAEVKFEGDTLVNRIELKGIDFKEDGNILGHKLEYNFNSHNVYITADKQKNGIKANFKIRHNVEDGSVQLADHYQQNTPIGDGPVLLPDNHYLSTQSVLSKDPNEKRDHMVLLEFVTAAGITHGMDELYK* |
| mCherry | ATGGTTTCAAAAGGCGAAGAAGACAACATGGCGATTATCAAGGAATTTATGCGTTTCAAGGTCCACATGGAAGGCAGCGTCAATGGTCACGAATTTGAAATTGAAGGCGAAGGTGAAGGCCGTCCGTATGAAGGCACCCAGACGGCAAAACTGAAGGTCACCAAAGGCGGTCCGCTGCCGTTTGCTTGGGATATTCTGTCACCGCAATTCATGTATGGTTCGAAAGCGTACGTTAAGCATCCGGCCGATATCCCGGACTATCTGAAACTGTCCTTTCCGGAAGGCTTCAAATGGGAACGTGTTATGAACTTCGAAGATGGCGGTGTGGTTACCGTCACGCAGGATAGCTCTCTGCAAGACGGTGAATTTATTTATAAAGTGAAGCTGCGCGGCACCAATTTCCCGAGCGATGGTCCGGTTATGCAGAAAAAGACGATGGGCTGGGAAGCGAGTTCCGAACGTATGTACCCGGAAGACGGTGCCCTGAAAGGCGAAATCAAGCAGCGCCTGAAACTGAAGGATGGCGGTCACTATGACGCAGAAGTGAAAACCACGTACAAGGCTAAAAAGCCGGTCCAACTGCCGGGTGCATACAACGTGAACATCAAGCTGGATATCACCAGCCATAACGAAGACTATACGATCGTTGAACAGTACGAACGTGCAGAAGGCCGCCACTCTACCGGCGGTATGGATGAACTGTACAAATAA | MVSKGEEDNMAIIKEFMRFKVHMEGSVNGHEFEIEGEGEGRPYEGTQTAKLKVTKGGPLPFAWDILSPQFMYGSKAYVKHPADIPDYLKLSFPEGFKWERVMNFEDGGVVTVTQDSSLQDGEFIYKVKLRGTNFPSDGPVMQKKTMGWEASSERMYPEDGALKGEIKQRLKLKDGGHYDAEVKTTYKAKKPVQLPGAYNVNIKLDITSHNEDYTIVEQYERAEGRHSTGGMDELYK* |
| mCherry; D. radiodurans Codon-Optimized | ATGGTGAGCAAGGGCGAGGAGGACAACATGGCCATCATCAAGGAGTTCATGCGCTTCAAGGTGCACATGGAGGGCAGCGTGAACGGCCACGAGTTCGAGATCGAGGGCGAGGGCGAGGGCCGCCCCTACGAGGGCACCCAGACCGCCAAGCTGAAGGTGACCAAGGGCGGCCCCCTGCCCTTCGCCTGGGACATCCTGAGCCCCCAGTTCATGTACGGCAGCAAGGCCTACGTGAAGCACCCCGCCGACATCCCCGACTACCTGAAGCTGAGCTTCCCCGAGGGCTTCAAGTGGGAGCGCGTGATGAACTTCGAGGACGGCGGCGTGGTGACCGTGACCCAGGACAGCAGCCTGCAGGACGGCGAGTTCATCTACAAGGTGAAGCTGCGCGGCACCAACTTCCCCAGCGACGGCCCCGTGATGCAGAAGAAGACCATGGGCTGGGAGGCCAGCAGCGAGCGCATGTACCCCGAGGACGGCGCCCTGAAGGGCGAGATCAAGCAGCGCCTGAAGCTGAAGGACGGCGGCCACTACGACGCCGAGGTGAAGACCACCTACAAGGCCAAGAAGCCCGTGCAGCTGCCCGGCGCCTACAACGTGAACATCAAGCTGGACATCACCAGCCACAACGAGGACTACACCATCGTGGAGCAGTACGAGCGCGCCGAGGGCCGCCACAGCACCGGCGGCATGGACGAGCTGTACAAGTGA | MVSKGEEDNMAIIKEFMRFKVHMEGSVNGHEFEIEGEGEGRPYEGTQTAKLKVTKGGPLPFAWDILSPQFMYGSKAYVKHPADIPDYLKLSFPEGFKWERVMNFEDGGVVTVTQDSSLQDGEFIYKVKLRGTNFPSDGPVMQKKTMGWEASSERMYPEDGALKGEIKQRLKLKDGGHYDAEVKTTYKAKKPVQLPGAYNVNIKLDITSHNEDYTIVEQYERAEGRHSTGGMDELYK* |
| mTurq | ATGGTGAGCAAGGGCGAGGAGCTGTTCACCGGGGTGGTGCCCATCCTGGTCGAGCTGGACGGCGACGTAAACGGCCACAAGTTCAGCGTGTCCGGCGAGGGCGAGGGCGATGCCACCTACGGCAAGCTGACCCTGAAGTTCATCTGCACCACCGGCAAGCTGCCCGTGCCCTGGCCCACCCTCGTGACCACCCTGTCCTGGGGCGTGCAGTGCTTCGCCCGCTACCCCGACCACATGAAGCAGCACGACTTCTTCAAGTCCGCCATGCCCGAAGGCTACGTCCAGGAGCGCACCATCTTCTTCAAGGACGACGGCAACTACAAGACCCGCGCCGAGGTGAAGTTCGAGGGCGACACCCTGGTGAACCGCATCGAGCTGAAGGGCATCGACTTCAAGGAGGACGGCAACATCCTGGGGCACAAGCTGGAGTACAACTACATCAGCGACAACGTCTATATCACCGCCGACAAGCAGAAGAACGGCATCAAGGCCAACTTCAAGATCCGCCACAACATCGAGGACGGCGGCGTGCAGCTCGCCGACCACTACCAGCAGAACACCCCCATCGGCGACGGCCCCGTGCTGCTGCCCGACAACCACTACCTGAGCACCCAGTCCAAGCTGAGCAAAGACCCCAACGAGAAGCGCGATCACATGGTCCTGCTGGAGTTCGTGACCGCCGCCGGGATCACTCTCGGCATGGACGAGCTGTACAAGTAA | MVSKGEELFTGVVPILVELDGDVNGHKFSVSGEGEGDATYGKLTLKFICTTGKLPVPWPTLVTTLSWGVQCFARYPDHMKQHDFFKSAMPEGYVQERTIFFKDDGNYKTRAEVKFEGDTLVNRIELKGIDFKEDGNILGHKLEYNYISDNVYITADKQKNGIKANFKIRHNIEDGGVQLADHYQQNTPIGDGPVLLPDNHYLSTQSKLSKDPNEKRDHMVLLEFVTAAGITLGMDELYK* |
| mTurq; D. radiodurans Codon-Optimized | ATGGTGAGCAAGGGCGAGGAGCTGTTCACCGGCGTGGTGCCCATCCTGGTGGAGCTGGACGGCGACGTGAACGGCCACAAGTTCAGCGTGAGCGGCGAGGGCGAGGGCGACGCCACCTACGGCAAGCTGACCCTGAAGTTCATCTGCACCACCGGCAAGCTGCCCGTGCCCTGGCCCACCCTGGTGACCACCCTGAGCTGGGGCGTGCAGTGCTTCGCCCGCTACCCCGACCACATGAAGCAGCACGACTTCTTCAAGAGCGCCATGCCCGAGGGCTACGTGCAGGAGCGCACCATCTTCTTCAAGGACGACGGCAACTACAAGACCCGCGCCGAGGTGAAGTTCGAGGGCGACACCCTGGTGAACCGCATCGAGCTGAAGGGCATCGACTTCAAGGAGGACGGCAACATCCTGGGCCACAAGCTGGAGTACAACTACATCAGCGACAACGTGTACATCACCGCCGACAAGCAGAAGAACGGCATCAAGGCCAACTTCAAGATCCGCCACAACATCGAGGACGGCGGCGTGCAGCTGGCCGACCACTACCAGCAGAACACCCCCATCGGCGACGGCCCCGTGCTGCTGCCCGACAACCACTACCTGAGCACCCAGAGCAAGCTGAGCAAGGACCCCAACGAGAAGCGCGACCACATGGTGCTGCTGGAGTTCGTGACCGCCGCCGGCATCACCCTGGGCATGGACGAGCTGTACAAGTGA | MVSKGEELFTGVVPILVELDGDVNGHKFSVSGEGEGDATYGKLTLKFICTTGKLPVPWPTLVTTLSWGVQCFARYPDHMKQHDFFKSAMPEGYVQERTIFFKDDGNYKTRAEVKFEGDTLVNRIELKGIDFKEDGNILGHKLEYNYISDNVYITADKQKNGIKANFKIRHNIEDGGVQLADHYQQNTPIGDGPVLLPDNHYLSTQSKLSKDPNEKRDHMVLLEFVTAAGITLGMDELYK* |
| mOrange | ATGGTGAGCAAGGGCGAGGAGAATAACATGGCCATCATCAAGGAGTTCATGCGCTTCAAGGTGCGCATGGAGGGCTCCGTGAACGGCCACGAGTTCGAGATCGAGGGCGAGGGCGAGGGCCGCCCCTACGAGGGCTTTCAGACCGCTAAGCTGAAGGTGACCAAGGGTGGCCCCCTGCCCTTCGCCTGGGACATCCTGTCCCCTCAGTTCACCTACGGCTCCAAGGCCTACGTGAAGCACCCCGCCGACATCCCCGACTACTTCAAGCTGTCCTTCCCCGAGGGCTTCAAGTGGGAGCGCGTGATGAACTTCGAGGACGGCGGCGTGGTGACCGTGACCCAGGACTCCTCCCTGCAGGACGGCGAGTTCATCTACAAGGTGAAGCTGCGCGGCACCAACTTCCCCTCCGACGGCCCCGTAATGCAGAAGAAGACCATGGGCTGGGAGGCCTCCTCCGAGCGGATGTACCCCGAGGACGGCGCCCTGAAGGGCGAGATCAAGATGAGGCTGAAGCTGAAGGACGGCGGCCACTACACCTCCGAGGTCAAGACCACCTACAAGGCCAAGAAGCCCGTGCAGCTGCCCGGCGCCTACATCGTCGGCATCAAGTTGGACATCACCTCCCACAACGAGGACTACACCATCGTGGAACAGTACGAACGCGCCGAGGGCCGCCACTCCACCGGCGGCATGGACGAGCTGTACAAGTAG | MVSKGEENNMAIIKEFMRFKVRMEGSVNGHEFEIEGEGEGRPYEGFQTAKLKVTKGGPLPFAWDILSPQFTYGSKAYVKHPADIPDYFKLSFPEGFKWERVMNFEDGGVVTVTQDSSLQDGEFIYKVKLRGTNFPSDGPVMQKKTMGWEASSERMYPEDGALKGEIKMRLKLKDGGHYTSEVKTTYKAKKPVQLPGAYIVGIKLDITSHNEDYTIVEQYERAEGRHSTGGMDELYK* |
| mOrange; D. radiodurans Codon-Optimized | ATGGTGAGCAAGGGCGAGGAGAACAACATGGCCATCATCAAGGAGTTCATGCGCTTCAAGGTGCGCATGGAGGGCAGCGTGAACGGCCACGAGTTCGAGATCGAGGGCGAGGGCGAGGGCCGCCCCTACGAGGGCTTCCAGACCGCCAAGCTGAAGGTGACCAAGGGCGGCCCCCTGCCCTTCGCCTGGGACATCCTGAGCCCCCAGTTCACCTACGGCAGCAAGGCCTACGTGAAGCACCCCGCCGACATCCCCGACTACTTCAAGCTGAGCTTCCCCGAGGGCTTCAAGTGGGAGCGCGTGATGAACTTCGAGGACGGCGGCGTGGTGACCGTGACCCAGGACAGCAGCCTGCAGGACGGCGAGTTCATCTACAAGGTGAAGCTGCGCGGCACCAACTTCCCCAGCGACGGCCCCGTGATGCAGAAGAAGACCATGGGCTGGGAGGCCAGCAGCGAGCGCATGTACCCCGAGGACGGCGCCCTGAAGGGCGAGATCAAGATGCGCCTGAAGCTGAAGGACGGCGGCCACTACACCAGCGAGGTGAAGACCACCTACAAGGCCAAGAAGCCCGTGCAGCTGCCCGGCGCCTACATCGTGGGCATCAAGCTGGACATCACCAGCCACAACGAGGACTACACCATCGTGGAGCAGTACGAGCGCGCCGAGGGCCGCCACAGCACCGGCGGCATGGACGAGCTGTACAAGTGA | MVSKGEENNMAIIKEFMRFKVRMEGSVNGHEFEIEGEGEGRPYEGFQTAKLKVTKGGPLPFAWDILSPQFTYGSKAYVKHPADIPDYFKLSFPEGFKWERVMNFEDGGVVTVTQDSSLQDGEFIYKVKLRGTNFPSDGPVMQKKTMGWEASSERMYPEDGALKGEIKMRLKLKDGGHYTSEVKTTYKAKKPVQLPGAYIVGIKLDITSHNEDYTIVEQYERAEGRHSTGGMDELYK* |
| sfGFP | atggtgagcaagggcgaggagctgttcaccggggtggtgcccatcctggtcgagctggacggcgacgtaaacggccacaagttcagcgtgcgcggcgagggcgagggcgatgccaccaacggcaagctgaccctgaagttcatctgcaccaccggcaagctgcccgtgccctggcccaccctcgtgaccaccctgacctacggcgtgcagtgcttcagccgctaccccgaccacatgaagcgccacgacttcttcaagtccgccatgcccgaaggctacgtccaggagcgcaccatcagcttcaaggacgacggcacctacaagacccgcgccgaggtgaagttcgagggcgacaccctggtgaaccgcatcgagctgaagggcatcgacttcaaggaggacggcaacatcctggggcacaagctggagtacaacttcaacagccacaacgtctatatcaccgccgacaagcagaagaacggcatcaaggccaacttcaagatccgccacaacgtggaggacggcagcgtgcagctcgccgaccactaccagcagaacacccccatcggcgacggccccgtgctgctgcccgacaaccactacctgagcacccagtccgtgctgagcaaagaccccaacgagaagcgcgatcacatggtcctgctggagttcgtgaccgccgccgggatcactcacggcatggacgagctgtacaagtaa | MVSKGEELFTGVVPILVELDGDVNGHKFSVRGEGEGDATNGKLTLKFICTTGKLPVPWPTLVTTLTYGVQCFSRYPDHMKRHDFFKSAMPEGYVQERTISFKDDGTYKTRAEVKFEGDTLVNRIELKGIDFKEDGNILGHKLEYNFNSHNVYITADKQKNGIKANFKIRHNVEDGSVQLADHYQQNTPIGDGPVLLPDNHYLSTQSVLSKDPNEKRDHMVLLEFVTAAGITHGMDELYK* |

**Supplementary Table S4.** RBS Library Sequences

| Name | Variable RBS Sequence | Predicted Strength | *In vivo* Signal |
| --- | --- | --- | --- |
| RBS001 | GGAGG | 50.6212217 | 242.09 |
| RBS002 | ATCGA | 1.41667595 | 63.5 |
| RBS003 | AAGTT | 1.59971518 | 174.73 |
| RBS004 | GGTCT | 2.9635481 | 38.2 |
| RBS005 | TTCGT | 1.59971518 | 30.78 |
| RBS006 | CAATC | 1.12613458 | 29.72 |
| RBS007 | GGGTG | 0.91967995 | 50.41 |
| RBS008 | CGGAC | 0.40909404 | 22.9 |
| RBS009 | AGCCT | 1.12613458 | 11.72 |
| RBS010 | GGGCT | 2.29452456 | 85.71 |
| RBS011 | ACCTA | 0.27040035 | 54.57 |
| RBS012 | ATAGG | 2.29452456 | 57.66 |
| RBS013 | ATCGA | 0.27040035 | 105.3 |
| RBS014 | AGCTG | 0.66215064 | 34.86 |
| RBS015 | GTTTC | 1.59971518 | 38.6 |
| RBS016 | ATGGT | 1.59971518 | 200.89 |
| RBS017 | CTCTG | 1.59971518 | 31.11 |
| RBS018 | TTTGG | 2.93699242 | 27.4 |
| RBS019 | GGAAG | 4.50383461 | 240.79 |
| RBS020 | CAGTT | 4.28630024 | 84.98 |
| RBS021 | AGGCC | 3.11395121 | 82.06 |
| RBS022 | AGGGG | 21.2760983 | 63.55 |
| RBS023 | CGCTGC | 4.00649751 | 191.6 |
| RBS024 | GCCGG | 2.35576185 | 1267.19 |
| RBS025 | TGGTA | 1.92388078 | 96.34 |
| RBS026 | AGGTT | 4.50383461 | 39.42 |
| RBS027 | AGGTA | 4.50383461 | 79 |
| RBS028 | AGGTC | 2.99034356 | 34.87 |
| RBS029 | CATGC | 4.1347234 | 22.29 |
| RBS030 | AGTAT | 2.72067182 | 49.43 |
| RBS031 | AGTAA | 2.72067182 | 129.53 |
| RBS032 | CTCGAT | 2.72067182 | 39.49 |
| RBS033 | AGTCG | 1.07174249 | 32.52 |
| RBS034 | GGATG | 1.49528829 | 1266.42 |
| RBS035 | AATTC | 0.70521245 | 43.77 |
| RBS036 | TGGAA | 1.49528829 | 76.3 |
| RBS037 | GGGAG | 4.5856462 | 884.85 |
| RBS038 | TCGCG | 1.26592362 | 53.8 |
| RBS039 | ATGGA | 5.98419089 | 355.27 |
| RBS040 | GGGCT | 0.74434519 | 239.8 |
| RBS041 | TGGGC | 2.06752268 | 106.09 |
| RBS042 | CTTCC | 2.72067182 | 15.62 |
| RBS043 | GTAGG | 2.72067182 | 531.8 |
| RBS044 | CGTGT | 2.68418511 | 62.85 |
| RBS045 | ATAAG | 1.46861131 | 20.61 |
| RBS046 | ATTTT | 1.06213905 | 39.05 |
| RBS047 | GTGGT | 1.06213905 | 71.46 |
| RBS048 | GTGTA | 1.06213905 | 56.29 |
| RBS049 | ATACG | 1.20988819 | 36.18 |
| RBS050 | AAACT | 1.06213905 | 38.72 |
| RBS051 | TTAAT | 0.70521255 | 244.26 |
| RBS052 | AGCAA | 1.06213905 | 76.08 |
| RBS053 | GAATG | 9.5470717 | 315.8 |
| RBS054 | ATAGT | 1.03822539 | 64.09 |
| RBS055 | GAGTC | 1.41455754 | 72.2 |
| RBS056 | CAGTC | 0.68623953 | 55.5 |
| RBS057 | GGGAC | 0.5888733 | 147.45 |
| RBS058 | ATATT | 1.06213905 | 71.41 |
| RBS059 | CTAAC | 1.06213905 | 31.82 |
| RBS060 | AGGGC | 0.89517797 | 57.33 |
| RBS061 | ACTGC | 1.37893469 | 37.16 |
| RBS062 | ATCAT | 1.0017801 | 57.04 |
| RBS063 | CGTTT | 0.73436293 | 64.11 |
| RBS064 | TAACC | 1.0017801 | 39.98 |
| RBS065 | ATCCG | 0.28023736 | 187.5 |
| RBS066 | TGCG | 0.60788421 | 62.54 |
| RBS067 | ATCCA | 0.57076499 | 33.3 |
| RBS068 | ATCCC | 0.60788421 | 68.74 |
| RBS069 | ATCGG | 4.09656932 | 384.38 |
| RBS070 | ATCGT | 0.96201547 | 38.96 |
| RBS071 | ATCGA | 1.86142908 | 96.14 |
| RBS072 | ATCGC | 0.85194177 | 44.93 |
| RBS073 | ATCTG | 0.68315779 | 1366.77 |
| RBS074 | GGCCC | 1.06213905 | 67.9 |
| RBS075 | CAGTC | 1.06213905 | 122.54 |
| RBS076 | CCTTG | 0.82552138 | 90.68 |
| RBS077 | ATGAG | 3.03099098 | 38.96 |
| RBS078 | CTATG | 1.50880736 | 117.46 |
| RBS079 | CCACG | 1.93255851 | 17.77 |
| RBS080 | CACCC | 0.91143918 | 53.75 |
| RBS081 | ATGCG | 0.53818464 | 52.31 |
| RBS082 | ATGCT | 0.72451445 | 23.27 |
| RBS083 | ATGCA | 0.48978329 | 38.24 |
| RBS084 | ATGCC | 0.45987566 | 16.99 |
| RBS085 | CCACG | 7.90278409 | 33.66 |
| RBS086 | ATGGT | 1.42401071 | 43.35 |
| RBS087 | ATGGC | 1.5372345 | 49.86 |
| RBS088 | ATGTG | 0.50532191 | 414.09 |
| RBS089 | ATGTT | 0.35903926 | 264.56 |
| RBS090 | ATGTA | 0.44361301 | 43.82 |
| RBS091 | ATGTC | 0.16334311 | 33.42 |
| RBS092 | ATTAG | 1.46861131 | 25.03 |
| RBS093 | CTCCG | 1.06213905 | 11.25 |
| RBS095 | GGGCG | 1.06213905 | 86.81 |
| RBS096 | ATTCG | 0.86328896 | 71.52 |
| RBS097 | GGGAG | 1.06213905 | 256.42 |
| RBS098 | AGCGT | 0.45167134 | 193.93 |
| RBS099 | ATTCC | 0.54564781 | 1004.03 |
| RBS100 | ATTGG | 3.77779926 | 599.94 |
| RBS101 | ATTTT | 0.59973193 | 68.04 |
| RBS102 | TCTCCC | 1.17621118 | 10829.78 |
| RBS103 | TCGGA | 0.96201547 | 88.84 |
| RBS104 | TATAA | 0.81790296 | 69.82 |
| RBS105 | TATGT | 1.06213905 | 43.71 |
| RBS106 | GTGAAA | 1.06213905 | 66.42 |
| RBS107 | ATTTC | 1.04318928 | 57.55 |
| RBS108 | CAAAG | 1.94005236 | 37.07 |
| RBS109 | GCAAG | 0.92799498 | 43.27 |
| RBS110 | CAAAA | 1.00629904 | 77.57 |
| RBS111 | ATTCA | 0.45167134 | 37.09 |
| RBS112 | CCCCC | 1.00629904 | 27.70 |
| RBS114 | TTCCC | 1.00629904 | 40.11 |
| RBS115 | ACGTC | 0.18197397 | 179.69 |
| RBS116 | CAGGC | 3.92913401 | 54.61 |
| RBS119 | AAGGG | 11.1119732 | 56.26 |
| RBS121 | ACCAA | 0.63873445 | 47.93 |
| RBS122 | CTCCC | 1.00629904 | 66.38 |
| RBS123 | CTTCA | 0.33110108 | 66.29 |
| RBS126 | TCCGA | 0.72343096 | 31.42 |
| RBS128 | CGGTG | 1.08143253 | 85.71 |
| RBS129 | AGAAG | 4.07927645 | 52.52 |
| RBS131 | CTCAG | 1.00629904 | 61.40 |
| RBS133 | TGGAC | 4.97258025 | 185.79 |
| RBS134 | TGAAT | 1.65537764 | 33.45 |
| RBS137 | AGTAC | 2.72067182 | 35.68 |

**Supplementary Table S5.** XenO Fusion Proteins used in this study

| Name | Sequence (5'-3') | Amino Acid Sequence |
| --- | --- | --- |
| Xenotext Candidate 1; C-Terminal | ATGCATCACCATCACCATCACGTGACCGCCTTCCAGGCCTACCGCGAGAAGTACCTGAAGCGCCTGTTCTGGATGCTGCCCATCGAGTGCTTCCAGGTGGCCCCCGCCTACGCCATGCGCATCGGCCTGCAGTGCGGCCTGTACGCCTGGTACGAGGCCTTCCCCGAGGTGACCCAGTGCCGCATGCTGAAGAAGATCCAGTGCCGCATGCTGAAGAAGGAGAAGGTGACCGCCCAGAACACCCGCGTGTACGAGAACGACATCCCCGAGGTGACCACCGAGGGCCCCGCCAAGGTGCAGCGCATCCTGACCAAGGTGCAGCGCGGCCGCATGCGCTACGCCATCCCCGCCCCCGCCCAGTGCACCGAGGGCCTGTTCCAGTGCCGCGGCGAGATGGACATCCAGTGCCGCGGCGAGATGGACGAGAAGTACCTGAAGCGCGGTGGTTCTGGTATGGTCTCGAAAGGGGAGGAAGACAACATGGCGATCATCAAAGAGTTTATGCGCTTCAAAGTCCACATGGAGGGCAGCGTGAACGGTCACGAATTCGAAATTGAAGGGGAAGGCGAGGGCCGGCCCTACGAGGGCACCCAAACGGCCAAGCTGAAGGTCACCAAGGGTGGTCCTCTCCCCTTCGCGTGGGACATCCTCAGCCCGCAGTTCATGTATGGCTCCAAGGCGTACGTCAAGCACCCGGCCGACATCCCGGACTACCTGAAGCTCTCGTTCCCCGAAGGCTTCAAGTGGGAACGCGTGATGAACTTCGAAGACGGCGGCGTCGTGACGGTGACCCAGGATTCCAGCCTCCAGGACGGCGAGTTCATTTACAAGGTGAAGCTCCGTGGCACCAACTTCCCCTCGGACGGCCCCGTGATGCAGAAGAAGACCATGGGCTGGGAAGCCAGCAGCGAACGCATGTACCCCGAAGACGGCGCTCTGAAGGGCGAAATCAAGCAGCGCCTGAAGCTGAAGGACGGCGGCCACTACGACGCCGAGGTGAAGACCACCTACAAGGCCAAGAAGCCCGTGCAGCTGCCCGGCGCCTACAACGTGAACATCAAGCTGGACATCACCAGCCACAACGAAGACTACACCATCGTGGAACAGTACGAGCGCGCCGAGGGCCGCCACAGCACCGGCGGCATGGACGAGCTGTACAAGTGA | MHHHHHHVTAFQAYREKYLKRLFWMLPIECFQVAPAYAMRIGLQCGLYAWYEAFPEVTQC RMLKKIQCRMLKKEKVTAQNTRVYENDIPEVTTEGPAKVQRILTKVQRGRMRYAIPAPAQ CTEGLFQCRGEMDIQCRGEMDEKYLKRGGSGMVSKGEEDNMAIIKEFMRFKVHMEGSVNG HEFEIEGEGEGRPYEGTQTAKLKVTKGGPLPFAWDILSPQFMYGSKAYVKHPADIPDYLK LSFPEGFKWERVMNFEDGGVVTVTQDSSLQDGEFIYKVKLRGTNFPSDGPVMQKKTMGWE ASSERMYPEDGALKGEIKQRLKLKDGGHYDAEVKTTYKAKKPVQLPGAYNVNIKLDITSH NEDYTIVEQYERAEGRHSTGGMDELYK* |
| Xenotext Candidate 1; N-Terminal | ATGGTCTCGAAAGGGGAGGAAGACAACATGGCGATCATCAAAGAGTTTATGCGCTTCAAAGTCCACATGGAGGGCAGCGTGAACGGTCACGAATTCGAAATTGAAGGGGAAGGCGAGGGCCGGCCCTACGAGGGCACCCAAACGGCCAAGCTGAAGGTCACCAAGGGTGGTCCTCTCCCCTTCGCGTGGGACATCCTCAGCCCGCAGTTCATGTATGGCTCCAAGGCGTACGTCAAGCACCCGGCCGACATCCCGGACTACCTGAAGCTCTCGTTCCCCGAAGGCTTCAAGTGGGAACGCGTGATGAACTTCGAAGACGGCGGCGTCGTGACGGTGACCCAGGATTCCAGCCTCCAGGACGGCGAGTTCATTTACAAGGTGAAGCTCCGTGGCACCAACTTCCCCTCGGACGGCCCCGTGATGCAGAAGAAGACCATGGGCTGGGAAGCCAGCAGCGAACGCATGTACCCCGAAGACGGCGCTCTGAAGGGCGAAATCAAGCAGCGCCTGAAGCTGAAGGACGGCGGCCACTACGACGCCGAGGTGAAGACCACCTACAAGGCCAAGAAGCCCGTGCAGCTGCCCGGCGCCTACAACGTGAACATCAAGCTGGACATCACCAGCCACAACGAAGACTACACCATCGTGGAACAGTACGAGCGCGCCGAGGGCCGCCACAGCACCGGCGGCATGGACGAGCTGTACAAGGGTGGTTCTGGTGTGACCGCCTTCCAGGCCTACCGCGAGAAGTACCTGAAGCGCCTGTTCTGGATGCTGCCCATCGAGTGCTTCCAGGTGGCCCCCGCCTACGCCATGCGCATCGGCCTGCAGTGCGGCCTGTACGCCTGGTACGAGGCCTTCCCCGAGGTGACCCAGTGCCGCATGCTGAAGAAGATCCAGTGCCGCATGCTGAAGAAGGAGAAGGTGACCGCCCAGAACACCCGCGTGTACGAGAACGACATCCCCGAGGTGACCACCGAGGGCCCCGCCAAGGTGCAGCGCATCCTGACCAAGGTGCAGCGCGGCCGCATGCGCTACGCCATCCCCGCCCCCGCCCAGTGCACCGAGGGCCTGTTCCAGTGCCGCGGCGAGATGGACATCCAGTGCCGCGGCGAGATGGACGAGAAGTACCTGAAGCGCCATCACCATCACCATCACTGA | MVSKGEEDNMAIIKEFMRFKVHMEGSVNGHEFEIEGEGEGRPYEGTQTAKLKVTKGGPLP FAWDILSPQFMYGSKAYVKHPADIPDYLKLSFPEGFKWERVMNFEDGGVVTVTQDSSLQD GEFIYKVKLRGTNFPSDGPVMQKKTMGWEASSERMYPEDGALKGEIKQRLKLKDGGHYDA EVKTTYKAKKPVQLPGAYNVNIKLDITSHNEDYTIVEQYERAEGRHSTGGMDELYKGGSG VTAFQAYREKYLKRLFWMLPIECFQVAPAYAMRIGLQCGLYAWYEAFPEVTQCRMLKKIQ CRMLKKEKVTAQNTRVYENDIPEVTTEGPAKVQRILTKVQRGRMRYAIPAPAQCTEGLFQ CRGEMDIQCRGEMDEKYLKRHHHHHH* |
| Xenotext Candidate 2; C-Terminal | ATGCATCACCATCACCATCACGTAACTGCATTTCAAGCATACCGTGAAAAATACCTCAAACGTCTCTTTTGGATGCTCCCTATCGAATGCTTTCAAGTAGCACCTGCATACGCAATGCGTATCGGCCTCCAATGCGGCCTCTACGCATGGTACGAAGCATTTCCTGAAGTAACTCAATGCCGTATGCTCAAAAAAATCCAATGCCGTATGCTCAAAAAAGAAAAAGTAACTGCACAAAATACTCGTGTATACGAAAATGATATCCCTGAAGTAACTACTGAAGGCCCTGCAAAAGTACAACGTATCCTCACTAAAGTACAACGTGGCCGTATGCGTTACGCAATCCCTGCACCTGCACAATGCACTGAAGGCCTCTTTCAATGCCGTGGCGAAATGGATATCCAATGCCGTGGCGAAATGGATGAAAAATACCTCAAACGTGGTGGTTCTGGTATGGTCTCGAAAGGGGAGGAAGACAACATGGCGATCATCAAAGAGTTTATGCGCTTCAAAGTCCACATGGAGGGCAGCGTGAACGGTCACGAATTCGAAATTGAAGGGGAAGGCGAGGGCCGGCCCTACGAGGGCACCCAAACGGCCAAGCTGAAGGTCACCAAGGGTGGTCCTCTCCCCTTCGCGTGGGACATCCTCAGCCCGCAGTTCATGTATGGCTCCAAGGCGTACGTCAAGCACCCGGCCGACATCCCGGACTACCTGAAGCTCTCGTTCCCCGAAGGCTTCAAGTGGGAACGCGTGATGAACTTCGAAGACGGCGGCGTCGTGACGGTGACCCAGGATTCCAGCCTCCAGGACGGCGAGTTCATTTACAAGGTGAAGCTCCGTGGCACCAACTTCCCCTCGGACGGCCCCGTGATGCAGAAGAAGACCATGGGCTGGGAAGCCAGCAGCGAACGCATGTACCCCGAAGACGGCGCTCTGAAGGGCGAAATCAAGCAGCGCCTGAAGCTGAAGGACGGCGGCCACTACGACGCCGAGGTGAAGACCACCTACAAGGCCAAGAAGCCCGTGCAGCTGCCCGGCGCCTACAACGTGAACATCAAGCTGGACATCACCAGCCACAACGAAGACTACACCATCGTGGAACAGTACGAGCGCGCCGAGGGCCGCCACAGCACCGGCGGCATGGACGAGCTGTACAAGTGA | MHHHHHHVTAFQAYREKYLKRLFWMLPIECFQVAPAYAMRIGLQCGLYAWYEAFPEVTQC RMLKKIQCRMLKKEKVTAQNTRVYENDIPEVTTEGPAKVQRILTKVQRGRMRYAIPAPAQ CTEGLFQCRGEMDIQCRGEMDEKYLKRGGSGMVSKGEEDNMAIIKEFMRFKVHMEGSVNG HEFEIEGEGEGRPYEGTQTAKLKVTKGGPLPFAWDILSPQFMYGSKAYVKHPADIPDYLK LSFPEGFKWERVMNFEDGGVVTVTQDSSLQDGEFIYKVKLRGTNFPSDGPVMQKKTMGWE ASSERMYPEDGALKGEIKQRLKLKDGGHYDAEVKTTYKAKKPVQLPGAYNVNIKLDITSH NEDYTIVEQYERAEGRHSTGGMDELYK* |
| Xenotext Candidate 2; N-Terminal | ATGGTCTCGAAAGGGGAGGAAGACAACATGGCGATCATCAAAGAGTTTATGCGCTTCAAAGTCCACATGGAGGGCAGCGTGAACGGTCACGAATTCGAAATTGAAGGGGAAGGCGAGGGCCGGCCCTACGAGGGCACCCAAACGGCCAAGCTGAAGGTCACCAAGGGTGGTCCTCTCCCCTTCGCGTGGGACATCCTCAGCCCGCAGTTCATGTATGGCTCCAAGGCGTACGTCAAGCACCCGGCCGACATCCCGGACTACCTGAAGCTCTCGTTCCCCGAAGGCTTCAAGTGGGAACGCGTGATGAACTTCGAAGACGGCGGCGTCGTGACGGTGACCCAGGATTCCAGCCTCCAGGACGGCGAGTTCATTTACAAGGTGAAGCTCCGTGGCACCAACTTCCCCTCGGACGGCCCCGTGATGCAGAAGAAGACCATGGGCTGGGAAGCCAGCAGCGAACGCATGTACCCCGAAGACGGCGCTCTGAAGGGCGAAATCAAGCAGCGCCTGAAGCTGAAGGACGGCGGCCACTACGACGCCGAGGTGAAGACCACCTACAAGGCCAAGAAGCCCGTGCAGCTGCCCGGCGCCTACAACGTGAACATCAAGCTGGACATCACCAGCCACAACGAAGACTACACCATCGTGGAACAGTACGAGCGCGCCGAGGGCCGCCACAGCACCGGCGGCATGGACGAGCTGTACAAGGGTGGTTCTGGTGTAACTGCATTTCAAGCATACCGTGAAAAATACCTCAAACGTCTCTTTTGGATGCTCCCTATCGAATGCTTTCAAGTAGCACCTGCATACGCAATGCGTATCGGCCTCCAATGCGGCCTCTACGCATGGTACGAAGCATTTCCTGAAGTAACTCAATGCCGTATGCTCAAAAAAATCCAATGCCGTATGCTCAAAAAAGAAAAAGTAACTGCACAAAATACTCGTGTATACGAAAATGATATCCCTGAAGTAACTACTGAAGGCCCTGCAAAAGTACAACGTATCCTCACTAAAGTACAACGTGGCCGTATGCGTTACGCAATCCCTGCACCTGCACAATGCACTGAAGGCCTCTTTCAATGCCGTGGCGAAATGGATATCCAATGCCGTGGCGAAATGGATGAAAAATACCTCAAACGTCATCACCATCACCATCACTGA | MVSKGEEDNMAIIKEFMRFKVHMEGSVNGHEFEIEGEGEGRPYEGTQTAKLKVTKGGPLP FAWDILSPQFMYGSKAYVKHPADIPDYLKLSFPEGFKWERVMNFEDGGVVTVTQDSSLQD GEFIYKVKLRGTNFPSDGPVMQKKTMGWEASSERMYPEDGALKGEIKQRLKLKDGGHYDA EVKTTYKAKKPVQLPGAYNVNIKLDITSHNEDYTIVEQYERAEGRHSTGGMDELYKGGSG VTAFQAYREKYLKRLFWMLPIECFQVAPAYAMRIGLQCGLYAWYEAFPEVTQCRMLKKIQ CRMLKKEKVTAQNTRVYENDIPEVTTEGPAKVQRILTKVQRGRMRYAIPAPAQCTEGLFQ CRGEMDIQCRGEMDEKYLKRHHHHHH* |
| **Xenotext Candidate 3; C-Terminal (XenO)** | ATGCATCACCATCACCATCACGTGACCCGTTTCCAGCGTTACGCCGAGAAGTACCTGAAGGCCCTGTTCTGGATGCTGCCCATCGAGTGCTTCCAGGTGCGTCCCCGTTACCGTATGGCCATCGGTCTGCAGTGCGGTCTGTACCGTTGGTACGAGCGTTTCCCCGAGGTGACCCAGTGCGCCATGCTGAAGAAGATCCAGTGCGCCATGCTGAAGAAGGAGAAGGTGACCCGTCAGAACACCGCCGTGTACGAGAACGACATCCCCGAGGTGACCACCGAGGGTCCCCGTAAGGTGCAGGCCATCCTGACCAAGGTGCAGGCCGGTGCCATGGCCTACCGTATCCCCCGTCCCCGTCAGTGCACCGAGGGTCTGTTCCAGTGCGCCGGTGAGATGGACATCCAGTGCGCCGGTGAGATGGACGAGAAGTACCTGAAGGCCGGTGGTTCTGGTATGGTCTCGAAAGGGGAGGAAGACAACATGGCGATCATCAAAGAGTTTATGCGCTTCAAAGTCCACATGGAGGGCAGCGTGAACGGTCACGAATTCGAAATTGAAGGGGAAGGCGAGGGCCGGCCCTACGAGGGCACCCAAACGGCCAAGCTGAAGGTCACCAAGGGTGGTCCTCTCCCCTTCGCGTGGGACATCCTCAGCCCGCAGTTCATGTATGGCTCCAAGGCGTACGTCAAGCACCCGGCCGACATCCCGGACTACCTGAAGCTCTCGTTCCCCGAAGGCTTCAAGTGGGAACGCGTGATGAACTTCGAAGACGGCGGCGTCGTGACGGTGACCCAGGATTCCAGCCTCCAGGACGGCGAGTTCATTTACAAGGTGAAGCTCCGTGGCACCAACTTCCCCTCGGACGGCCCCGTGATGCAGAAGAAGACCATGGGCTGGGAAGCCAGCAGCGAACGCATGTACCCCGAAGACGGCGCTCTGAAGGGCGAAATCAAGCAGCGCCTGAAGCTGAAGGACGGCGGCCACTACGACGCCGAGGTGAAGACCACCTACAAGGCCAAGAAGCCCGTGCAGCTGCCCGGCGCCTACAACGTGAACATCAAGCTGGACATCACCAGCCACAACGAAGACTACACCATCGTGGAACAGTACGAGCGCGCCGAGGGCCGCCACAGCACCGGCGGCATGGACGAGCTGTACAAGTGA | MHHHHHHVTRFQRYAEKYLKALFWMLPIECFQVRPRYRMAIGLQCGLYRWYERFPEVTQC AMLKKIQCAMLKKEKVTRQNTAVYENDIPEVTTEGPRKVQAILTKVQAGAMAYRIPRPRQ CTEGLFQCAGEMDIQCAGEMDEKYLKAGGSGMVSKGEEDNMAIIKEFMRFKVHMEGSVNG HEFEIEGEGEGRPYEGTQTAKLKVTKGGPLPFAWDILSPQFMYGSKAYVKHPADIPDYLK LSFPEGFKWERVMNFEDGGVVTVTQDSSLQDGEFIYKVKLRGTNFPSDGPVMQKKTMGWE ASSERMYPEDGALKGEIKQRLKLKDGGHYDAEVKTTYKAKKPVQLPGAYNVNIKLDITSH NEDYTIVEQYERAEGRHSTGGMDELYK* |
| Xenotext Candidate 3; N-Terminal | ATGGTCTCGAAAGGGGAGGAAGACAACATGGCGATCATCAAAGAGTTTATGCGCTTCAAAGTCCACATGGAGGGCAGCGTGAACGGTCACGAATTCGAAATTGAAGGGGAAGGCGAGGGCCGGCCCTACGAGGGCACCCAAACGGCCAAGCTGAAGGTCACCAAGGGTGGTCCTCTCCCCTTCGCGTGGGACATCCTCAGCCCGCAGTTCATGTATGGCTCCAAGGCGTACGTCAAGCACCCGGCCGACATCCCGGACTACCTGAAGCTCTCGTTCCCCGAAGGCTTCAAGTGGGAACGCGTGATGAACTTCGAAGACGGCGGCGTCGTGACGGTGACCCAGGATTCCAGCCTCCAGGACGGCGAGTTCATTTACAAGGTGAAGCTCCGTGGCACCAACTTCCCCTCGGACGGCCCCGTGATGCAGAAGAAGACCATGGGCTGGGAAGCCAGCAGCGAACGCATGTACCCCGAAGACGGCGCTCTGAAGGGCGAAATCAAGCAGCGCCTGAAGCTGAAGGACGGCGGCCACTACGACGCCGAGGTGAAGACCACCTACAAGGCCAAGAAGCCCGTGCAGCTGCCCGGCGCCTACAACGTGAACATCAAGCTGGACATCACCAGCCACAACGAAGACTACACCATCGTGGAACAGTACGAGCGCGCCGAGGGCCGCCACAGCACCGGCGGCATGGACGAGCTGTACAAGGGTGGTTCTGGTGTGACCCGTTTCCAGCGTTACGCCGAGAAGTACCTGAAGGCCCTGTTCTGGATGCTGCCCATCGAGTGCTTCCAGGTGCGTCCCCGTTACCGTATGGCCATCGGTCTGCAGTGCGGTCTGTACCGTTGGTACGAGCGTTTCCCCGAGGTGACCCAGTGCGCCATGCTGAAGAAGATCCAGTGCGCCATGCTGAAGAAGGAGAAGGTGACCCGTCAGAACACCGCCGTGTACGAGAACGACATCCCCGAGGTGACCACCGAGGGTCCCCGTAAGGTGCAGGCCATCCTGACCAAGGTGCAGGCCGGTGCCATGGCCTACCGTATCCCCCGTCCCCGTCAGTGCACCGAGGGTCTGTTCCAGTGCGCCGGTGAGATGGACATCCAGTGCGCCGGTGAGATGGACGAGAAGTACCTGAAGGCCCATCACCATCACCATCACTGA | MVSKGEEDNMAIIKEFMRFKVHMEGSVNGHEFEIEGEGEGRPYEGTQTAKLKVTKGGPLP FAWDILSPQFMYGSKAYVKHPADIPDYLKLSFPEGFKWERVMNFEDGGVVTVTQDSSLQD GEFIYKVKLRGTNFPSDGPVMQKKTMGWEASSERMYPEDGALKGEIKQRLKLKDGGHYDA EVKTTYKAKKPVQLPGAYNVNIKLDITSHNEDYTIVEQYERAEGRHSTGGMDELYKGGSG VTRFQRYAEKYLKALFWMLPIECFQVRPRYRMAIGLQCGLYRWYERFPEVTQCAMLKKIQ CAMLKKEKVTRQNTAVYENDIPEVTTEGPRKVQAILTKVQAGAMAYRIPRPRQCTEGLFQ CAGEMDIQCAGEMDEKYLKAHHHHHH* |

**Supplementary Table S6.** Stains and Plasmids used in this study

| Strain or Plasmid | Description/Genotype | Reference |
| --- | --- | --- |
| *E. coli DH5α* | *fhuA2 lac(del)U169 phoA glnV44 Φ80' lacZ(del)M15 gyrA96 recA1 relA1 endA1 thi-1 hsdR17* | Contreras Lab, U. of Texas at Austin |
| *D. radiodurans R1* | Wild-type D. radiodurans R1 strain (ATCC 13939) | Contreras Lab, U. of Texas at Austin |
| CML 3791 | D. radiodurans R1 strain variant expressing a non-naitive, synthetic, genomically inserted mCherry-fused protein, Xenotext 1C | This Study |
| CML 3792 | D. radiodurans R1 strain variant expressing a non-naitive, synthetic, genomically inserted mCherry-fused protein, Xenotext 3C | This Study |
| CML 4209 | D. radiodurans R1 strain variant expressing a non-naitive, synthetic, genomically inserted mCherry-fused protein, Xenotext 3C, alongside a panel of ReRNA targeting sites upstream of the protein. | This Study |
| pUC19mPheS | Plamid for gene mutation in D. radiodurans containing mPhes cassette for counterselection by 4-cholorphenylalanine | Roland J. Saldanha and Thomas Lamkin |
| pMLKp1261_Xenotext_Candidate2_NTerm_mCherry | pMLKp1261 backbone from pRad-DR2009_3xFLAG, utilizing BamHI & SacII Rest. Digest to insert Candidate Gene from puc19 Genscript plasmid | This Study |
| pMLKp1261_Xenotext_Candidate2_CTerm_mCherry | pMLKp1261 backbone from pRad-DR2009_3xFLAG, utilizing BamHI & SacII Rest. Digest to insert Candidate Gene from puc19 Genscript plasmid | This Study |
| pMLKp1261_Xenotext_Candidate3_NTerm_mCherry | pMLKp1261 backbone from pRad-DR2009_3xFLAG, utilizing BamHI & SacII Rest. Digest to insert Candidate Gene from puc19 Genscript plasmid | This Study |
| pMLKp1261_Xenotext_Candidate3_CTerm_mCherry | pMLKp1261 backbone from pRad-DR2009_3xFLAG, utilizing BamHI & SacII Rest. Digest to insert Candidate Gene from puc19 Genscript plasmid | This Study |
| pMLKp1261_Xenotext_Candidate1_CTerm_mCherry | pMLKp1261 backbone from pRad-DR2009_3xFLAG, utilizing Gibson Asembly t to insert Candidate Gene from puc19 Genscript plasmid | This Study |
| pMLKp1261_Xenotext_Candidate2_CTerm_mCherry_Gibson | pMLKp1261 backbone from pRad-DR2009_3xFLAG, utilizing Gibson Assembly to insert Candidate Gene from puc19 Genscript plasmid | This Study |
| pAT00_20_001_sfGFP | pAT00 Base Plasmid construct containing: ColE1/pMB1, pGroES, Native GroES RBS Seq, sfGFP | This Study |
| pAT03_20_001_sfGFP | pAT00 Base Plasmid construct containing: pBBR1, pGroES, Native GroES Seq, sfGFP | This Study |
| pAT04_20_001_sfGFP | pAT00 Base Plasmid construct containing: RK2, pGroES, Native GroES Seq, sfGFP | This Study |
| pAT06_20_001_sfGFP | pAT00 Base Plasmid construct containing: pRO1600, pGroES, Native GroES Seq, sfGFP | This Study |
| pAT08_20_001_sfGFP | pAT00 Base Plasmid construct containing: pUB1110, pGroES, Native GroES Seq, sfGFP | This Study |
| pAT16_20_001_sfGFP | pAT00 Base Plasmid construct containing: cisII, pGroES, Native GroES Seq, sfGFP | This Study |
| pAT17_20_001_sfGFP | pAT00 Base Plasmid construct containing: cisMP, pGroES, Native GroES Seq, sfGFP | This Study |
| pAT00_00_001_sfGFP | pAT00 Base Plasmid construct containing: ColE1/pMB1, Anderson 100, Native GroES Seq, sfGFP | This Study |
| pAT00_01_001_sfGFP | pAT00 Base Plasmid construct containing: ColE1/pMB1, Anderson 101, Native GroES Seq, sfGFP | This Study |
| pAT00_02_001_sfGFP | pAT00 Base Plasmid construct containing: ColE1/pMB1, Anderson 102, Native GroES Seq, sfGFP | This Study |
| pAT00_03_001_sfGFP | pAT00 Base Plasmid construct containing: ColE1/pMB1, Anderson 103, Native GroES Seq, sfGFP | This Study |
| pAT00_04_001_sfGFP | pAT00 Base Plasmid construct containing: ColE1/pMB1, Anderson 104, Native GroES Seq, sfGFP | This Study |
| pAT00_05_001_sfGFP | pAT00 Base Plasmid construct containing: ColE1/pMB1, Anderson 105, Native GroES Seq, sfGFP | This Study |
| pAT00_06_001_sfGFP | pAT00 Base Plasmid construct containing: ColE1/pMB1, Anderson 106, Native GroES Seq, sfGFP | This Study |
| pAT00_07_001_sfGFP | pAT00 Base Plasmid construct containing: ColE1/pMB1, Anderson 107, Native GroES Seq, sfGFP | This Study |
| pAT00_08_001_sfGFP | pAT00 Base Plasmid construct containing: ColE1/pMB1, Anderson 108, Native GroES Seq, sfGFP | This Study |
| pAT00_09_001_sfGFP | pAT00 Base Plasmid construct containing: ColE1/pMB1, Anderson 109, Native GroES Seq, sfGFP | This Study |
| pAT00_10_001_sfGFP | pAT00 Base Plasmid construct containing: ColE1/pMB1, Anderson 110, Native GroES Seq, sfGFP | This Study |
| pAT00_11_001_sfGFP | pAT00 Base Plasmid construct containing: ColE1/pMB1, Anderson 111, Native GroES Seq, sfGFP | This Study |
| pAT00_12_001_sfGFP | pAT00 Base Plasmid construct containing: ColE1/pMB1, Anderson 112, Native GroES Seq, sfGFP | This Study |
| pAT00_13_001_sfGFP | pAT00 Base Plasmid construct containing: ColE1/pMB1, Anderson 113, Native GroES Seq, sfGFP | This Study |
| pAT00_14_001_sfGFP | pAT00 Base Plasmid construct containing: ColE1/pMB1, Anderson 114, Native GroES Seq, sfGFP | This Study |
| pAT00_15_001_sfGFP | pAT00 Base Plasmid construct containing: ColE1/pMB1, Anderson 115, Native GroES Seq, sfGFP | This Study |
| pAT00_16_001_sfGFP | pAT00 Base Plasmid construct containing: ColE1/pMB1, Anderson 116, Native GroES Seq, sfGFP | This Study |
| pAT00_17_001_sfGFP | pAT00 Base Plasmid construct containing: ColE1/pMB1, Anderson 117, Native GroES Seq, sfGFP | This Study |
| pAT00_18_001_sfGFP | pAT00 Base Plasmid construct containing: ColE1/pMB1, Anderson 118, Native GroES Seq, sfGFP | This Study |
| pAT00_19_001_sfGFP | pAT00 Base Plasmid construct containing: ColE1/pMB1, Anderson 119, Native GroES Seq, sfGFP | This Study |
| pAT00_20_001_sfGFP | pAT00 Base Plasmid construct containing: ColE1/pMB1, pGroES, Native GroES Seq, sfGFP | This Study |
| pAT00_21_001_sfGFP | pAT00 Base Plasmid construct containing: ColE1/pMB1, p1261, Native GroES Seq, sfGFP | This Study |
| pAT00_22_001_sfGFP | pAT00 Base Plasmid construct containing: ColE1/pMB1, p1348, Native GroES Seq, sfGFP | This Study |
| pAT00_23_001_sfGFP | pAT00 Base Plasmid construct containing: ColE1/pMB1, p1473, Native GroES Seq, sfGFP | This Study |
| pAT00_24_001_sfGFP | pAT00 Base Plasmid construct containing: ColE1/pMB1, p2508, Native GroES Seq, sfGFP | This Study |
| pAT00_25_001_sfGFP | pAT00 Base Plasmid construct containing: ColE1/pMB1, pAmyE, Native GroES Seq, sfGFP | This Study |
| pAT00_26_001_sfGFP | pAT00 Base Plasmid construct containing: ColE1/pMB1, pClpB, Native GroES Seq, sfGFP | This Study |
| pAT00_27_001_sfGFP | pAT00 Base Plasmid construct containing: ColE1/pMB1, pDnaK, Native GroES Seq, sfGFP | This Study |
| pAT00_28_001_sfGFP | pAT00 Base Plasmid construct containing: ColE1/pMB1, pGroEL, Native GroES Seq, sfGFP | This Study |
| pAT00_29_001_sfGFP | pAT00 Base Plasmid construct containing: ColE1/pMB1, pKatA, Native GroES Seq, sfGFP | This Study |
| pAT00_30_001_sfGFP | pAT00 Base Plasmid construct containing: ColE1/pMB1, pLexA, Native GroES Seq, sfGFP | This Study |
| pAT00_31_001_sfGFP | pAT00 Base Plasmid construct containing: ColE1/pMB1, pRecA, Native GroES Seq, sfGFP | This Study |
| pAT00_32_001_sfGFP | pAT00 Base Plasmid construct containing: ColE1/pMB1, pRecQ, Native GroES Seq, sfGFP | This Study |
| pAT00_33_001_sfGFP | pAT00 Base Plasmid construct containing: ColE1/pMB1, pRplL, Native GroES Seq, sfGFP | This Study |
| pAT00_34_001_sfGFP | pAT00 Base Plasmid construct containing: ColE1/pMB1, pRpmB, Native GroES Seq, sfGFP | This Study |
| pAT00_35_001_sfGFP | pAT00 Base Plasmid construct containing: ColE1/pMB1, pTufB, Native GroES Seq, sfGFP | This Study |
| pAT00_40_001_sfGFP | pAT00 Base Plasmid construct containing: ColE1/pMB1, LacI - IPTG, Native GroES Seq, sfGFP | This Study |
| pAT00_41_001_sfGFP | pAT00 Base Plasmid construct containing: ColE1/pMB1, LacIQ - DHBA, Native GroES Seq, sfGFP | This Study |
| pAT00_42_001_sfGFP | pAT00 Base Plasmid construct containing: ColE1/pMB1, LacI - OHC14, Native GroES Seq, sfGFP | This Study |
| pAT00_43_001_sfGFP | pAT00 Base Plasmid construct containing: ColE1/pMB1, LacIQ - Sal, Native GroES Seq, sfGFP | This Study |
| pAT00_44_001_sfGFP | pAT00 Base Plasmid construct containing: ColE1/pMB1, LacIQ - CA, Native GroES Seq, sfGFP | This Study |
| pAT00_45_001_sfGFP | pAT00 Base Plasmid construct containing: ColE1/pMB1, LacI - aTc, Native GroES Seq, sfGFP | This Study |
| pAT00_46_001_sfGFP | pAT00 Base Plasmid construct containing: ColE1/pMB1, LacIQ - Cho, Native GroES Seq, sfGFP | This Study |
| pAT00_47_001_sfGFP | pAT00 Base Plasmid construct containing: ColE1/pMB1, LacIQ - Ery, Native GroES Seq, sfGFP | This Study |
| pAT00_48_001_sfGFP | pAT00 Base Plasmid construct containing: ColE1/pMB1, LacIQ - Nar, Native GroES Seq, sfGFP | This Study |
| pAT00_49_001_sfGFP | pAT00 Base Plasmid construct containing: ColE1/pMB1, LacIQ - Va, Native GroES Seq, sfGFP | This Study |
| pAT00_50_001_sfGFP | pAT00 Base Plasmid construct containing: ColE1/pMB1, LacIQ - Ara, Native GroES Seq, sfGFP | This Study |
| pAT00_51_001_sfGFP | pAT00 Base Plasmid construct containing: ColE1/pMB1, LacIQ - Acr, Native GroES Seq, sfGFP | This Study |
| pAT00_52_001_sfGFP | pAT00 Base Plasmid construct containing: ColE1/pMB1, LacI - Oc6, Native GroES Seq, sfGFP | This Study |
| pAT00_20_001_sfGFP_CO | pAT00 Base Plasmid construct containing: ColE1/pMB1, pGroES, Native GroES Seq, Deinoccous codon optimized sfGFP | This Study |
| pAT00_20_001_mCherry | pAT00 Base Plasmid construct containing: ColE1/pMB1, pGroES, Native GroES Seq, mCherry | This Study |
| pAT00_20_001_mCherry_CO | pAT00 Base Plasmid construct containing: ColE1/pMB1, pGroES, Native GroES Seq, Deinoccous codon optimized mCherry | This Study |
| pAT00_20_001_mOrange | pAT00 Base Plasmid construct containing: ColE1/pMB1, pGroES, Native GroES Seq, mOrange | This Study |
| pAT00_20_001_mOrange_CO | pAT00 Base Plasmid construct containing: ColE1/pMB1, pGroES, Native GroES Seq, Deinoccous codon optimized mOrange | This Study |
| pAT00_20_001_mTurq | pAT00 Base Plasmid construct containing: ColE1/pMB1, pGroES, Native GroES Seq, mTurq | This Study |
| pAT00_20_001_mTurq_CO | pAT00 Base Plasmid construct containing: ColE1/pMB1, pGroES, Native GroES Seq, Deinoccous codon optimized mTurq | This Study |
| pAT00_20_002_sfGFP | pAT00 Base Plasmid construct containing: ColE1/pMB1, pGroES, Randomized RBS Seq 2, sfGFP | This Study |
| pAT00_20_003_sfGFP | pAT00 Base Plasmid construct containing: ColE1/pMB1, pGroES, Randomized RBS Seq 3, sfGFP | This Study |
| pAT00_20_004_sfGFP | pAT00 Base Plasmid construct containing: ColE1/pMB1, pGroES, Randomized RBS Seq 4, sfGFP | This Study |
| pAT00_20_005_sfGFP | pAT00 Base Plasmid construct containing: ColE1/pMB1, pGroES, Randomized RBS Seq 5, sfGFP | This Study |
| pAT00_20_006_sfGFP | pAT00 Base Plasmid construct containing: ColE1/pMB1, pGroES, Randomized RBS Seq 6, sfGFP | This Study |
| pAT00_20_007_sfGFP | pAT00 Base Plasmid construct containing: ColE1/pMB1, pGroES, Randomized RBS Seq 7, sfGFP | This Study |
| pAT00_20_008_sfGFP | pAT00 Base Plasmid construct containing: ColE1/pMB1, pGroES, Randomized RBS Seq 8, sfGFP | This Study |
| pAT00_20_009_sfGFP | pAT00 Base Plasmid construct containing: ColE1/pMB1, pGroES, Randomized RBS Seq 9, sfGFP | This Study |
| pAT00_20_010_sfGFP | pAT00 Base Plasmid construct containing: ColE1/pMB1, pGroES, Randomized RBS Seq 10, sfGFP | This Study |
| pAT00_20_011_sfGFP | pAT00 Base Plasmid construct containing: ColE1/pMB1, pGroES, Randomized RBS Seq 11, sfGFP | This Study |
| pAT00_20_012_sfGFP | pAT00 Base Plasmid construct containing: ColE1/pMB1, pGroES, Randomized RBS Seq 12, sfGFP | This Study |
| pAT00_20_013_sfGFP | pAT00 Base Plasmid construct containing: ColE1/pMB1, pGroES, Randomized RBS Seq 13, sfGFP | This Study |
| pAT00_20_014_sfGFP | pAT00 Base Plasmid construct containing: ColE1/pMB1, pGroES, Randomized RBS Seq 14, sfGFP | This Study |
| pAT00_20_015_sfGFP | pAT00 Base Plasmid construct containing: ColE1/pMB1, pGroES, Randomized RBS Seq 15, sfGFP | This Study |
| pAT00_20_016_sfGFP | pAT00 Base Plasmid construct containing: ColE1/pMB1, pGroES, Randomized RBS Seq 16, sfGFP | This Study |
| pAT00_20_017_sfGFP | pAT00 Base Plasmid construct containing: ColE1/pMB1, pGroES, Randomized RBS Seq 17, sfGFP | This Study |
| pAT00_20_018_sfGFP | pAT00 Base Plasmid construct containing: ColE1/pMB1, pGroES, Randomized RBS Seq 18, sfGFP | This Study |
| pAT00_20_019_sfGFP | pAT00 Base Plasmid construct containing: ColE1/pMB1, pGroES, Randomized RBS Seq 19, sfGFP | This Study |
| pAT00_20_020_sfGFP | pAT00 Base Plasmid construct containing: ColE1/pMB1, pGroES, Randomized RBS Seq 20, sfGFP | This Study |
| pAT00_20_021_sfGFP | pAT00 Base Plasmid construct containing: ColE1/pMB1, pGroES, Randomized RBS Seq 21, sfGFP | This Study |
| pAT00_20_022_sfGFP | pAT00 Base Plasmid construct containing: ColE1/pMB1, pGroES, Randomized RBS Seq 22, sfGFP | This Study |
| pAT00_20_023_sfGFP | pAT00 Base Plasmid construct containing: ColE1/pMB1, pGroES, Randomized RBS Seq 23, sfGFP | This Study |
| pAT00_20_024_sfGFP | pAT00 Base Plasmid construct containing: ColE1/pMB1, pGroES, Randomized RBS Seq 24, sfGFP | This Study |
| pAT00_20_025_sfGFP | pAT00 Base Plasmid construct containing: ColE1/pMB1, pGroES, Randomized RBS Seq 25, sfGFP | This Study |
| pAT00_20_026_sfGFP | pAT00 Base Plasmid construct containing: ColE1/pMB1, pGroES, Randomized RBS Seq 26, sfGFP | This Study |
| pAT00_20_027_sfGFP | pAT00 Base Plasmid construct containing: ColE1/pMB1, pGroES, Randomized RBS Seq 27, sfGFP | This Study |
| pAT00_20_028_sfGFP | pAT00 Base Plasmid construct containing: ColE1/pMB1, pGroES, Randomized RBS Seq 28, sfGFP | This Study |
| pAT00_20_029_sfGFP | pAT00 Base Plasmid construct containing: ColE1/pMB1, pGroES, Randomized RBS Seq 29, sfGFP | This Study |
| pAT00_20_030_sfGFP | pAT00 Base Plasmid construct containing: ColE1/pMB1, pGroES, Randomized RBS Seq 30, sfGFP | This Study |
| pAT00_20_031_sfGFP | pAT00 Base Plasmid construct containing: ColE1/pMB1, pGroES, Randomized RBS Seq 31, sfGFP | This Study |
| pAT00_20_032_sfGFP | pAT00 Base Plasmid construct containing: ColE1/pMB1, pGroES, Randomized RBS Seq 32, sfGFP | This Study |
| pAT00_20_033_sfGFP | pAT00 Base Plasmid construct containing: ColE1/pMB1, pGroES, Randomized RBS Seq 33, sfGFP | This Study |
| pAT00_20_034_sfGFP | pAT00 Base Plasmid construct containing: ColE1/pMB1, pGroES, Randomized RBS Seq 34, sfGFP | This Study |
| pAT00_20_035_sfGFP | pAT00 Base Plasmid construct containing: ColE1/pMB1, pGroES, Randomized RBS Seq 35, sfGFP | This Study |
| pAT00_20_036_sfGFP | pAT00 Base Plasmid construct containing: ColE1/pMB1, pGroES, Randomized RBS Seq 36, sfGFP | This Study |
| pAT00_20_037_sfGFP | pAT00 Base Plasmid construct containing: ColE1/pMB1, pGroES, Randomized RBS Seq 37, sfGFP | This Study |
| pAT00_20_038_sfGFP | pAT00 Base Plasmid construct containing: ColE1/pMB1, pGroES, Randomized RBS Seq 38, sfGFP | This Study |
| pAT00_20_039_sfGFP | pAT00 Base Plasmid construct containing: ColE1/pMB1, pGroES, Randomized RBS Seq 39, sfGFP | This Study |
| pAT00_20_040_sfGFP | pAT00 Base Plasmid construct containing: ColE1/pMB1, pGroES, Randomized RBS Seq 40, sfGFP | This Study |
| pAT00_20_041_sfGFP | pAT00 Base Plasmid construct containing: ColE1/pMB1, pGroES, Randomized RBS Seq 41, sfGFP | This Study |
| pAT00_20_042_sfGFP | pAT00 Base Plasmid construct containing: ColE1/pMB1, pGroES, Randomized RBS Seq 42, sfGFP | This Study |
| pAT00_20_043_sfGFP | pAT00 Base Plasmid construct containing: ColE1/pMB1, pGroES, Randomized RBS Seq 43, sfGFP | This Study |
| pAT00_20_044_sfGFP | pAT00 Base Plasmid construct containing: ColE1/pMB1, pGroES, Randomized RBS Seq 44, sfGFP | This Study |
| pAT00_20_045_sfGFP | pAT00 Base Plasmid construct containing: ColE1/pMB1, pGroES, Randomized RBS Seq 45, sfGFP | This Study |
| pAT00_20_046_sfGFP | pAT00 Base Plasmid construct containing: ColE1/pMB1, pGroES, Randomized RBS Seq 46, sfGFP | This Study |
| pAT00_20_047_sfGFP | pAT00 Base Plasmid construct containing: ColE1/pMB1, pGroES, Randomized RBS Seq 47, sfGFP | This Study |
| pAT00_20_048_sfGFP | pAT00 Base Plasmid construct containing: ColE1/pMB1, pGroES, Randomized RBS Seq 48, sfGFP | This Study |
| pAT00_20_049_sfGFP | pAT00 Base Plasmid construct containing: ColE1/pMB1, pGroES, Randomized RBS Seq 49, sfGFP | This Study |
| pAT00_20_050_sfGFP | pAT00 Base Plasmid construct containing: ColE1/pMB1, pGroES, Randomized RBS Seq 50, sfGFP | This Study |
| pAT00_20_051_sfGFP | pAT00 Base Plasmid construct containing: ColE1/pMB1, pGroES, Randomized RBS Seq 51, sfGFP | This Study |
| pAT00_20_052_sfGFP | pAT00 Base Plasmid construct containing: ColE1/pMB1, pGroES, Randomized RBS Seq 52, sfGFP | This Study |
| pAT00_20_053_sfGFP | pAT00 Base Plasmid construct containing: ColE1/pMB1, pGroES, Randomized RBS Seq 53, sfGFP | This Study |
| pAT00_20_054_sfGFP | pAT00 Base Plasmid construct containing: ColE1/pMB1, pGroES, Randomized RBS Seq 54, sfGFP | This Study |
| pAT00_20_055_sfGFP | pAT00 Base Plasmid construct containing: ColE1/pMB1, pGroES, Randomized RBS Seq 55, sfGFP | This Study |
| pAT00_20_056_sfGFP | pAT00 Base Plasmid construct containing: ColE1/pMB1, pGroES, Randomized RBS Seq 56, sfGFP | This Study |
| pAT00_20_057_sfGFP | pAT00 Base Plasmid construct containing: ColE1/pMB1, pGroES, Randomized RBS Seq 57, sfGFP | This Study |
| pAT00_20_058_sfGFP | pAT00 Base Plasmid construct containing: ColE1/pMB1, pGroES, Randomized RBS Seq 58, sfGFP | This Study |
| pAT00_20_059_sfGFP | pAT00 Base Plasmid construct containing: ColE1/pMB1, pGroES, Randomized RBS Seq 59, sfGFP | This Study |
| pAT00_20_060_sfGFP | pAT00 Base Plasmid construct containing: ColE1/pMB1, pGroES, Randomized RBS Seq 60, sfGFP | This Study |
| pAT00_20_061_sfGFP | pAT00 Base Plasmid construct containing: ColE1/pMB1, pGroES, Randomized RBS Seq 61, sfGFP | This Study |
| pAT00_20_062_sfGFP | pAT00 Base Plasmid construct containing: ColE1/pMB1, pGroES, Randomized RBS Seq 62, sfGFP | This Study |
| pAT00_20_063_sfGFP | pAT00 Base Plasmid construct containing: ColE1/pMB1, pGroES, Randomized RBS Seq 63, sfGFP | This Study |
| pAT00_20_064_sfGFP | pAT00 Base Plasmid construct containing: ColE1/pMB1, pGroES, Randomized RBS Seq 64, sfGFP | This Study |
| pAT00_20_065_sfGFP | pAT00 Base Plasmid construct containing: ColE1/pMB1, pGroES, Randomized RBS Seq 65, sfGFP | This Study |
| pAT00_20_066_sfGFP | pAT00 Base Plasmid construct containing: ColE1/pMB1, pGroES, Randomized RBS Seq 66, sfGFP | This Study |
| pAT00_20_067_sfGFP | pAT00 Base Plasmid construct containing: ColE1/pMB1, pGroES, Randomized RBS Seq 67, sfGFP | This Study |
| pAT00_20_068_sfGFP | pAT00 Base Plasmid construct containing: ColE1/pMB1, pGroES, Randomized RBS Seq 68, sfGFP | This Study |
| pAT00_20_069_sfGFP | pAT00 Base Plasmid construct containing: ColE1/pMB1, pGroES, Randomized RBS Seq 69, sfGFP | This Study |
| pAT00_20_070_sfGFP | pAT00 Base Plasmid construct containing: ColE1/pMB1, pGroES, Randomized RBS Seq 70, sfGFP | This Study |
| pAT00_20_071_sfGFP | pAT00 Base Plasmid construct containing: ColE1/pMB1, pGroES, Randomized RBS Seq 71, sfGFP | This Study |
| pAT00_20_072_sfGFP | pAT00 Base Plasmid construct containing: ColE1/pMB1, pGroES, Randomized RBS Seq 72, sfGFP | This Study |
| pAT00_20_073_sfGFP | pAT00 Base Plasmid construct containing: ColE1/pMB1, pGroES, Randomized RBS Seq 73, sfGFP | This Study |
| pAT00_20_074_sfGFP | pAT00 Base Plasmid construct containing: ColE1/pMB1, pGroES, Randomized RBS Seq 74, sfGFP | This Study |
| pAT00_20_075_sfGFP | pAT00 Base Plasmid construct containing: ColE1/pMB1, pGroES, Randomized RBS Seq 75, sfGFP | This Study |
| pAT00_20_076_sfGFP | pAT00 Base Plasmid construct containing: ColE1/pMB1, pGroES, Randomized RBS Seq 76, sfGFP | This Study |
| pAT00_20_077_sfGFP | pAT00 Base Plasmid construct containing: ColE1/pMB1, pGroES, Randomized RBS Seq 77, sfGFP | This Study |
| pAT00_20_078_sfGFP | pAT00 Base Plasmid construct containing: ColE1/pMB1, pGroES, Randomized RBS Seq 78, sfGFP | This Study |
| pAT00_20_079_sfGFP | pAT00 Base Plasmid construct containing: ColE1/pMB1, pGroES, Randomized RBS Seq 79, sfGFP | This Study |
| pAT00_20_080_sfGFP | pAT00 Base Plasmid construct containing: ColE1/pMB1, pGroES, Randomized RBS Seq 80, sfGFP | This Study |
| pAT00_20_081_sfGFP | pAT00 Base Plasmid construct containing: ColE1/pMB1, pGroES, Randomized RBS Seq 81, sfGFP | This Study |
| pAT00_20_082_sfGFP | pAT00 Base Plasmid construct containing: ColE1/pMB1, pGroES, Randomized RBS Seq 82, sfGFP | This Study |
| pAT00_20_083_sfGFP | pAT00 Base Plasmid construct containing: ColE1/pMB1, pGroES, Randomized RBS Seq 83, sfGFP | This Study |
| pAT00_20_084_sfGFP | pAT00 Base Plasmid construct containing: ColE1/pMB1, pGroES, Randomized RBS Seq 84, sfGFP | This Study |
| pAT00_20_085_sfGFP | pAT00 Base Plasmid construct containing: ColE1/pMB1, pGroES, Randomized RBS Seq 85, sfGFP | This Study |
| pAT00_20_086_sfGFP | pAT00 Base Plasmid construct containing: ColE1/pMB1, pGroES, Randomized RBS Seq 86, sfGFP | This Study |
| pAT00_20_087_sfGFP | pAT00 Base Plasmid construct containing: ColE1/pMB1, pGroES, Randomized RBS Seq 87, sfGFP | This Study |
| pAT00_20_088_sfGFP | pAT00 Base Plasmid construct containing: ColE1/pMB1, pGroES, Randomized RBS Seq 88, sfGFP | This Study |
| pAT00_20_089_sfGFP | pAT00 Base Plasmid construct containing: ColE1/pMB1, pGroES, Randomized RBS Seq 89, sfGFP | This Study |
| pAT00_20_090_sfGFP | pAT00 Base Plasmid construct containing: ColE1/pMB1, pGroES, Randomized RBS Seq 90, sfGFP | This Study |
| pAT00_20_091_sfGFP | pAT00 Base Plasmid construct containing: ColE1/pMB1, pGroES, Randomized RBS Seq 91, sfGFP | This Study |
| pAT00_20_092_sfGFP | pAT00 Base Plasmid construct containing: ColE1/pMB1, pGroES, Randomized RBS Seq 92, sfGFP | This Study |
| pAT00_20_093_sfGFP | pAT00 Base Plasmid construct containing: ColE1/pMB1, pGroES, Randomized RBS Seq 93, sfGFP | This Study |
| pAT00_20_094_sfGFP | pAT00 Base Plasmid construct containing: ColE1/pMB1, pGroES, Randomized RBS Seq 94, sfGFP | This Study |
| pAT00_20_095_sfGFP | pAT00 Base Plasmid construct containing: ColE1/pMB1, pGroES, Randomized RBS Seq 95, sfGFP | This Study |
| pAT00_20_096_sfGFP | pAT00 Base Plasmid construct containing: ColE1/pMB1, pGroES, Randomized RBS Seq 96, sfGFP | This Study |
| pAT00_20_097_sfGFP | pAT00 Base Plasmid construct containing: ColE1/pMB1, pGroES, Randomized RBS Seq 97, sfGFP | This Study |
| pAT00_20_098_sfGFP | pAT00 Base Plasmid construct containing: ColE1/pMB1, pGroES, Randomized RBS Seq 98, sfGFP | This Study |
| pAT00_20_099_sfGFP | pAT00 Base Plasmid construct containing: ColE1/pMB1, pGroES, Randomized RBS Seq 99, sfGFP | This Study |
| pAT00_20_100_sfGFP | pAT00 Base Plasmid construct containing: ColE1/pMB1, pGroES, Randomized RBS Seq 100, sfGFP | This Study |
| pAT00_20_101_sfGFP | pAT00 Base Plasmid construct containing: ColE1/pMB1, pGroES, Randomized RBS Seq 101, sfGFP | This Study |
| pAT00_20_102_sfGFP | pAT00 Base Plasmid construct containing: ColE1/pMB1, pGroES, Randomized RBS Seq 102, sfGFP | This Study |
| pAT00_20_103_sfGFP | pAT00 Base Plasmid construct containing: ColE1/pMB1, pGroES, Randomized RBS Seq 103, sfGFP | This Study |
| pAT00_20_104_sfGFP | pAT00 Base Plasmid construct containing: ColE1/pMB1, pGroES, Randomized RBS Seq 104, sfGFP | This Study |
| pAT00_20_105_sfGFP | pAT00 Base Plasmid construct containing: ColE1/pMB1, pGroES, Randomized RBS Seq 105, sfGFP | This Study |
| pAT00_20_106_sfGFP | pAT00 Base Plasmid construct containing: ColE1/pMB1, pGroES, Randomized RBS Seq 106, sfGFP | This Study |
| pAT00_20_107_sfGFP | pAT00 Base Plasmid construct containing: ColE1/pMB1, pGroES, Randomized RBS Seq 107, sfGFP | This Study |
| pAT00_20_108_sfGFP | pAT00 Base Plasmid construct containing: ColE1/pMB1, pGroES, Randomized RBS Seq 108, sfGFP | This Study |
| pAT00_20_109_sfGFP | pAT00 Base Plasmid construct containing: ColE1/pMB1, pGroES, Randomized RBS Seq 109, sfGFP | This Study |
| pAT00_20_110_sfGFP | pAT00 Base Plasmid construct containing: ColE1/pMB1, pGroES, Randomized RBS Seq 110, sfGFP | This Study |
| pAT00_20_111_sfGFP | pAT00 Base Plasmid construct containing: ColE1/pMB1, pGroES, Randomized RBS Seq 111, sfGFP | This Study |
| pAT00_20_112_sfGFP | pAT00 Base Plasmid construct containing: ColE1/pMB1, pGroES, Randomized RBS Seq 112, sfGFP | This Study |
| pAT00_20_113_sfGFP | pAT00 Base Plasmid construct containing: ColE1/pMB1, pGroES, Randomized RBS Seq 113, sfGFP | This Study |
| pAT00_20_114_sfGFP | pAT00 Base Plasmid construct containing: ColE1/pMB1, pGroES, Randomized RBS Seq 114, sfGFP | This Study |
| pAT00_20_115_sfGFP | pAT00 Base Plasmid construct containing: ColE1/pMB1, pGroES, Randomized RBS Seq 115, sfGFP | This Study |
| pAT00_20_116_sfGFP | pAT00 Base Plasmid construct containing: ColE1/pMB1, pGroES, Randomized RBS Seq 116, sfGFP | This Study |
| pAT00_20_117_sfGFP | pAT00 Base Plasmid construct containing: ColE1/pMB1, pGroES, Randomized RBS Seq 117, sfGFP | This Study |
| pAT00_20_118_sfGFP | pAT00 Base Plasmid construct containing: ColE1/pMB1, pGroES, Randomized RBS Seq 118, sfGFP | This Study |
| pAT00_20_119_sfGFP | pAT00 Base Plasmid construct containing: ColE1/pMB1, pGroES, Randomized RBS Seq 119, sfGFP | This Study |
| pAT00_20_120_sfGFP | pAT00 Base Plasmid construct containing: ColE1/pMB1, pGroES, Randomized RBS Seq 120, sfGFP | This Study |
| pAT00_20_121_sfGFP | pAT00 Base Plasmid construct containing: ColE1/pMB1, pGroES, Randomized RBS Seq 121, sfGFP | This Study |
| pAT00_20_122_sfGFP | pAT00 Base Plasmid construct containing: ColE1/pMB1, pGroES, Randomized RBS Seq 122, sfGFP | This Study |
| pAT00_20_123_sfGFP | pAT00 Base Plasmid construct containing: ColE1/pMB1, pGroES, Randomized RBS Seq 123, sfGFP | This Study |
| pAT00_20_124_sfGFP | pAT00 Base Plasmid construct containing: ColE1/pMB1, pGroES, Randomized RBS Seq 124, sfGFP | This Study |
| pAT00_20_125_sfGFP | pAT00 Base Plasmid construct containing: ColE1/pMB1, pGroES, Randomized RBS Seq 125, sfGFP | This Study |
| pAT00_20_126_sfGFP | pAT00 Base Plasmid construct containing: ColE1/pMB1, pGroES, Randomized RBS Seq 126, sfGFP | This Study |
| pAT00_20_127_sfGFP | pAT00 Base Plasmid construct containing: ColE1/pMB1, pGroES, Randomized RBS Seq 127, sfGFP | This Study |
| pAT00_20_128_sfGFP | pAT00 Base Plasmid construct containing: ColE1/pMB1, pGroES, Randomized RBS Seq 128, sfGFP | This Study |
| pAT00_20_129_sfGFP | pAT00 Base Plasmid construct containing: ColE1/pMB1, pGroES, Randomized RBS Seq 129, sfGFP | This Study |
| pAT00_20_130_sfGFP | pAT00 Base Plasmid construct containing: ColE1/pMB1, pGroES, Randomized RBS Seq 130, sfGFP | This Study |
| pAT00_20_131_sfGFP | pAT00 Base Plasmid construct containing: ColE1/pMB1, pGroES, Randomized RBS Seq 131, sfGFP | This Study |
| pAT00_20_132_sfGFP | pAT00 Base Plasmid construct containing: ColE1/pMB1, pGroES, Randomized RBS Seq 132, sfGFP | This Study |
| pAT00_20_133_sfGFP | pAT00 Base Plasmid construct containing: ColE1/pMB1, pGroES, Randomized RBS Seq 133, sfGFP | This Study |
| pAT00_20_134_sfGFP | pAT00 Base Plasmid construct containing: ColE1/pMB1, pGroES, Randomized RBS Seq 134, sfGFP | This Study |
| pAT00_20_135_sfGFP | pAT00 Base Plasmid construct containing: ColE1/pMB1, pGroES, Randomized RBS Seq 135, sfGFP | This Study |
| pAT00_20_136_sfGFP | pAT00 Base Plasmid construct containing: ColE1/pMB1, pGroES, Randomized RBS Seq 136, sfGFP | This Study |
| pAT00_20_137_sfGFP | pAT00 Base Plasmid construct containing: ColE1/pMB1, pGroES, Randomized RBS Seq 137, sfGFP | This Study |
| pAT00_40_001_dTnpB_20_ReRNA_00 | pAT00 Base Plasmid construct containing: ColE1/pMB1, IPTG Inducible Promoter, Native GroES Seq, dTnpB, pGroES, Guide RNA Number 00 | This Study |
| pAT00_45_001_dTnpB_20_ReRNA_00 | pAT00 Base Plasmid construct containing: ColE1/pMB1, aTc Inducible Promoter, Native GroES Seq, dTnpB, pGroES, Guide RNA Number 00 | This Study |
| pAT00_45_001_dTnpB_20_ReRNA_01 | pAT00 Base Plasmid construct containing: ColE1/pMB1, aTc Inducible Promoter, Native GroES Seq, dTnpB, pGroES, Guide RNA Number 01 | This Study |
| pAT00_45_001_dTnpB_20_ReRNA_02 | pAT00 Base Plasmid construct containing: ColE1/pMB1, aTc Inducible Promoter, Native GroES Seq, dTnpB, pGroES, Guide RNA Number 02 | This Study |
| pAT00_45_001_dTnpB_20_ReRNA_03 | pAT00 Base Plasmid construct containing: ColE1/pMB1, aTc Inducible Promoter, Native GroES Seq, dTnpB, pGroES, Guide RNA Number 03 | This Study |
| pAT00_45_001_dTnpB_20_ReRNA_04 | pAT00 Base Plasmid construct containing: ColE1/pMB1, aTc Inducible Promoter, Native GroES Seq, dTnpB, pGroES, Guide RNA Number 04 | This Study |
| pAT00_45_001_dTnpB_20_ReRNA_05 | pAT00 Base Plasmid construct containing: ColE1/pMB1, aTc Inducible Promoter, Native GroES Seq, dTnpB, pGroES, Guide RNA Number 05 | This Study |
| pAT00_45_001_dTnpB_20_ReRNA_06 | pAT00 Base Plasmid construct containing: ColE1/pMB1, aTc Inducible Promoter, Native GroES Seq, dTnpB, pGroES, Guide RNA Number 06 | This Study |
| pAT00_45_001_dTnpB_20_ReRNA_07 | pAT00 Base Plasmid construct containing: ColE1/pMB1, aTc Inducible Promoter, Native GroES Seq, dTnpB, pGroES, Guide RNA Number 07 | This Study |
| pAT00_45_001_dTnpB_20_ReRNA_08 | pAT00 Base Plasmid construct containing: ColE1/pMB1, aTc Inducible Promoter, Native GroES Seq, dTnpB, pGroES, Guide RNA Number 08 | This Study |
| pAT00_45_001_dTnpB_20_ReRNA_09 | pAT00 Base Plasmid construct containing: ColE1/pMB1, aTc Inducible Promoter, Native GroES Seq, dTnpB, pGroES, Guide RNA Number 09 | This Study |
| pAT00_45_001_dTnpB_20_ReRNA_10 | pAT00 Base Plasmid construct containing: ColE1/pMB1, aTc Inducible Promoter, Native GroES Seq, dTnpB, pGroES, Guide RNA Number 10 | This Study |
| pAT00_20_084_sfGFP | Plamid for gene mutation in D. radiodurans containing mPhes cassette for counterselection by 4-cholorphenylalanine | Roland J. Saldanha and Thomas Lamkin |
| pAT00_20_085_sfGFP | pMLKp1261 backbone from pRad-DR2009_3xFLAG, utilizing BamHI & SacII Rest. Digest to insert Candidate Gene from puc19 Genscript plasmid | This Study |
| pAT00_20_086_sfGFP | pMLKp1261 backbone from pRad-DR2009_3xFLAG, utilizing BamHI & SacII Rest. Digest to insert Candidate Gene from puc19 Genscript plasmid | This Study |
| pAT00_20_087_sfGFP | pMLKp1261 backbone from pRad-DR2009_3xFLAG, utilizing BamHI & SacII Rest. Digest to insert Candidate Gene from puc19 Genscript plasmid | This Study |
| pAT00_20_088_sfGFP | pMLKp1261 backbone from pRad-DR2009_3xFLAG, utilizing BamHI & SacII Rest. Digest to insert Candidate Gene from puc19 Genscript plasmid | This Study |
| pAT00_20_089_sfGFP | pMLKp1261 backbone from pRad-DR2009_3xFLAG, utilizing Gibson Asembly t to insert Candidate Gene from puc19 Genscript plasmid | This Study |
| pAT00_20_090_sfGFP | pMLKp1261 backbone from pRad-DR2009_3xFLAG, utilizing Gibson Assembly to insert Candidate Gene from puc19 Genscript plasmid | This Study |
| pAT00_20_091_sfGFP | pAT00 Base Plasmid construct containing: ColE1/pMB1, pGroES, Native GroES Seq, sfGFP | This Study |
| pAT00_20_092_sfGFP | pAT00 Base Plasmid construct containing: pBBR1, pGroES, Native GroES Seq, sfGFP | This Study |
| pAT00_20_093_sfGFP | pAT00 Base Plasmid construct containing: RK2, pGroES, Native GroES Seq, sfGFP | This Study |
| pAT00_20_094_sfGFP | pAT00 Base Plasmid construct containing: pRO1600, pGroES, Native GroES Seq, sfGFP | This Study |
| pAT00_20_095_sfGFP | pAT00 Base Plasmid construct containing: pUB1110, pGroES, Native GroES Seq, sfGFP | This Study |
| pAT00_20_096_sfGFP | pAT00 Base Plasmid construct containing: cisII, pGroES, Native GroES Seq, sfGFP | This Study |
| pAT00_20_097_sfGFP | pAT00 Base Plasmid construct containing: cisMP, pGroES, Native GroES Seq, sfGFP | This Study |
| pAT00_20_098_sfGFP | pAT00 Base Plasmid construct containing: ColE1/pMB1, Anderson 100, Native GroES Seq, sfGFP | This Study |
| pAT00_20_099_sfGFP | pAT00 Base Plasmid construct containing: ColE1/pMB1, Anderson 101, Native GroES Seq, sfGFP | This Study |
| pAT00_20_100_sfGFP | pAT00 Base Plasmid construct containing: ColE1/pMB1, Anderson 102, Native GroES Seq, sfGFP | This Study |
| pAT00_20_101_sfGFP | pAT00 Base Plasmid construct containing: ColE1/pMB1, Anderson 103, Native GroES Seq, sfGFP | This Study |
| pAT00_20_102_sfGFP | pAT00 Base Plasmid construct containing: ColE1/pMB1, Anderson 104, Native GroES Seq, sfGFP | This Study |
| pAT00_20_103_sfGFP | pAT00 Base Plasmid construct containing: ColE1/pMB1, Anderson 105, Native GroES Seq, sfGFP | This Study |
| pAT00_20_104_sfGFP | pAT00 Base Plasmid construct containing: ColE1/pMB1, Anderson 106, Native GroES Seq, sfGFP | This Study |
| pAT00_20_105_sfGFP | pAT00 Base Plasmid construct containing: ColE1/pMB1, Anderson 107, Native GroES Seq, sfGFP | This Study |
| pAT00_20_106_sfGFP | pAT00 Base Plasmid construct containing: ColE1/pMB1, Anderson 108, Native GroES Seq, sfGFP | This Study |
| pAT00_20_107_sfGFP | pAT00 Base Plasmid construct containing: ColE1/pMB1, Anderson 109, Native GroES Seq, sfGFP | This Study |
| pAT00_20_108_sfGFP | pAT00 Base Plasmid construct containing: ColE1/pMB1, Anderson 110, Native GroES Seq, sfGFP | This Study |
| pAT00_20_109_sfGFP | pAT00 Base Plasmid construct containing: ColE1/pMB1, Anderson 111, Native GroES Seq, sfGFP | This Study |
| pAT00_20_110_sfGFP | pAT00 Base Plasmid construct containing: ColE1/pMB1, Anderson 112, Native GroES Seq, sfGFP | This Study |
| pAT00_20_111_sfGFP | pAT00 Base Plasmid construct containing: ColE1/pMB1, Anderson 113, Native GroES Seq, sfGFP | This Study |
| pAT00_20_112_sfGFP | pAT00 Base Plasmid construct containing: ColE1/pMB1, Anderson 114, Native GroES Seq, sfGFP | This Study |
| pAT00_20_113_sfGFP | pAT00 Base Plasmid construct containing: ColE1/pMB1, Anderson 115, Native GroES Seq, sfGFP | This Study |
| pAT00_20_114_sfGFP | pAT00 Base Plasmid construct containing: ColE1/pMB1, Anderson 116, Native GroES Seq, sfGFP | This Study |
| pAT00_20_115_sfGFP | pAT00 Base Plasmid construct containing: ColE1/pMB1, Anderson 117, Native GroES Seq, sfGFP | This Study |
| pAT00_20_116_sfGFP | pAT00 Base Plasmid construct containing: ColE1/pMB1, Anderson 118, Native GroES Seq, sfGFP | This Study |
| pAT00_20_117_sfGFP | pAT00 Base Plasmid construct containing: ColE1/pMB1, Anderson 119, Native GroES Seq, sfGFP | This Study |
| pAT00_20_118_sfGFP | pAT00 Base Plasmid construct containing: ColE1/pMB1, pGroES, Native GroES Seq, sfGFP | This Study |
| pAT00_20_119_sfGFP | pAT00 Base Plasmid construct containing: ColE1/pMB1, p1261, Native GroES Seq, sfGFP | This Study |
| pAT00_20_120_sfGFP | pAT00 Base Plasmid construct containing: ColE1/pMB1, p1348, Native GroES Seq, sfGFP | This Study |
| pAT00_20_121_sfGFP | pAT00 Base Plasmid construct containing: ColE1/pMB1, p1473, Native GroES Seq, sfGFP | This Study |
| pAT00_20_122_sfGFP | pAT00 Base Plasmid construct containing: ColE1/pMB1, p2508, Native GroES Seq, sfGFP | This Study |
| pAT00_20_123_sfGFP | pAT00 Base Plasmid construct containing: ColE1/pMB1, pAmyE, Native GroES Seq, sfGFP | This Study |
| pAT00_20_124_sfGFP | pAT00 Base Plasmid construct containing: ColE1/pMB1, pClpB, Native GroES Seq, sfGFP | This Study |
| pAT00_20_125_sfGFP | pAT00 Base Plasmid construct containing: ColE1/pMB1, pDnaK, Native GroES Seq, sfGFP | This Study |
| pAT00_20_126_sfGFP | pAT00 Base Plasmid construct containing: ColE1/pMB1, pGroEL, Native GroES Seq, sfGFP | This Study |
| pAT00_20_127_sfGFP | pAT00 Base Plasmid construct containing: ColE1/pMB1, pKatA, Native GroES Seq, sfGFP | This Study |
| pAT00_20_128_sfGFP | pAT00 Base Plasmid construct containing: ColE1/pMB1, pLexA, Native GroES Seq, sfGFP | This Study |
| pAT00_20_129_sfGFP | pAT00 Base Plasmid construct containing: ColE1/pMB1, pRecA, Native GroES Seq, sfGFP | This Study |
| pAT00_20_130_sfGFP | pAT00 Base Plasmid construct containing: ColE1/pMB1, pRecQ, Native GroES Seq, sfGFP | This Study |
| pAT00_20_131_sfGFP | pAT00 Base Plasmid construct containing: ColE1/pMB1, pRplL, Native GroES Seq, sfGFP | This Study |
| pAT00_20_132_sfGFP | pAT00 Base Plasmid construct containing: ColE1/pMB1, pRpmB, Native GroES Seq, sfGFP | This Study |
| pAT00_20_133_sfGFP | pAT00 Base Plasmid construct containing: ColE1/pMB1, pTufB, Native GroES Seq, sfGFP | This Study |
| pAT00_20_134_sfGFP | pAT00 Base Plasmid construct containing: ColE1/pMB1, LacI - IPTG, Native GroES Seq, sfGFP | This Study |
| pAT00_20_135_sfGFP | pAT00 Base Plasmid construct containing: ColE1/pMB1, LacIQ - DHBA, Native GroES Seq, sfGFP | This Study |
| pAT00_20_136_sfGFP | pAT00 Base Plasmid construct containing: ColE1/pMB1, LacI - OHC14, Native GroES Seq, sfGFP | This Study |
| pAT00_20_137_sfGFP | pAT00 Base Plasmid construct containing: ColE1/pMB1, LacIQ - Sal, Native GroES Seq, sfGFP | This Study |
| pAT00_40_001_dTnpB_20_ReRNA_00 | pAT00 Base Plasmid construct containing: ColE1/pMB1, LacIQ - CA, Native GroES Seq, sfGFP | This Study |
| pAT00_45_001_dTnpB_20_ReRNA_00 | pAT00 Base Plasmid construct containing: ColE1/pMB1, LacI - aTc, Native GroES Seq, sfGFP | This Study |
| pAT00_45_001_dTnpB_20_ReRNA_01 | pAT00 Base Plasmid construct containing: ColE1/pMB1, LacIQ - Cho, Native GroES Seq, sfGFP | This Study |
| pAT00_45_001_dTnpB_20_ReRNA_02 | pAT00 Base Plasmid construct containing: ColE1/pMB1, LacIQ - Ery, Native GroES Seq, sfGFP | This Study |
| pAT00_45_001_dTnpB_20_ReRNA_03 | pAT00 Base Plasmid construct containing: ColE1/pMB1, LacIQ - Nar, Native GroES Seq, sfGFP | This Study |
| pAT00_45_001_dTnpB_20_ReRNA_04 | pAT00 Base Plasmid construct containing: ColE1/pMB1, LacIQ - Va, Native GroES Seq, sfGFP | This Study |
| pAT00_45_001_dTnpB_20_ReRNA_05 | pAT00 Base Plasmid construct containing: ColE1/pMB1, LacIQ - Ara, Native GroES Seq, sfGFP | This Study |
| pAT00_45_001_dTnpB_20_ReRNA_06 | pAT00 Base Plasmid construct containing: ColE1/pMB1, LacIQ - Acr, Native GroES Seq, sfGFP | This Study |
| pAT00_45_001_dTnpB_20_ReRNA_07 | pAT00 Base Plasmid construct containing: ColE1/pMB1, LacI - Oc6, Native GroES Seq, sfGFP | This Study |
| pAT00_45_001_dTnpB_20_ReRNA_08 | pAT00 Base Plasmid construct containing: ColE1/pMB1, pGroES, Native GroES Seq, Deinoccous codon optimized sfGFP | This Study |
| pAT00_45_001_dTnpB_20_ReRNA_09 | pAT00 Base Plasmid construct containing: ColE1/pMB1, pGroES, Native GroES Seq, mCherry | This Study |
| pAT00_45_001_dTnpB_20_ReRNA_10 | pAT00 Base Plasmid construct containing: ColE1/pMB1, pGroES, Native GroES Seq, Deinoccous codon optimized mCherry | This Study |

**Supplementary Table S7.** Oligonucleotides used in this study

| **Name** | **Sequence (5'-3')** | **Description** |
| --- | --- | --- |
| oAT001 | agctaccattcttgcagc | FW Primer to amplify pRAD1 Backbone towards d rad ori |
| oAT002 | taacgcaggaaagaacatg | RV Primer to amplify pRAD1 towards ColE1 ori |
| oAT003 | aatgaaataagatcactaccgggcgtattttttgagttatcgagattttcaggagctaag | FW Gibson Primer to amplify pRAD1 from cat promoter, includes full cat promoter in homology arm |
| oAT004 | gagtaaacttggtctgacagattcacagttctccgcaa | RV Gibson primer to amplify pRAD1 downstream of CamR cassette, has homology to seq directly upstream of ColE1 ori |
| oAT005 | tcttgcggagaactgtgaatctgtcagaccaagtttactc | FW promoter to amplify pRAD1 upstream of ColE1 ori, homology to oAT004 |
| oAT006 | agtgatcttatttcattatggtgaaagttggaacctcttacgttcgaagctcggcggatt | RV Gibson Primer to amplify pRAD1 towards d rad ori, includes full cat promoter in homology arm |
| oAT007 | tgggacatttgcagacaggacctgctctacgagttgctg | FW Gibson Primer to amplify pAT00 upstream of CamR cassette to remove d rad ori |
| oAT008 | tcctgtctgcaaatgtccca | FW Gibson Primer to amplify pAT00 downstream of rrnBT12 term to remove d rad ori |
| oAT009 | ggaaatgtgctgtcagacca | FW Primer to amplify ColE1 ori thru rrnBT12 term on pAT01 |
| oAT010 | tggtctgacagcacatttcc | Rev primer to amplify CamR cassette to rrnBT12 on pAT01 |
| oAT011 | catgttctttcctgcgttatcc | FW Primer to amplify entire pAT backbone except ori |
| oAT012 | gctgaccctgaagttcatctg | FW Primer for copy number RT-qPCR, lies inside sfgfp CDS |
| oAT013 | gcttgtcggcggtgatatag | RV Primer for copy number RT-qPCR, lies inside sfgfp CDS |
| oAT014 | CGCGCCATCGAATGGTG | FW Primer to amplify PlacIQ ONLY marionette regulator regions |
| oAT015 | GTGAGCGAGGAAGCACCT | RV Primer to amplify marionette regulators |
| oAT016 | CGCGCCATCGAATGGCG | FW Primer to amplify PlacI WT ONLY marionette regulator regions |
| oAT017 | TGAGGTGCTTCCTCGCTCACtcagcaaactgagaaccc | FW Gibson Primer to amplify pAT00 backbone with homology to marionette regulator regions |
| oAT018 | CCATTCGATGGCGCGCCGCtagatacaaagaacacgtcaag | RV Gibson Primer to amplify pAT00 backbone with homology to marionette regulator regions |
| oAT019 | CAACCGCACTTGATTTAATAGACCATACCGTCTATTATTTCTGGCCATcggattgaaggaggaccc | FW Gibson Primer to insert Pacu promoter |
| oAT020 | TATTAAATCAAGTGCGGTTGTGAAGCGTCGGCCTTACTTGCTAGCGttatccacagaatcagggg | RV Gibson Primer to insert Pacu promoter |
| oAT021 | cggattgaaggaggacc | FW Primer binding at RBS01 on pAT00-based plasmids, for cloning inducible promoter seqs |
| oAT022 | TATCAGTGATAGAGATTGACATCCCTATCAGTGATAGAGATACTGAGCACcggattgaaggaggaccc | FW Primer to insert PTet_star promoter into pAT00 plasmids |
| oAT023 | GTCAATCTCTATCACTGATAGGGAGGTCAGTGCGTCCTGCTGAAAAttatccacagaatcagggg | RV Primer to insert PTet_star promoter into pAT00 plasmids |
| oAT024 | gaagttcatctgcaccacc | FW Primer to insert PCymRC Promoter into pAT00 |
| oAT025 | TACAAACAACCATGAATGTAAGTATATTCCTTAGCAAcggattgaaggaggaccc | FW Gibson Primer to insert Ptgt promoter into pAT00 |
| oAT026 | CTTACATTCATGGTTGTTTGTAAATACTGCTGGGTGttatccacagaatcagggg | RV Gibson Primer to insert Ptgt promoter into pAT00 |
| oAT027 | GGTTTACGCAAGAAAATGGTTTGTTACAGTCGAATAAAcggattgaaggaggacc | FW Gibson Primer to insert PLuxB Promoter into pAT00 |
| oAT028 | ACCATTTTCTTGCGTAAACCTGTACGATCCTACAGGTttatccacagaatcagggg | RV Gibson Primer to insert PLuxB Promoter into pAT00 |
| oAT029 | TGACAGCTAGCTCAGTCCTAGGTACCATTGGATCCAATcggattgaaggaggacc | FW Gibson Primer to insert PVan promoter into pAT00 |
| oAT030 | CCTAGGACTGAGCTAGCTGTCAATTGGATCCAATttatccacagaatcagggg | RV Gibson Primer to insert PVan promoter into pAT00 |
| oAT031 | GTTACCAATAGACAATTGATTGGACGTTCAATATAATGCTAGCcggattgaaggaggaccc | FW Gibson Primer to insert PBetI promoter into pAT00 |
| oAT032 | CCAATCAATTGTCTATTGGTAACGAATCCCTCTCACCCGCGCTttatccacagaatcagggg | RV Gibson Primer to insert PBetI promoter into pAT00 |
| oAT033 | AACCGACGTGACTGTTACATTTAGGTGGCTAAACCCGTCAAcggattgaaggaggaccc | FW Gibson Primer to insert Pmph promoter into pAT00 |
| oAT034 | GCCACCTAAATGTAACAGTCACGTCGGTTATATTCAATCCttatccacagaatcagggg | RV Gibson Primer to insert Pmph promoter into pAT00 |
| oAT035 | TCGTATAATGTGTGGAATTGTGAGCGCTCACAATTcggattgaaggaggaccc | FW Gibson Primer to insert Ptac promoter into pAT00 |
| oAT036 | TCACAATTCCACACATTATACGAGCCGATGATTAATTGTCAACAttatccacagaatcagggg | RV Gibson Primer to insert Ptac promoter into pAT00 |
| oAT037 | TGGACGAGCTGTACAAGTGActgttttggcggatgagaga | FW gibson primer to amplify pAT00 at rrnbt12 term |
| oAT038 | TCCTCGCCCTTGCTCACCATgtggggtcctccttcaatc | RV gibson primer to amplify pAT00 at RBS01 |
| oAT039 | cgttagggtatgtctttagtGGGCCCGGTATAATGCCGGG | FW Gibson primer to insert MY1 ori part 1 for RepU in pAT00 |
| oAT040 | aagttggaacctcttacgtCCCGGGACCAGCTCCCCC | RV Gibson primer to insert MY1 part 2 ori for RepU in pAT00 |
| oAT041 | acgtaagaggttccaactt | FW Primer to amplify pAT00 backbone at cat promoter |
| oAT042 | acgtaagaggttccaactt | RV Primer to amplify pAToo backbone downstream of rrnB1 term |
| oAT043 | CTGGCCGTGGCCCTGGGAA | FW Primer to amplify MY1 ori part 2 (use with oAT40) |
| oAT044 | TTCCCAGGGCCACGGCCAG | RV Primer to amplify MY1 ori part 1 (use with oAT39) |
| oAT045 | gacggtcgtcttcatggtcc | FW Primer to amplify TsYS45 ori part 2 |
| oAT046 | ggaccatgaagacgaccgtc | RV Primer to amplify TsYS45 ori part 1 |
| oAT047 | aagttggaacctcttacgtcacgcactccaaccaaggaa | RV Gibson primer to insert TsYS45 part 2 ori for RepU in pAT00 |
| oAT048 | cgttagggtatgtctttagtgctagccgtatgcattagtgaacc | FW Gibson primer to insert TsYS45 ori part 1 for RepU in pAT00 |
| oAT049 | TTGGAAAGTCTTTGCCAGTT | FW Primer to amplify IP501 ori part 2 |
| oAT050 | AACTGGCAAAGACTTTCCAA | RV Primer to amplify IP501 ori part 1 |
| oAT051 | cgttagggtatgtctttagtAACTAACTCAACGCTAGTAGTG | FW Gibson primer to insert IP501 ori part 1 for RepU in pAT00 |
| oAT052 | aagttggaacctcttacgtCCAACTAAAAGTTTTCGGGC | RV Gibson primer to insert IP501 part 2 ori for RepU in pAT00 |
| oAT053 | acgcacaagtgtctcacgtcg | FW Primer to amplify pAT00 plasmid with homology to insert TnpB reRNA seq |
| oAT054 | cacgcgactttagtcgtgtgactgttttggcggatgagaga | FW Gibson Primer to insert TnpB into pAT00_40_001 plasmid (IPTG-inducible) |
| oAT055 | gaaagccttattccttatcatgtggggtcctccttcaatc | RV Gibson Primer to insert TnpB into pAT00_40_001 plasmid (IPTG-inducible) |
| oAT056 | gatagtgttcacccttgttaca | FW Primer to amplify c-terminus of CmR on pAT00, to use with inducible promoter primers |
| oAT057 | tgtaacaagggtgaacactatc | RV Primer to amplify n-terminus of CmR on pAT, to use with inducible promoters |
| oAT058 | acaagtgtctcacgtcg | FW Primer to amplify pAT00.40.001 backbone to insert reRNA (re-attempt) |
| oAT059 | taagaaaaatgtcaactgacttggactaaagacataccctaac | RV Gibson primer to amplify pAT00.40.001 backbone to insert reRNA |
| oAT060 | TCAAgttgctatcacgccgggatctgacagctagcataacccc | FW Gibson primer to remove HDV from reRNA |
| oAT061 | gttatgctagctgtcagatcccggcgtgatagcaacTTGA | RV Gibson primer to remove HDV from reRNA |
| oAT062 | TGAATCACTTCGTGGGggccggcatggtcccag | FW Gibson primer to insert reRNA2-mcheery |
| oAT063 | CCCACGAAGTGATTCATTGAACCTCACACGACTAAAGTCGC | RV Gibson primer to insert reRNA2-mcheery |
| oAT064 | CCCACGAAGTGATTCATTGAACCTCACACGACTAAAGTCGC | FW Gibson primer to insert reRNA3-mcheery |
| oAT065 | GCTTTGCCCAGATTGATTGAACCTCACACGACTAAAGTCGC | RV Gibson primer to insert reRNA3-mcheery |
| oAT066 | cgttagggtatgtctttagtACGTCTACCGCTAGGGAC | FW Gibson Primer to insert TT8 ori into pAT00 |
| oAT067 | aaagttggaacctcttacgtCCTGGGGAGAGAGGCGG | RV Gibson Primer to insert TT8 ori into pAT00 |
| oAT068 | cggctcgtataatgtgtggaattgtgagcggataacaacggattgaaggaggacc | FW Gibson Primer to replace Plac with Ptrc in pAT00.40 plasmids |
| oAT069 | acaattccacacattatacgagccggatgattaattgtcaattatccacagaat | RV Gibson Primer to replace Plac with Ptrc in pAT00.40 plasmids |
| oAT070 | cgtataatgtgtggaattgtgagcggataacaacggattgaaggaggacc | FW Gibson Primer to replace Plac with PlacUV5 |
| oAT071 | ccgctcacaattccacacattatacgagccggaagcttatccacagaatcagggg | RV Gibson Primer to replace Plac with PlacUV5 |
| oAT072 | cgcagtcatgacacaact | RV Primer to truncate dTnpB protein |
| oAT073 | ccagttgtgtcatgactgcgtgactgttttggcggatg | FW Gibson Primer to truncate dTnpB protein |
| oAT074 | AGGACCCCACtTGGTGAGCAA | Fw Primer to Change start codon in pAT00 to TTG |
| oAT075 | CCTTCAATCCGTCACGG | Rv Primer to Change start codon in pAT00 to TTG |
| oAT076 | AGGACCCCACgTGGTGAGCAA | Fw Primer to Change start codon in pAT00 to GTG |
| oAT077 | CCTTCAATCCGTCACGGATG | Rv Primer to Change start codon in pAT00 to GTG |
| oAT078 | GAAGGCCTTGTTGCGGATCATgtggggtcctccttcaatc | RV Primer to insert Codon Optimized TnpB (D191A) |
| oAT079 | gaagacgttgttgacttcctcgacagtc | Forward qPCR primer for control GAPDH |
| oAT080 | gcgatcaacatcattcccacctcg | Reverse qPCR primer for control GAPDH |
| oAT081 | GCCATTCGATGGCGCGCCGCtagatacaaagaacacgtcaagaccc | Gibson Primer to edit pAT00.40.001-TnpB-ReRNA by Swapping IPTG and ATC Promoter/Regulators |
| oAT082 | cttgacgtgttctttgtatctaGCGGCGCGCCATCGA | Gibson Primer to edit pAT00.40.001-TnpB-ReRNA by Swapping IPTG and ATC Promoter/Regulators |
| oAT083 | gagtgagcgaggaagcacctCAGATAAAATATTTGCTCATGAGCCCGA | Gibson Primer to edit pAT00.40.001-TnpB-ReRNA by Swapping IPTG and ATC Promoter/Regulators |
| oAT084 | ATGAGCAAATATTTTATCTGaggtgcttcctcgct | Gibson Primer to edit pAT00.40.001-TnpB-ReRNA by Swapping IPTG and ATC Promoter/Regulators |
| oAT085 | GGTATGGCATGATAGCGCCCGGAAGAGAGTCAATTCAGGGT | Gibson Primer to edit pAT00.40.001-TnpB-ReRNA by Swapping IPTG and ATC Promoter/Regulators |
| oAT086 | CCCTGAATTGACTCTCTTCCGGGCGCTATCATGCCAT | Gibson Primer to edit pAT00.40.001-TnpB-ReRNA by Swapping IPTG and ATC Promoter/Regulators |
| kTO033 | gtggggtcctccttcaatccggccctcaccgaaagg | remove the predicted repU promoter. Can be paired with oAT057 |
| kTO034 | cggattgaaggaggaccccacttgagacacaatccattcatg | remove the predicted repU promoter. Can be paired with oAT056 |
| kTO035 | gctagcattatacctaggactgagctagctgtcaagccctcaccgaaagg | Add J23119 upstream of RBS01 in RepU D. rad Origin. Can be used with oAT057 |
| kTO036 | ttgacagctagctcagtcctaggtataatgctagccggattgaaggaggacc | Add J23119 upstream of RBS01 in RepU D. rad Origin. Can be used with oAT056 |
| kTO041 | ATaagttgcgattatagactgatggctagctcagtcc | Insert J3 Modified into pAT00-13-01-sfgfp, pair with oAT057 |
| kTO042 | gatATCAAcgcaaatgctttatccacagaatcagggg | Insert J3 Modified into pAT00-xx-01-sfgfp, pair with oAT056 |
| kTO043 | GATaagttgcgattatagattgacagctagctcagtc | Insert J3 Modified into pAT00-19-01-sfgfp, pair with oAT057 |
| kTO094 | actaaagacataccctaacgg | Amplify BB pAT00 to insert all new D. rad genetic origins |
| kTO095 | acgtaagaggttccaactt | Amplify BB pAT00 to insert all new D. rad genetic origins |
| kTO096 | ccgttagggtatgtctttagtgatcaaatcttttaaagataaaaacaaaaaaat | FW Primer to Copy cisII Ori gene from R1 genome |
| kTO097 | aagttggaacctcttacgtcttaatagacctgtaattgtctagac | RV Primer to Copy cisII Ori gene from R1 genome |
| kTO098 | ccgttagggtatgtctttagtcaaggacggcttctctc | FW Primer to copy cisMP Ori gene from R1 genome |
| kTO099 | aagttggaacctcttacgtagttgcagaccatagggg | RV Primer to copy cisMP Ori gene from R1 genome |
| kTO100 | ccgttagggtatgtctttagtCGGCCATTCTACGCC | FW Primer to copy oriC Ori gene from R1 genome |
| kTO101 | aagttggaacctcttacgtGGTTGCCGGGCCA | RV Primer to copy oriC Ori gene from R1 genome |
| AC496 | ctgcaggtcgaatcggatccTCACTTGTACAGCTCGTCCA | RV Gibson Primer for all C-Candidate Xenotext Genes |
| AC497 | cttccatgctgaaaccgcggATGGTCTCGAAAGGGGAGGA | FW Gibson Primer for all N-Candidate Xenotext Genes |
| AC498 | cttccatgctgaaaccgcggATGCATCACCATCACCATCACGTGA | FW Gibson Primer for 1C-Candidate Xenotext Gene |
| AC499 | ctgcaggtcgaatcggatccTCAGTGATGGTGATGGTGATG | RV Gibson Primer for 1N-Candidate Xenotext Gene |
| AC500 | cttccatgctgaaaccgcggATGCATCACCATCACCATCACGTA | FW Gibson Primer for 2C-Candidate Xenotext Gene |
| AC501 | ctgcaggtcgaatcggatccTCAGTGATGGTGATGGTGATGA | RV Gibson Primer for 2N-Candidate Xenotext Gene |
| AC502 | cttccatgctgaaaccgcggATGCATCACCATCACCATCAC | FW Gibson Primer for 3C-Candidate Xenotext Gene |
| AC503 | ctgcaggtcgaatcggatccTCAGTGATGGTGATGGTGATGG | RV Gibson Primer for 3N-Candidate Xenotext Gene |
| AC511 | agtcgacctgcaggcatgcaGGTCAAATATCTGTTCACCCTCA | Primer to Perform Genome Integration for Xenotext Candidates into DRAD |
| AC512 | cgtggctcccagggcaggtcCCCACGAAGTGATTCAATCT | Primer to Perform Genome Integration for Xenotext Candidates into DRAD |
| AC513 | agattgaatcacttcgtgggGACCTGCCCTGGGAGCCA | Primer to Perform Genome Integration for Xenotext Candidates into DRAD |
| AC514 | gtatgctatacgaacggtatCACTTGTACAGCTCGTCCATG | Primer to Perform Genome Integration for Xenotext Candidates into DRAD |
| AC515 | tggacgagctgtacaagtgaTACCGTTCGTATAGCATAC | Primer to Perform Genome Integration for Xenotext Candidates into DRAD |
| AC516 | gcccaactccgattgaatcgTACCGTTCGTATAATGTATGC | Primer to Perform Genome Integration for Xenotext Candidates into DRAD |
| AC517 | catacattatacgaacggtaCGATTCAATCGGAGTTGGGCCCA | Primer to Perform Genome Integration for Xenotext Candidates into DRAD |
| AC518 | ctatgaccatgattacgccaTTTCCGGGTCGGCAGCG | Primer to Perform Genome Integration for Xenotext Candidates into DRAD |
| AC519 | TGGACGAGCTGTACAAGTGATACCGTTCGTATAGCATAC | Primer to Perform Colony PCR to confirm Genome Integration into DRAD |
| AC520 | gcccaactccgattgaatcgTACCGTTCGTATAATGTATGC | Primer to Perform Colony PCR to confirm Genome Integration into DRAD |

**Supplementary Figures**

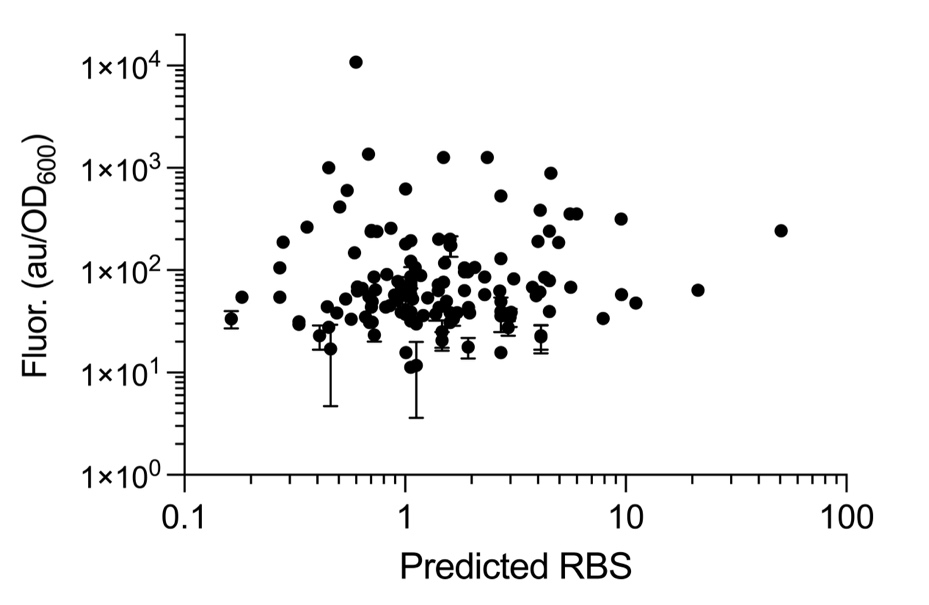

**Supplementary Figure S1. Computationally Predicted RBS strengths vs Relative fluorescence of each RBS design expressing sfGFP.**

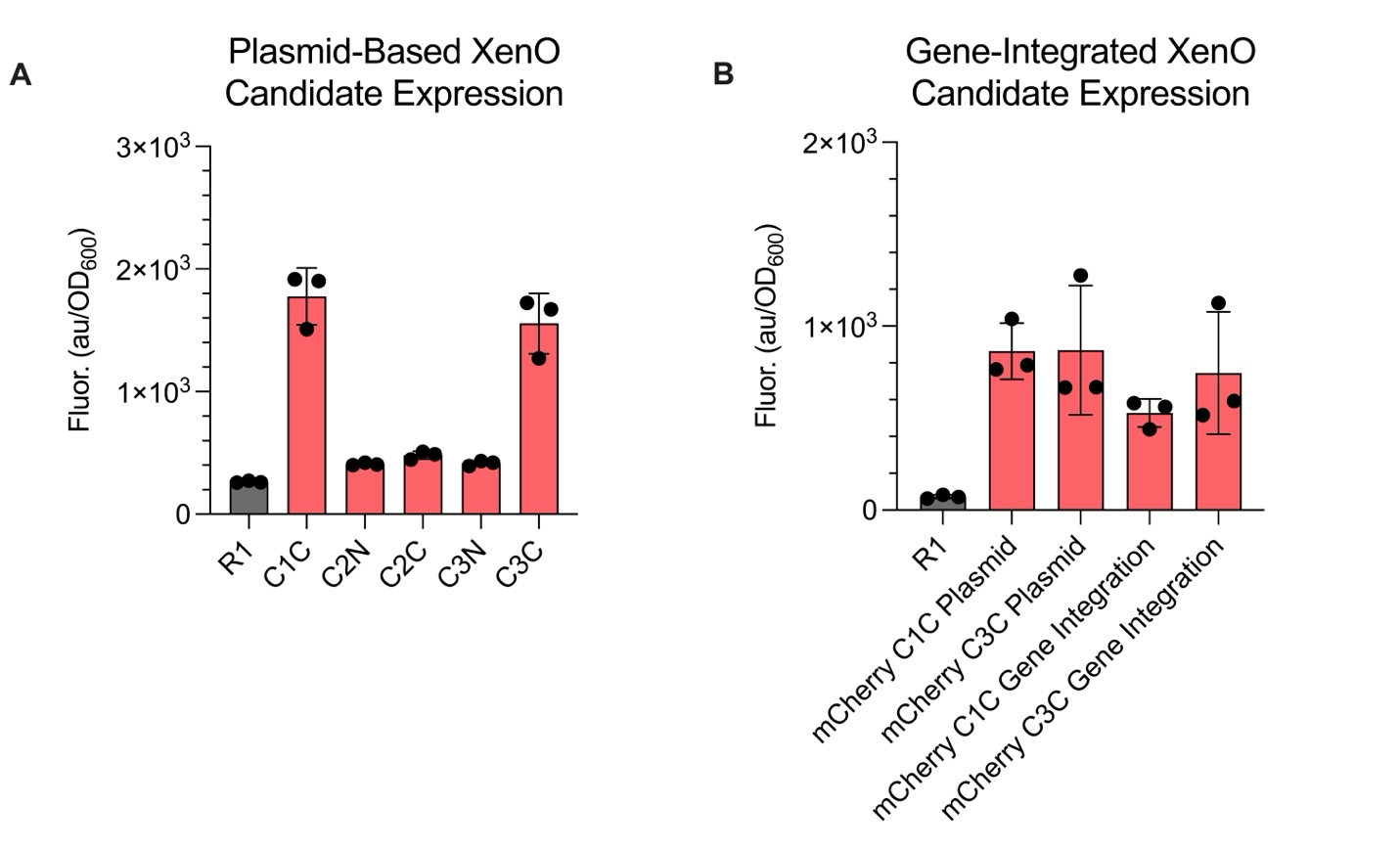
**Supplementary Figure S2. Determining XenO-mCherry Fusion Candidates.** A) Preliminary XenO candidates (C1-3) with mCherry fused to the N-terminus (N) or C-terminus (C) of XenO. B) Plasmid and Genomic fluorescence of the XenO Candidate 1-mCherry and XenO Candidate 3-mCherry fusions.

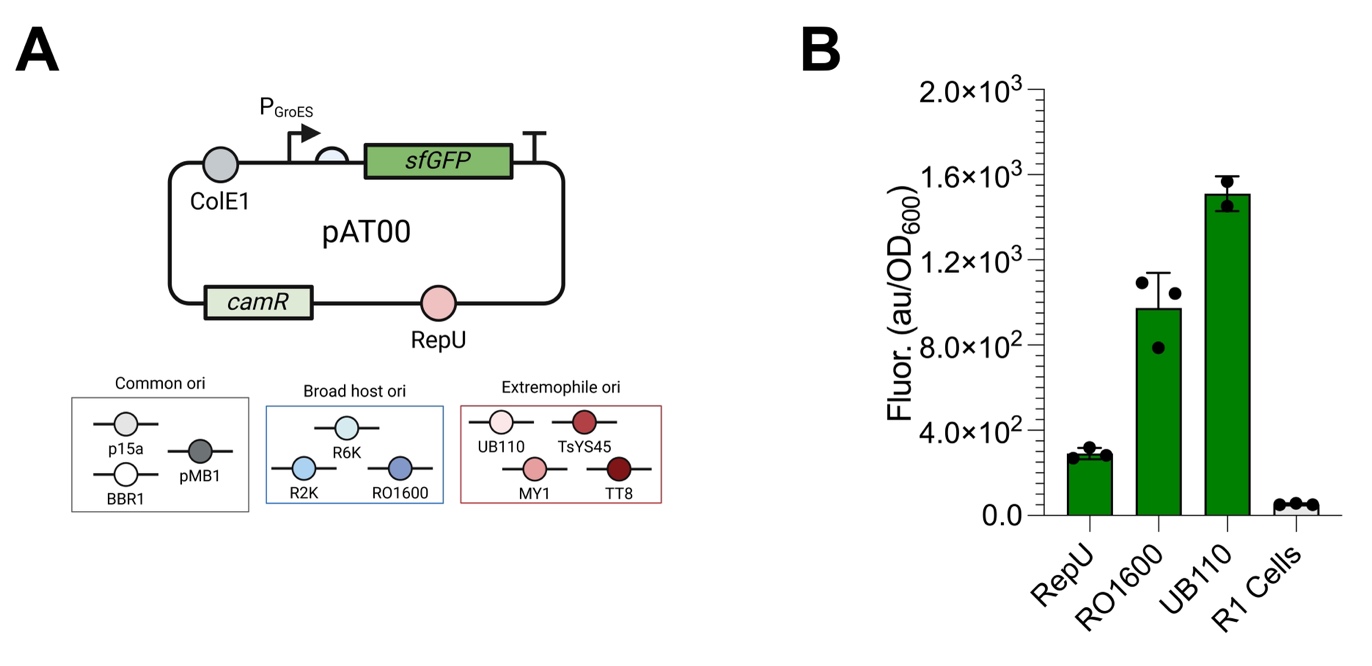

**Supplementary Figure S3. Designing a minimal base plasmid for expression in *D. radiodurans* and identifying novel origin of replications to expand plasmid versatility.** A) Base plasmid for standardization, pAT00, which contains *sfGFP* under P_GroES_ expression, the characterized GroES RBS sequence, the ColE1 origin of replication for sub-cloning in *E. coli*, a CamR cassette, and the RepU ori native to *D. radiodurans*. To evaluate ori compatibility in *D. rad.* the RepU sequence was swapped with oris from three main categories: Common oris, Broad host oris, and extremophile oris. B) sfGFP signal after 24 hours of culturing for each pAT plasmid containing a different ori. Oris without fluorescence values correspond to those that did not form colonies when transformed into *D. radiodurans.*
